# RapidGLM: genome-scale differential expression, CRISPR screen and eQTL analysis

**DOI:** 10.64898/2026.11.18.767389

**Authors:** Ling-Hong Hung, Ka Yee Yeung

## Abstract

Genome-scale differential expression, CRISPR screen and expression QTL analyses fit millions of count regressions and estimate *p*-values from billions of random draws. RapidGLM, a C++ library, makes both steps fast, with Newton fits specialized to count models and draws made in batches. Its DESeq2, edgeR, SCEPTRE and cis-QTL analyses agree with the reference tools and run 2.5–17.8 times faster on the same cores. On one compute node, a CRISPR screen’s 28.0 million tests took SCEPTRE 11.6 hours and RapidGLM 44 minutes, or 2.8 hours with more accurate small *p*-values. RapidGLM is MIT-licensed, with C, Python, R and command-line interfaces.

## Background

Differential expression, CRISPR screen analysis and expression quantitative trait locus (eQTL) mapping are now routinely run at genome scale, with thousands of treatments, perturbations or variants per gene. Tools that are fast enough for small assays do not scale to genome-wide analyses such as that of the Gasperini screen [1], where testing each of 3,776 targets against each of 7,418 genes takes 28.0 million regression tests. The *p*-value of each of these tests is estimated by simulation, from hundreds of random draws or more, billions in all. In the Replogle K562 screen [2], each of the 2,057 perturbations needs its own complete DESeq2 [3] or edgeR [4, 5] analysis, of up to 8,563 genes. In a cis-eQTL scan of Geuvadis [6], each of 20,427 genes is tested against the variants near it,6.3 million variants in all, and the *p*-value of each gene’s strongest association is estimated from 10,000 permutations.

Each individual test asks whether a treatment changes a gene’s expression and whether the observed effect is statistically significant. In differential expression, the treatment is a condition, such as a drug or a genetic perturbation, applied to some samples and not to the controls. In a CRISPR screen, it is the perturbation of a target, such as a gene or an enhancer, by the guides against it, and the controls are cells without those guides. In eQTL mapping, it is the alternative allele of a variant, of which each sample carries 0, 1 or 2 copies. Expression also depends on covariates, such as batch and sequencing depth, so each test compares a regression of the gene’s counts on the treatment and the covariates with the null model, in which the treatment has no effect. Two steps make these calculations expensive. The first is fitting the counts to the models. Each fit is iterative, and a test may need the model with the treatment as well as the null model. Tests that use different samples or cells need separate fits. The second is resampling. In screens and eQTL scans, the *p*-values cannot be calculated reliably from the fitted model alone, so they are estimated by Monte Carlo simulation, known as resampling. The fitted null model is kept, and the treatment is assigned to the cells or samples at random, hundreds of times or more; each random assignment is called a draw. In a screen at high multiplicity of infection (MOI), where each cell carries many guides, a draw marks each cell as perturbed or not, with the probability predicted from its covariates. At low MOI and in eQTL mapping, a draw shuffles the treatment among the cells or samples, a permutation. Either way, the simulated treatment carries no information about the gene’s counts beyond the covariates, so the statistics recomputed for the draws, against the same null fit, show how the statistic varies when the treatment has no effect on the gene’s expression. The *p*-value is estimated as the fraction of draws whose statistic is at least as extreme as the observed one. At genome scale, this requires millions of fits, and billions of random draws for the *p*-values from which the false discovery rate is controlled. These are the steps that must scale, and the objective of this paper is to describe methods that make them fast.

The regression is a generalized linear model (GLM) [7], which relates the distribution of a gene’s counts to the treatment and the covariates (Fig. 1a). For counts, the usual distribution is the negative binomial (NB), whose variance adds the variation between biological replicates to the Poisson variance of counting reads. The model works on the log scale: the logarithm of each sample’s or cell’s expected count is the sum of a baseline, the effect of the treatment, the effects of the covariates and a term for sequencing depth. The treatment’s coefficient is therefore the log fold change, and the treatment has no effect when it is zero. The coefficients are estimated by maximum likelihood, which has no closed form; the fit proceeds by Newton-type steps [7]. Three statistics test whether the treatment has an effect (Fig. 1b). The Wald statistic divides the estimated log fold change by its standard error. The likelihood-ratio statistic measures how much better the model with the treatment fits the counts than the null model. The score statistic measures how steeply the likelihood changes as the treatment’s effect moves away from zero, starting from the null fit, so it needs only that fit.

**Figure 1:**
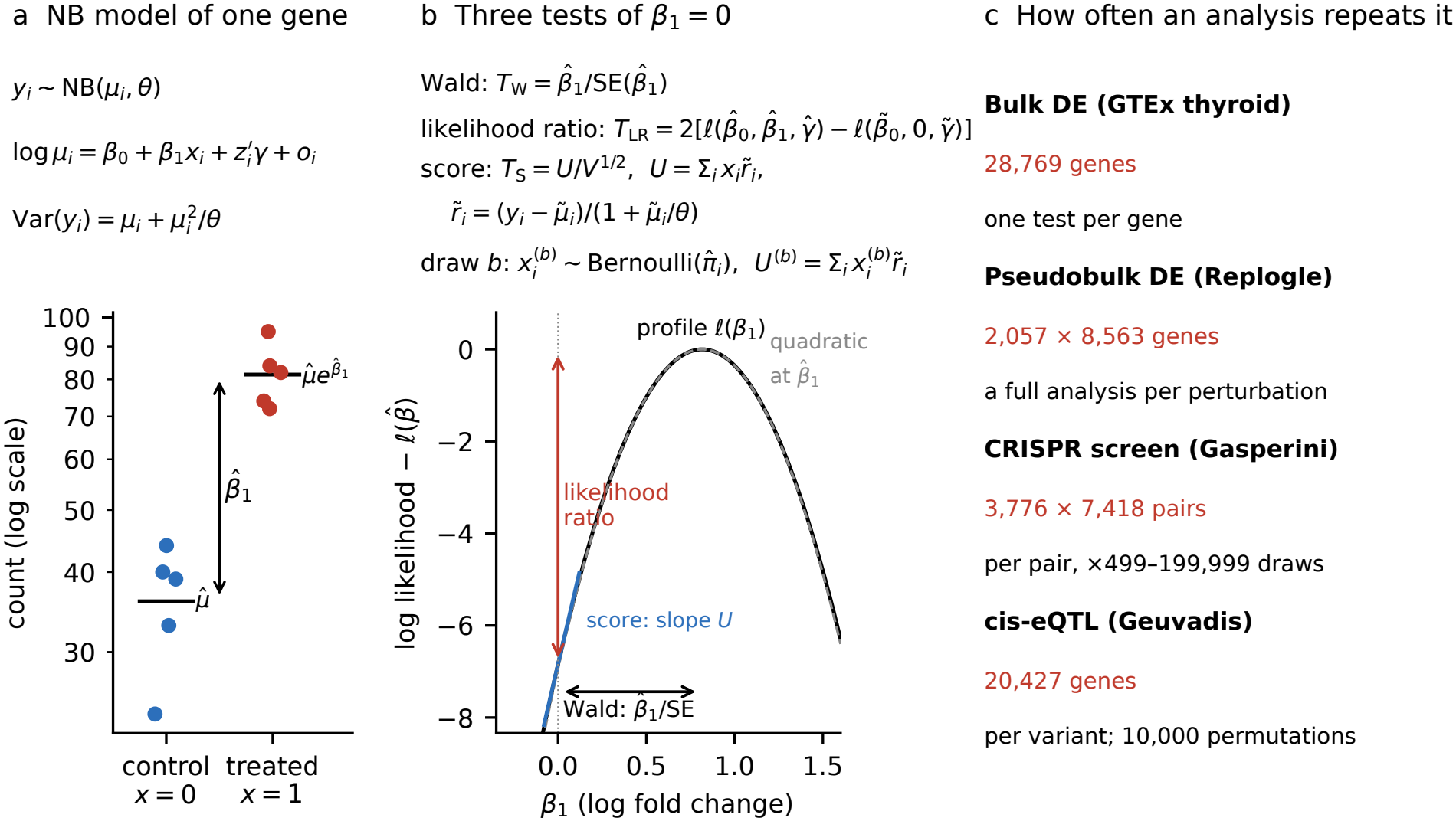
The regression behind each test, and how often it is repeated. (a) The NB model of one gene, with simulated counts of five control and five treated samples (dispersion 1*/θ* = 0.1) and their fitted means. *x*_*i*_ codes the treatment (1 for a treated sample or a perturbed cell and 0 for a control, or the number of copies of a variant’s alternative allele), *z*_*i*_ holds the other covariates, such as batch, with coefficients *γ*, and *o*_*i*_ is the log of the size factor, which accounts for sequencing depth. The variance is the Poisson variance *µ*_*i*_ of counting reads plus 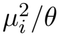 for the variation between biological replicates. A treated sample’s mean is 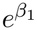 times a control’s, so *β*_1_ is the log fold change (per copy of the allele, for a variant). (b) The three tests of *β*_1_ = 0 on the log likelihood *l* of the same gene, profiled over *β*_0_. Hats mark estimates from the full model and tildes those from the null model, fitted with *β*_1_ = 0; *V* is the variance of *U* under the null hypothesis, allowing for the estimated covariate coefficients (Methods, Eq. 7). The Wald test measures the distance of 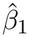 from zero in standard errors, set by the curvature at the maximum (dashed quadratic); the likelihood-ratio test, the drop from the maximum to the null; and the score test, the slope *U* at *β*_1_ = 0. Under the null hypothesis, *T*_W_ and *T*_S_ are approximately standard normal and *T*_LR_ approximately *χ*^2^ with one degree of freedom. The terms 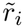 come from the null fit alone. In draw *b* of SCEPTRE’s conditional randomization test [8], each cell’s treatment is simulated with the probability 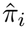 predicted by a logistic regression of the perturbation on the covariates, and only *x* changes. (c) How many times each analysis in this paper repeats the fit and the test.

The score test keeps both expensive steps down. When the tests of a gene share their cells or samples, as in QTL mapping and in screens at high MOI, one null fit serves every perturbation or variant tested against that gene. In resampling, the score of each draw is a new sum over the cells of the same null-fit terms, so no draw needs a new fit (Fig. 1b). Resampling has a limit of its own: a *p*-value counted from 9,999 draws cannot fall below 10^−4^ [9], whereas controlling the false discovery rate over millions of tests puts the discovery threshold far lower. SCEPTRE [10], a tool for CRISPR screens, therefore fits a skew-normal distribution to the statistics of the draws and takes small *p*-values from its tail [11], and cis-eQTL mappers, following FastQTL [12], fit a Beta distribution to the minimum *p*-values of each gene’s permutations.

Our methods for fast fitting and resampling are implemented in RapidGLM, a library that fits GLMs with Gaussian, Poisson, binomial or NB responses and computes Wald, likelihood-ratio and score tests, with or without resampling. A tool can use it in two ways. The simpler is to replace only the tool’s GLM library with RapidGLM. This alone should make the tool faster, because RapidGLM’s fits are faster than those of general-purpose libraries (Table 1); for example, PyDESeq2 [13] with RapidGLM as its back end, and no other change, ran 2.8 times faster on one thread (Table 4). The other is to optimize the whole analysis around the library, which gains more. We demonstrate this strategy on four widely used analyses: the DESeq2 procedure, edgeR’s quasi-likelihood workflow, the SCEPTRE method and cis-QTL mapping. The tools that implement them are described in Results. We implemented each from the method’s publications and documentation and from black-box runs of the reference tool, without reading its source (Methods). Both routes are open to other analyses built from the same steps, for example other eQTL mappers.

**Table 1:**
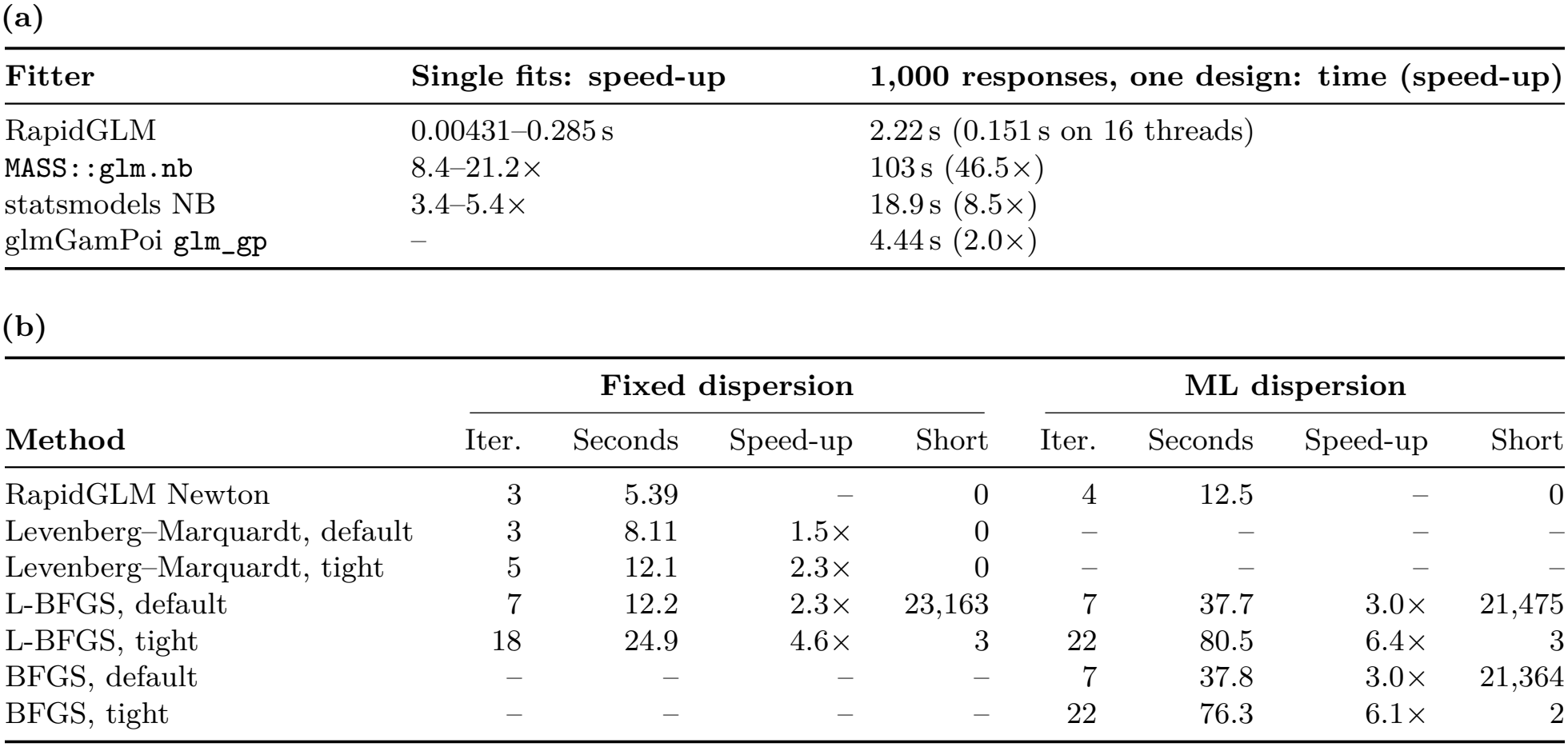
NB fits: RapidGLM against standard implementations. (RapidGLM 1.6.0; one core of an AMD Ryzen Threadripper PRO 5965WX per thread). (a) GLM packages, with the dispersion estimated, on one thread: single fits of six simulated problems (10^4^ and 10^5^ observations, 5, 20 and 50 design columns) and 1,000 responses with one design (5,000 observations, five columns); for each package, its time and RapidGLM’s speed-up over it (the package’s time over RapidGLM’s).(b) General optimizers of Ceres Solver [22] on the per-gene fits of GTEx thyroid (28,769 genes, 647 samples, 12 design columns), at a fixed dispersion and with the dispersion estimated by maximum likelihood (ML), at Ceres’s default tolerances and at tight ones: median Newton or solver iterations per gene, wall time, RapidGLM’s speed-up over each method, and the genes whose fit falls short of the best log likelihood found by more than 10^−6^ (relative). –, not run.

| **(a)** |  |  |  |  |  |  |  |  |
|---|---|---|---|---|---|---|---|---|
| **Fitter** | **Single fits: speed-up** | **1,000 responses, one design: time (speed-up)** |  |  |  |  |  |  |
| RapidGLM | 0.00431–0.285 s | 2.22 s (0.151 s on 16 threads) |  |  |  |  |  |  |
| MASS::glm.nb | 8.4–21.2× | 103 s (46.5×) |  |  |  |  |  |  |
| statsmodels NB | 3.4–5.4× | 18.9 s (8.5×) |  |  |  |  |  |  |
| glmGamPoi glm_gp | – | 4.44 s (2.0×) |  |  |  |  |  |  |
| **(b)** |  |  |  |  |  |  |  |  |
|  | **Fixed dispersion** |  |  |  | **ML dispersion** |  |  |  |
| **Method** | Iter. | Seconds | Speed-up | Short | Iter. | Seconds | Speed-up | Short |
| RapidGLM Newton | 3 | 5.39 | – | 0 | 4 | 12.5 | – | 0 |
| Levenberg–Marquardt, default | 3 | 8.11 | 1.5× | 0 | – | – | – | – |
| Levenberg–Marquardt, tight | 5 | 12.1 | 2.3× | 0 | – | – | – | – |
| L-BFGS, default | 7 | 12.2 | 2.3× | 23,163 | 7 | 37.7 | 3.0× | 21,475 |
| L-BFGS, tight | 18 | 24.9 | 4.6× | 3 | 22 | 80.5 | 6.4× | 3 |
| BFGS, default | – | – | – | – | 7 | 37.8 | 3.0× | 21,364 |
| BFGS, tight | – | – | – | – | 22 | 76.3 | 6.1× | 2 |

Our first contribution is a faster fit. General-purpose packages, from `glm` and `MASS::glm.nb` in R [14, 15] to fastglm [16], speedglm [17], statsmodels [18], scikit-learn [19] and glum [20], must handle any model, including ill-conditioned ones, and fit one model per call. The Poisson and NB models with a log link are better behaved: at a fixed dispersion, the log likelihood is concave in the coefficients and its exact second derivatives are cheap to compute. RapidGLM therefore takes Newton steps with the exact second derivatives, which converge in a few iterations, and applies its safeguards only where they are needed: it halves a step only when the step fails to improve the fit, switches to a more stable factorization only for an ill-conditioned problem, and checks the design once for all the genes fitted against it. It fits thousands of genes against one design in one call, so work that depends only on the design is done once, and it computes the exponentials and logarithms of the fits in vectorized kernels (Table 1). Our second contribution is faster resampling. Draws are generated in time proportional to the number of treated cells they contain and scored in blocks, with vectorized sums over the cells (Table 2), and the permuted statistics of QTL mapping are computed as blocked matrix products whose maxima are taken without storing them. Fits and draws run on threads that share one copy of the data, so an analysis can use every core of a machine. SCEPTRE, by contrast, runs in parallel as separate worker processes; on the Gasperini screen each held about 11 GiB, so on a 128-core compute node with 512 GB of memory it cannot use every core, and on a server with 128 GiB it can run only 10 workers.

**Table 2:**
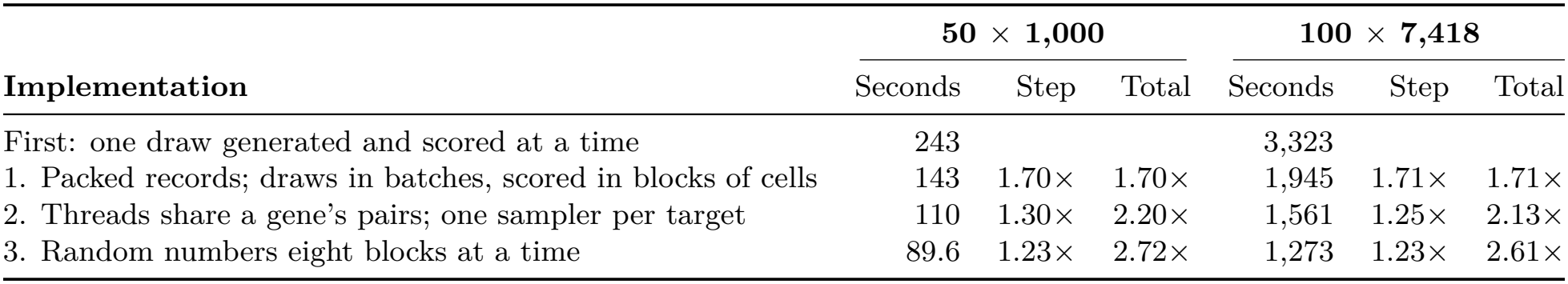
Resampling: the conditional randomization test, from the first implementation through three steps. Seconds spent on the null draws of two panels of the Gasperini screen (targets ×genes; 207,324 cells), timed with the RapidGLM 1.6.0 library, each step added to the one before, with the speed-up from the step and the total over the first implementation. The result tables were identical, byte for byte, at every step. On 16 threads, one per core: the median of three runs for the smaller panel and one run for the larger (Methods).

| Implementation | **50 × 1,000** |  |  | **100 × 7,418** |  |  |
|---|---|---|---|---|---|---|
|  | Seconds | Step | Total | Seconds | Step | Total |
| First: one draw generated and scored at a time | 243 |  |  | 3,323 |  |  |
| 1. Packed records; draws in batches, scored in blocks of cells | 143 | 1.70× | 1.70× | 1,945 | 1.71× | 1.71× |
| 2. Threads share a gene’s pairs; one sampler per target | 110 | 1.30× | 2.20× | 1,561 | 1.25× | 2.13× |
| 3. Random numbers eight blocks at a time | 89.6 | 1.23× | 2.72× | 1,273 | 1.23× | 2.61× |

Our third contribution is the optimization of each analysis as a whole. Once the fits and draws were fast, other steps came to dominate the time, such as dispersion estimation in the DESeq2 procedure and quality control in the SCEPTRE method, and each was optimized in turn [21] (Table 3). Each analysis runs entirely in compiled C++, from the counts to the results, with R and Python used only to call it. Our fourth contribution is the interfaces. A thin C layer with a stable binary interface lets any language with a C foreign-function interface use the library; the repository includes a Rust binding, tested in continuous integration. New Python and R packages are built on the same layer, with drop-in replacements for R’s `glm` and `MASS::glm.nb` and for statsmodels’ GLM, and a back end for PyDESeq2 [13]. RapidGLM is released under the MIT licence, so it can be embedded in software of any licence.

**Table 3:** What was done: the key optimizations of each part. Each part needed its own work once the part above it was fast (Discussion); details in Results and Methods.

| Part | Key optimizations |
|---|---|
| **GLM engine** | • Newton steps with the observed information and step halving; joint (β, log θ) steps with a concavity guard<br>• Jacobi-scaled Cholesky factorization, Householder QR when ill-conditioned; convergence without a confirming iteration<br>• Work that depends only on the design shared across responses (X′X, rank check, row groups, sparse rows); per-group solver for grouped designs<br>• Likelihood terms that depend only on the response by Stirling’s series; certificates for coefficients without a finite estimate<br>• Vectorized kernels: register-tiled cross products and leverages; special functions without division, with identical bits on every processor<br>• Random streams keyed by what is tested: results identical at any thread count and however a run is split |
| **DESeq2 procedure** | • Gene-wise and maximum a posteriori dispersions by Newton steps in the log dispersion, with analytic Cox–Reid derivatives<br>• Floored refits only where a fitted count falls below 0.5 or a coefficient diverges<br>• Leverages and Cook’s distances from each final fit’s factorization; genes with replaced counts re-estimated alone<br>• Genes in parallel; a PyDESeq2 back end and a PyDESeq2-style interface |
| **edgeR quasi-likelihood** | • Each fit started from the fit before it, through the passes of the workflow<br>• Convergence from a bound on the Newton decrement, without a new factorization<br>• Exact NB unit-deviance moments, tabulated for each dispersion |
| **SCEPTRE method** | • Pairwise quality control decided first, from nonzero cells; targets and responses can be named without listing pairs<br>• One null fit per response and one sampler per target, shared by their pairs<br>• Effect fits started from the null fit; pairs scheduled in parallel<br>• Treated cells drawn in time proportional to their expected number: a Poisson-process sampler with AES-based gaps, thinning as fallback<br>• Draws generated and scored 256 at a time; a ring of active pairs; Philox eight blocks at a time<br>• Batch scoring (1.6.0): all of a response’s pairs scored together, stage by stage, in one pass over its score records<br>• Refined null starts (1.6.0): at low MOI, each pair’s null started from the response’s reference fit by one Fisher-scoring step on additive totals, then refined to the full fit’s convergence criteria<br>• Optional early end to the first stage; the pairs that go on, and their results, are unchanged<br>• A run divided into tiles, checkpoints or machines merges into the table of an undivided run |
| **cis-QTL mapping** | • All variants of a gene scored in one cross-product pass; the parts that depend only on the weights computed once per gene<br>• Permutations scored 512 at a time in single precision, maxima taken in registers; observed statistics and calibration in double precision<br>• Shuffles from vectorized Philox with bounded integers drawn without division<br>• Genotypes kept as two-bit codes; per-thread workspaces reused from gene to gene<br>• Count models (1.6.0): permutations scored eight at a time in one blocked product, and their nulls refitted in lockstep |

**Table 4:** One analysis through every interface: the DESeq2 procedure on GTEx thyroid. (647 samples, 28,769 genes; RapidGLM 1.6.0). Wall-clock seconds at one and eight threads (*T*), with the speed-up over the tool replaced, and the discoveries at a false discovery rate of 0.1. The second column names the part of the analysis that RapidGLM computes. The command-line program’s time includes reading the input files; the others are times of the calls (Methods).

| **Called as** | **Runs in RapidGLM** | *T*=1 | *T*=8 | **Discoveries** |
|---|---|---|---|---|
| *In place of DESeq2* |  |  |  |  |
| DESeq2 | – | 401 | 70.2 | 7,177 |
| RapidGLM, command line | whole analysis | 29.4 (13.6×) | 5.29 (13.3×) | 7,178 |
| RapidGLM, Python package | whole analysis | 28.9 (13.9×) | 4.88 (14.4×) | 7,178 |
| RapidGLM, R package | whole analysis | 30.0 (13.4×) | 5.54 (12.7×) | 7,178 |
| *In place of PyDESeq2* |  |  |  |  |
| PyDESeq2 | – | 127 | 24.4 | 7,177 |
| RapidGLM, PyDESeq2-style interface | whole analysis | 29.4 (4.3×) | 5.43 (4.5×) | 7,178 |
| PyDESeq2 with RapidGLM as back end | fits, dispersions, Wald tests | 45.7 (2.8×) | 12.5 (1.9×) | 7,176 |

The four analyses agree with the reference tools and run 3.0 to 17.8 times faster on the same cores (Table 5). At genome scale (Table 6), RapidGLM’s DESeq2 procedure analysed all 2,057 Replogle perturbations in 1.09 hours, against 5.6 hours for DESeq2. On one compute node, RapidGLM tested every target–gene pair of the Gasperini screen in 44 minutes with SCEPTRE’s resampling scheme, in which the pairs of a target share their draws, and in 2.8 hours with its own defaults, against 11.6 hours for SCEPTRE. These runs took 90 and 360 core-hours, against 1,480 for SCEPTRE. The defaults spend part of the time saved on the accuracy of the small *p*-values that decide discoveries, on which the false discovery rate depends. They simulate independent draws for every pair, accept a fitted tail only where the statistics of the draws on the side of the observed statistic are no heavier-tailed than the fit, and confirm the smallest *p*-values with 199,999 draws.

**Table 5:** The four analyses against their reference tools. (RapidGLM 1.6.0). Wall-clock seconds of the reference tool and of RapidGLM on the same data and the same *T* threads, one per physical core, with RapidGLM’s speed-up. The Jaccard index is the number of discoveries made by both tools divided by the number made by either, over the genes or pairs that both tools tested, at a false discovery rate of 0.1; for cis-QTL mapping, the discoveries are the genes with an association at *q <* 0.05. The cis-QTL scans used 10,000 permutations for normalized expression and 1,000 for counts; tensorQTL and jaxQTL ran on CPUs. ^a^edgeR runs on one thread, and RapidGLM’s time on eight threads is compared with that time. ^b^FastQTL ran as twelve single-thread processes, and was timed with RapidGLM in a later run (Methods).

| **Data** | **Reference tool** | *T* | **Reference** | **RapidGLM** | **Speed-up** | **Jaccard** |
|---|---|---|---|---|---|---|
| *DESeq2 procedure* |  |  |  |  |  |  |
| airway, 8 samples | DESeq2 | 1 | 5.12 | 0.847 | 6.1× | 0.986 |
|  |  | 8 | 2.48 | 0.244 | 10.1× |  |
| GTEx thyroid, 128 samples | DESeq2 | 1 | 95.9 | 8.27 | 11.6× | 0.997 |
|  |  | 8 | 26.3 | 1.51 | 17.4× |  |
| GTEx thyroid, 647 samples | DESeq2 | 1 | 401 | 30.0 | 13.4× | 1.000 |
|  |  | 8 | 70.2 | 5.54 | 12.7× |  |
| *edgeR quasi-likelihood workflow* |  |  |  |  |  |  |
| GTEx thyroid, 647 samples | edgeR | 1 | 77.7 | 8.17 | 9.5× | 0.999 |
|  |  | 8 | 77.7<sup>a</sup> | 3.34 | 23.3× |  |
| *SCEPTRE method, high MOI* |  |  |  |  |  |  |
| Gasperini panel, 50,000 cells | SCEPTRE | 1 | 1,380 | 246 | 5.6× | 0.846 |
|  |  | 8 | 207 | 35.9 | 5.7× |  |
| Gasperini panel, 207,324 cells | SCEPTRE | 8 | 1,108 | 182 | 6.1× | 0.895 |
| *SCEPTRE method, low MOI* |  |  |  |  |  |  |
| Replogle pilot, 13,618 cells | SCEPTRE | 16 | 2,159 | 227 | 9.5× | 0.949 |
| *cis-QTL mapping, Geuvadis chromosome 22* |  |  |  |  |  |  |
| Normalized expression | FastQTL | 12 | 634<sup>b</sup> | 35.6 | 17.8× | 0.993 |
|  | tensorQTL | 12 | 107 | 35.9 | 3.0× | 0.996 |
| Counts | jaxQTL | 12 | 2,257 | 498 | 4.5× | 0.988 |

**Table 6:**
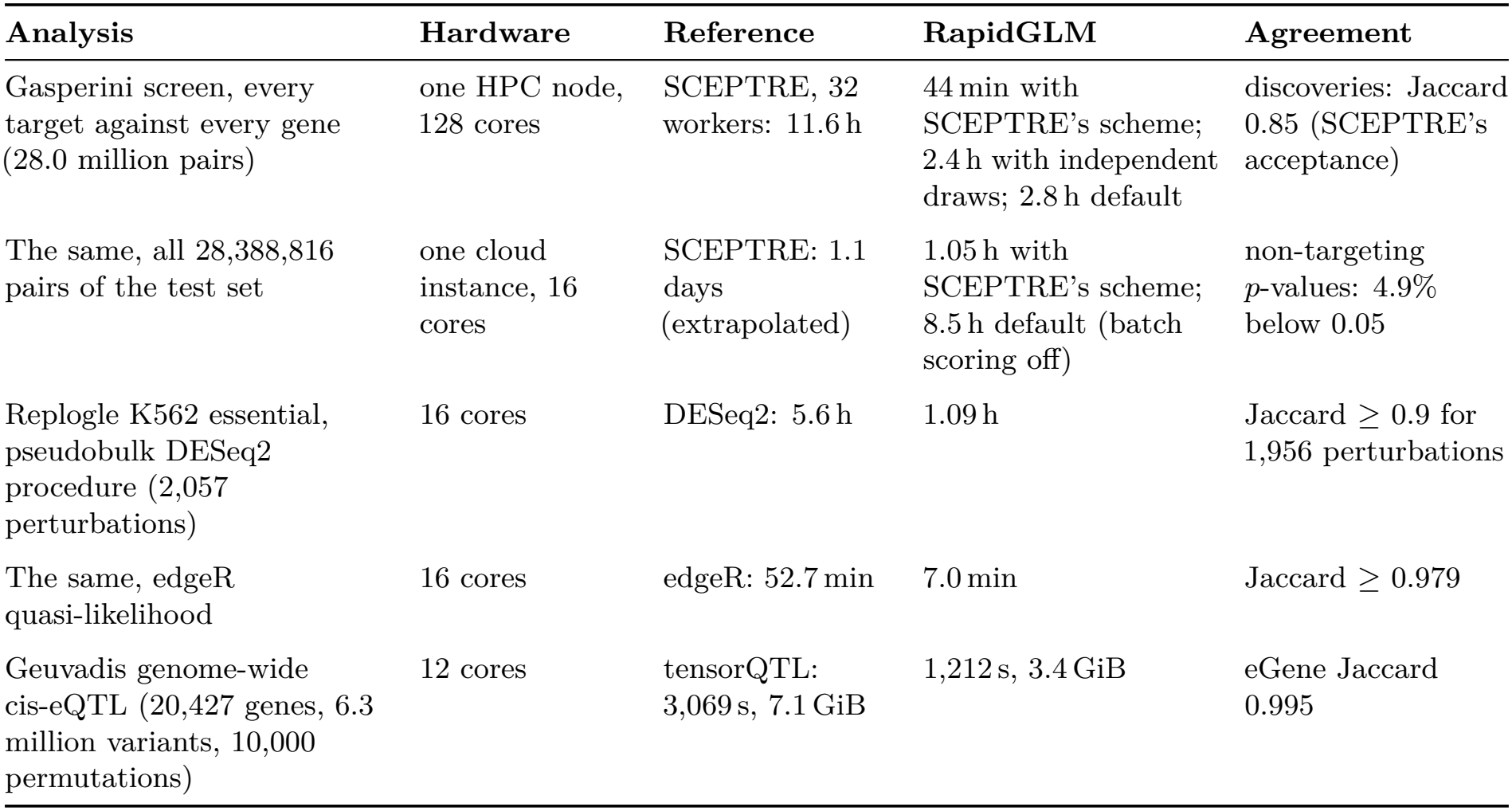
Analyses at genome scale. (RapidGLM 1.6.0). Wall-clock times; RapidGLM’s default unless stated; one thread per physical core. The HPC nodes have two 64-core AMD EPYC 7742 processors and 512 or 256 GB (Methods).

| **Analysis** | **Hardware** | **Reference** | **RapidGLM** | **Agreement** |
|---|---|---|---|---|
| Gasperini screen, every target against every gene (28.0 million pairs) | one HPC node, 128 cores | SCEPTRE, 32 workers: 11.6 h | 44 min with SCEPTRE's scheme; 2.4 h with independent draws; 2.8 h default | discoveries: Jaccard 0.85 (SCEPTRE's acceptance) |
| The same, all 28,388,816 pairs of the test set | one cloud instance, 16 cores | SCEPTRE: 1.1 days (extrapolated) | 1.05 h with SCEPTRE's scheme; 8.5 h default (batch scoring off) | non-targeting *p*-values: 4.9% below 0.05 |
| Replogle K562 essential, pseudobulk DESeq2 procedure (2,057 perturbations) | 16 cores | DESeq2: 5.6 h | 1.09 h | Jaccard ≥ 0.9 for 1,956 perturbations |
| The same, edgeR quasi-likelihood | 16 cores | edgeR: 52.7 min | 7.0 min | Jaccard ≥ 0.979 |
| Geuvadis genome-wide cis-eQTL (20,427 genes, 6.3 million variants, 10,000 permutations) | 12 cores | tensorQTL: 3,069 s, 7.1 GiB | 1,212 s, 3.4 GiB | eGene Jaccard 0.995 |

## Results

The sections below follow the contributions set out in the Background: the engine’s fits, then resampling, then the interfaces, then the four analyses against their reference tools, and finally analyses of whole genome-scale datasets. Unless an earlier release is named, all measurements are of RapidGLM 1.6.0, and each optimization new in 1.6.0 was also measured against the same build with it switched off.

### Fast and accurate fits

RapidGLM fits count models with an aggressive form of Newton’s method: full steps with the exact second derivatives of the log likelihood, a step halved only when it fails to improve the fit, and convergence declared without a confirming iteration. Work that depends only on the design is shared by all the responses fitted against it, and the remaining loops run in vectorized kernels (Methods). Table 1 compares its NB fits with standard implementations, and Table 3 lists these optimizations and those of each analysis built on the engine.

#### Fewer iterations, less time

On the per-gene NB fits of GTEx thyroid, RapidGLM converged in a median of 3 iterations at a fixed dispersion and 4 with the dispersion estimated, against 5–18 and 22 for the general optimizers of Ceres Solver at the same accuracy, and was 2.3–6.4 times faster than them (Table 1b). At their default tolerances, L-BFGS and BFGS stopped short of the best log likelihood for most genes. Against GLM packages, single NB fits were 8.4–21.2 times faster than `MASS::glm.nb` and 3.4–5.4 times faster than statsmodels, and 1,000 NB responses with one design took 2.22 s on one thread, 2.0, 8.5 and 46.5 times less than glmGamPoi, statsmodels and `MASS::glm.nb,` and 0.15 s on 16 threads (Table 1a). Over airway, a simulation and thyroid, RapidGLM was 2.0–3.1 times faster than Levenberg–Marquardt at a fixed dispersion, and 1.8–6.1 times faster than the faster of BFGS and L-BFGS with the dispersion estimated, all at tight tolerances. For Poisson, binomial and Gaussian models, single fits were a median of 6.5, 8.1 and 2.7 times faster than glm.fit, statsmodels and fastglm (Additional file 1).

#### No loss of accuracy

On 39 identifiable problems with reference fits computed in IEEE binary128 arithmetic, covering ill-conditioned designs and other edge cases (Methods), RapidGLM’s largest coefficient error was 1.7 × 10^−6^ reference standard errors and its standard errors were within 2.1 × 10^−8^ (relative), against 1.9 × 10^−5^ for R’s `glm.fit.` On the 13 NB problems its largest error was 1.3 × 10^−6^, against 1.5 × 10^−4^ for `MASS::glm.nb,` and statsmodels failed on 8 (Additional file 1, Section S1). Small experiments are harder: on airway, whose eight samples leave many dispersion likelihoods flat, 198 of 22,369 fits with the dispersion estimated ended more than 10^−6^ (relative) short of the best log likelihood found, by at most 6.0 × 10^−4^ in log likelihood, against 6 fits for BFGS at tight tolerances.

### Fast resampling

Beyond the fits, most of the time of a screen or a QTL scan goes to resampling: generating the draws and scoring each against the fitted null. The costliest case is the conditional randomization test of a screen at high MOI, with hundreds of draws or more for each of millions of pairs. RapidGLM’s first implementation generated a draw, scored it and went on to the next; three steps made it faster (Table 2). (1) Each cell’s values are packed into one record, and draws are generated in batches and scored in blocks of cells. Scoring a draw adds up a gene’s per-cell values over the cells that the draw treats, which are scattered through memory; a packed record takes one read where separate arrays took three, and in a batch each block of records is read once for all the draws, not once per draw. (2) The threads work together on a few pairs of one gene at a time, so that the gene’s values stay in the cache that the threads share, and a target’s sampler is built once for all its pairs, not once per pair. (3) The random numbers are computed eight blocks at a time with vector instructions, which speeds up the arithmetic of the generator. The details, two later changes to the sampler and to scoring, and the permutations of the other analyses are in Methods (Engine: Resampling; SCEPTRE method and cis-QTL mapping: Optimizations).

#### The gain from each step

Together the three steps made the draws 2.72 times faster on 50 targets against 1,000 genes of the Gasperini screen, and 2.61 times faster on 100 targets against all 7,418 genes, on the same threads (Table 2). Every step contributed, the first the most. Two later changes followed (Methods): drawing the treated cells of a draw by a Poisson process made the draws a further 1.34 times faster, and scoring all the pairs of a gene in one pass over its cells made whole runs 1.55 times faster.

#### Low MOI and QTL mapping

In the other resampling analyses the time lay elsewhere, and each optimization was measured by switching it off in RapidGLM 1.6.0. At low MOI each pair’s null model is fitted on its own cells (the target’s and the non-targeting controls’), and these fits dominated. Starting each from the gene’s fit on all cells made a pilot of 20 perturbations of the Replogle screen (174,761 pairs) 2.30 times faster when small *p*-values were accepted as SCEPTRE accepts them, and 1.25 times faster with RapidGLM’s default, which draws more (below). In QTL mapping, the permutations of a gene are scored together as matrix products. Scoring them in single precision made the Gaussian scan of Geuvadis chromosome 22 1.7 times faster, and for count models, where every permutation needs a new null fit, refitting eight nulls at a time made the NB scan 1.09 times faster. The comparisons with SCEPTRE, FastQTL, tensorQTL and jaxQTL on the same cores follow below (Table 5).

#### The same results

The three steps of Table 2, and scoring all the pairs of a gene together, gave byte-identical tables. The Poisson-process sampler uses different random numbers from the sampler it replaced, so its results differ as those of two seeds do. The Gaussian QTL scan found the same eGenes (genes with a significant association) in single and in double precision, and the NB scan the same 168 eGenes whether the permutation nulls were refitted together or separately, with Beta-approximated *p*-values within 7.9 × 10^−11^ (relative). With the null fits started from the gene’s fit, the statuses, empirical *p*-values and discoveries (10,979 with the default and 10,982 with SCEPTRE’s acceptance) were identical; *p*-values from the fitted skew-normal tail, which depend continuously on the statistic of every draw, moved by more than 10^−4^ (relative) for 44 and 72 pairs, respectively, all with *p* below 2.7 × 10^−17^. Results also do not depend on how a run is divided. Every draw comes from a counter-based random stream [23] keyed by the seed and by what is tested, the perturbation–gene pair or the QTL gene, so a test’s result is identical at any thread count, whichever other tests share the run, and when a run is split into tiles or jobs and merged. At genome scale this is what allows one analysis to be spread over machines (below).

#### Accurate small *p*-values

Speed also buys draws. SCEPTRE reads a small *p*-value from a skew-normal curve fitted to the statistics of its draws. On 100 targets against all 7,418 genes of the Gasperini screen, counting the tail of the draws directly for 26 pairs, over 1–4 million draws each, contradicted the fitted *p*-value for 7, all with positive statistics, by factors of 8 to 7,500; SCEPTRE reported 5 of them as discoveries. RapidGLM’s default therefore accepts a fitted *p*-value only if the draws in the tail on the statistic’s side are no heavier than the fit says, and confirms the smallest *p*-values on 199,999 draws (Methods). SCEPTRE’s acceptance remains available (`fit_check=ks`) and is used for the comparisons with SCEPTRE in Table 5. The default took 16–46% longer on the Gasperini panels of Table 5, and 1,058 s against 227 s on the Replogle pilot.

### Interfaces

RapidGLM was originally designed to be used with our C++ software suites as an easily integrated back end. We provided a thin C layer with a stable binary interface that exposes the engine, so that any language that can call C can use the library in the same way (Fig. 2). The repository includes Rust, Python and R bindings and a command-line program, which can be used to call RapidGLM directly. Alternatively, it is possible to use an existing package such as PyDESeq2 and replace its back end with RapidGLM. The C layer and the packages are described in Methods (Engine: Interfaces) and the results are shown in Table 4. Direct calls are faster, but it may be more convenient to use an existing package with a well-established API such as PyDESeq2.

**Figure 2:**
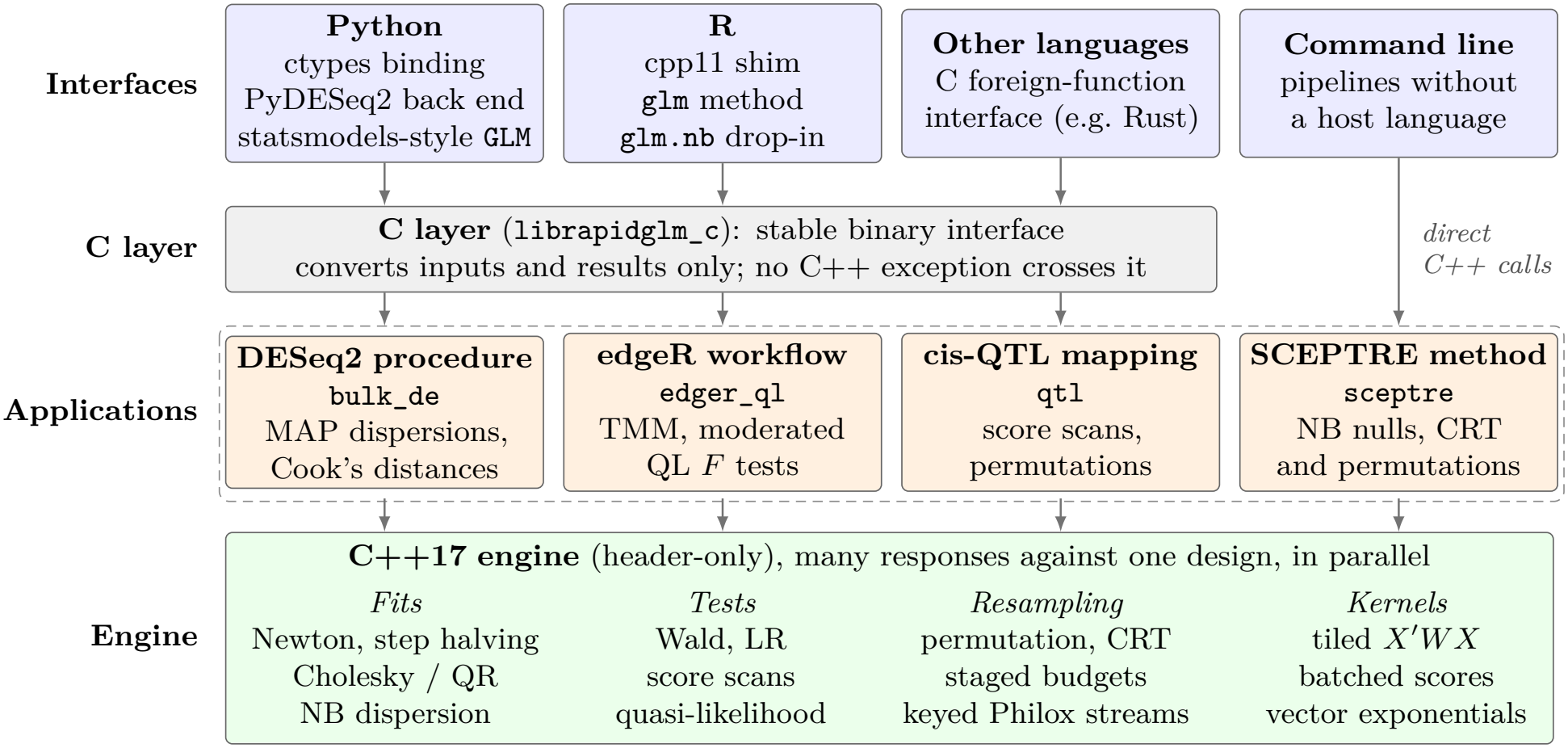
RapidGLM’s layers. The engine fits and tests many responses against one design, and the four analyses are built on it. The Python and R packages, and programs in other languages, call the analyses and the engine through the C layer, which converts inputs and results without changing them. The command-line program is written in C++ and calls them directly.

### Four analyses against their reference tools

RapidGLM’s four analyses each follow an established method, and each was benchmarked against the tools that implement that method (Table 5). DESeq2 [3] and edgeR [4, 5, 24] find the genes that are differentially expressed between conditions, in bulk RNA-seq samples or in pseudobulk samples, the summed counts of groups of single cells. Both fit an NB model to each gene. Most experiments have too few samples to estimate a gene’s dispersion from that gene alone, so both tools borrow information from the other genes. DESeq2 then tests each gene with the Wald or the likelihood-ratio test, and edgeR’s quasi-likelihood workflow with an *F*-test. PyDESeq2 [13] implements the DESeq2 procedure in Python (Table 4). SCEPTRE [10, 11] analyses single-cell CRISPR screens. For each pair of a target and a gene, it compares the gene’s counts in the cells that carry a guide against the target with its counts in control cells, using the NB score test, and estimates the *p*-value by resampling: the conditional randomization test at high MOI and permutations at low MOI. FastQTL [12] maps cis-eQTLs. It regresses each gene’s normalized expression on each variant near the gene, and estimates the *p*-value of the gene’s strongest association from permutations. tensorQTL [25] implements the same method for GPUs. jaxQTL [26], also designed for GPUs, tests the counts themselves, with the score test of an NB model. RapidGLM’s implementation of each method is described in Methods.

RapidGLM was 3.0 to 17.8 times faster than the reference tools on the same cores. DESeq2 and edgeR are deterministic, and RapidGLM’s discoveries were nearly identical to theirs (Jaccard index at least 0.986). The *p*-values of SCEPTRE are estimated from random draws, so two runs with different random seeds do not make exactly the same discoveries. On the Gasperini panel of 207,324 cells, RapidGLM runs with six different seeds overlapped SCEPTRE’s discoveries with Jaccard indices of 0.90–0.96, and overlapped one another with 0.90–0.97. The comparisons on further datasets are in the Methods section of each analysis.

### Analyses at genome scale

Table 6 collects the analyses of whole datasets.

### Every perturbation of the Replogle K562 essential screen

For each of the 2,057 pertur-bations we ran a complete pseudobulk analysis against the non-targeting samples (Table 7), with the DESeq2 procedure and with edgeR’s quasi-likelihood workflow, each tool as 16 single-thread workers taking perturbations from one queue. The DESeq2 procedure took 1.09 h against 5.6 h for DESeq2 (5.1×), at a median peak memory of 0.04 GB per perturbation against 0.89 GB; edgeR’s workflow took 7.0 min against 52.7 min for edgeR (7.5×). The discoveries of every perturbation agreed with edgeR’s with Jaccard of at least 0.979, and with DESeq2’s with Jaccard of at least 0.9 for 1,956 of them. In the others that we inspected, the *p*-values rank identically, and the discoveries differ through the independent-filtering threshold each tool chose.

**Table 7:**
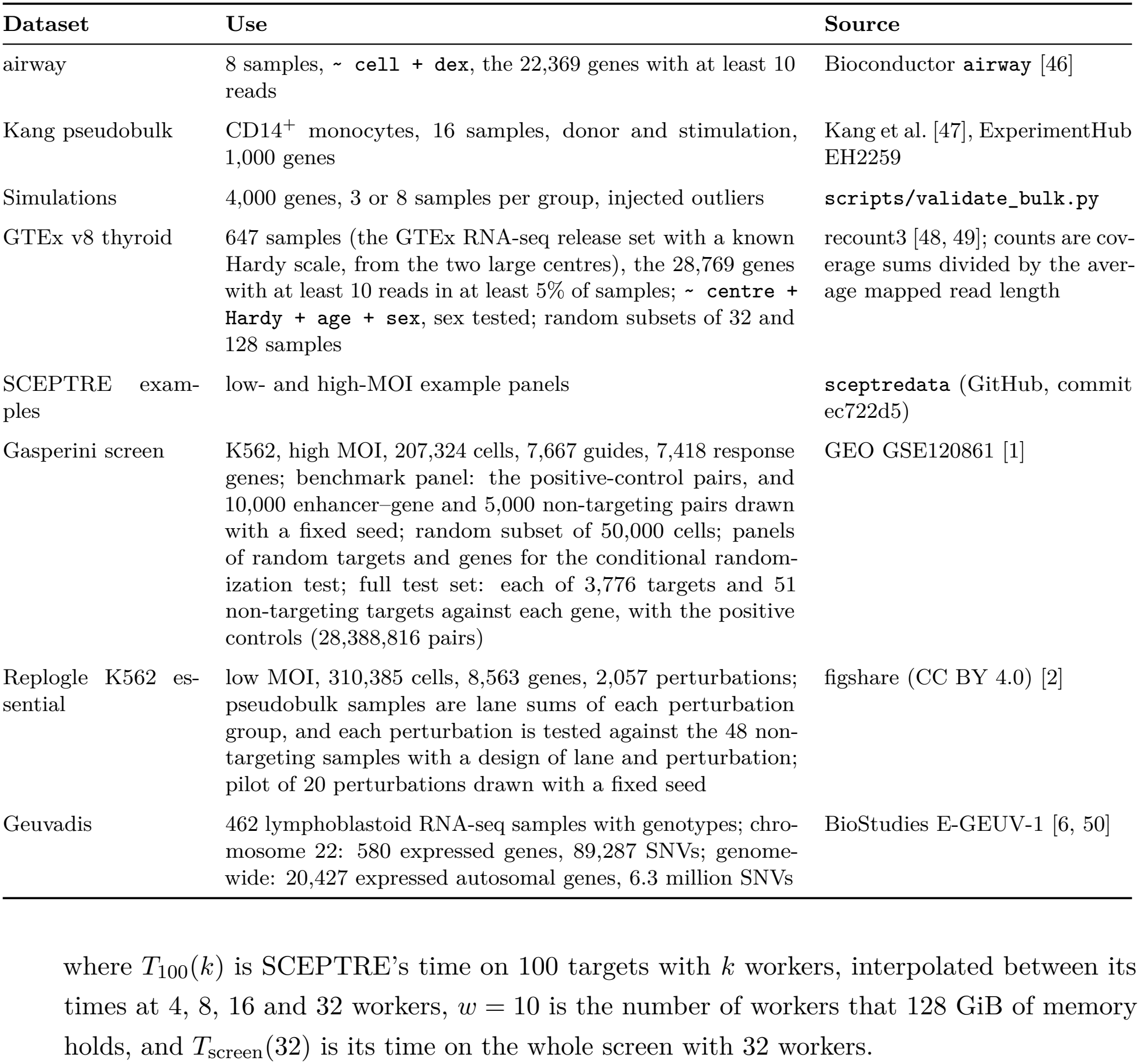
Datasets.

| Dataset | Use | Source |
|---|---|---|
| airway | 8 samples, ~ cell + dex, the 22,369 genes with at least 10 reads | Bioconductor airway [46] |
| Kang pseudobulk | CD14<sup>+</sup> monocytes, 16 samples, donor and stimulation, 1,000 genes | Kang et al. [47], ExperimentHub EH2259 |
| Simulations | 4,000 genes, 3 or 8 samples per group, injected outliers | scripts/validate_bulk.py |
| GTEx v8 thyroid | 647 samples (the GTEx RNA-seq release set with a known Hardy scale, from the two large centres), the 28,769 genes with at least 10 reads in at least 5% of samples; ~ centre + Hardy + age + sex, sex tested; random subsets of 32 and 128 samples | recount3 [48, 49]; counts are cov-erage sums divided by the aver-age mapped read length |
| SCEPTRE exam-ples | low- and high-MOI example panels | sceptredata (GitHub, commit ec722d5) |
| Gasperini screen | K562, high MOI, 207,324 cells, 7,667 guides, 7,418 response genes; benchmark panel: the positive-control pairs, and 10,000 enhancer–gene and 5,000 non-targeting pairs drawn with a fixed seed; random subset of 50,000 cells; panels of random targets and genes for the conditional random-ization test; full test set: each of 3,776 targets and 51 non-targeting targets against each gene, with the positive controls (28,388,816 pairs) | GEO GSE120861 [1] |
| Replogle K562 es-sential | low MOI, 310,385 cells, 8,563 genes, 2,057 perturbations; pseudobulk samples are lane sums of each perturbation group, and each perturbation is tested against the 48 non-targeting samples with a design of lane and perturbation; pilot of 20 perturbations drawn with a fixed seed | figshare (CC BY 4.0) [2] |
| Geuvadis | 462 lymphoblastoid RNA-seq samples with genotypes; chro-mosome 22: 580 expressed genes, 89,287 SNVs; genome-wide: 20,427 expressed autosomal genes, 6.3 million SNVs | BioStudies E-GEUV-1 [6, 50] |

#### A genome-wide cis-eQTL scan

On all 20,427 expressed genes of Geuvadis (20,248 with cis variants) and 6.3 million variants, the Gaussian scan with 10,000 permutations took 1,212 s on twelve cores at a peak of 3.4 GiB, against 3,069 s and 7.1 GiB for tensorQTL on the same cores (2.53×). It found 9,575 eGenes against tensorQTL’s 9,581 (Jaccard 0.995), with the same lead variant for 98.6% of genes. Every gene’s status, lead variant and observed statistic, and the eGenes, are those of a double-precision scan (RapidGLM 1.4.1); single-precision permutation scores moved 2,860 empirical *p*-values, each by at most 5 permutation counts.

#### Every pair of the Gasperini screen

The largest test is the whole Gasperini test set: each of the 3,776 targets against each of 7,418 genes, 28.0 million discovery pairs, of which 27.2 million passed pairwise quality control, with the benchmark panel’s 5,130 control pairs (Table 6, Fig. 3). We ran SCEPTRE and RapidGLM on it, and on 100 targets against every gene and on the benchmark panel, on nodes of the Bridges-2 supercomputer, each with two 64-core processors and 512 or 256 GB of memory (Methods). SCEPTRE runs in parallel in forked workers, each of which held about 11 GiB; on 32 workers it took 11.6 h at a peak of 302 GiB. RapidGLM’s threads share one copy of the data, and it used all 128 cores. With SCEPTRE’s resampling scheme, in which the pairs of each target share their draws and nothing is confirmed on further draws, RapidGLM took 44 min (16× faster), or 44 min with SCEPTRE’s acceptance of the fitted tail as well (16×). With independent draws for every pair and SCEPTRE’s acceptance it took 2.4 h at a peak of 400 GiB (4.8×), and with its defaults, which add the tail check and the confirmation draws, 2.8 h (4.1×). On 32 threads, as many as SCEPTRE’s workers, it took 51 min with SCEPTRE’s scheme (14×) and 5.0 h with independent draws (2.3×). Batch scoring, in tiles of 1,280 targets, made the run with independent draws 1.7 times faster (2.4 against 4.0 h) at a peak of 400 against 43 GiB. Time on shared HPC systems is allocated by the core-hour, and since a job is charged for the cores its memory keeps from others, the whole screen cost SCEPTRE 1,480 core-hours, and RapidGLM 90 with SCEPTRE’s scheme and 360 with its defaults.

**Figure 3:**
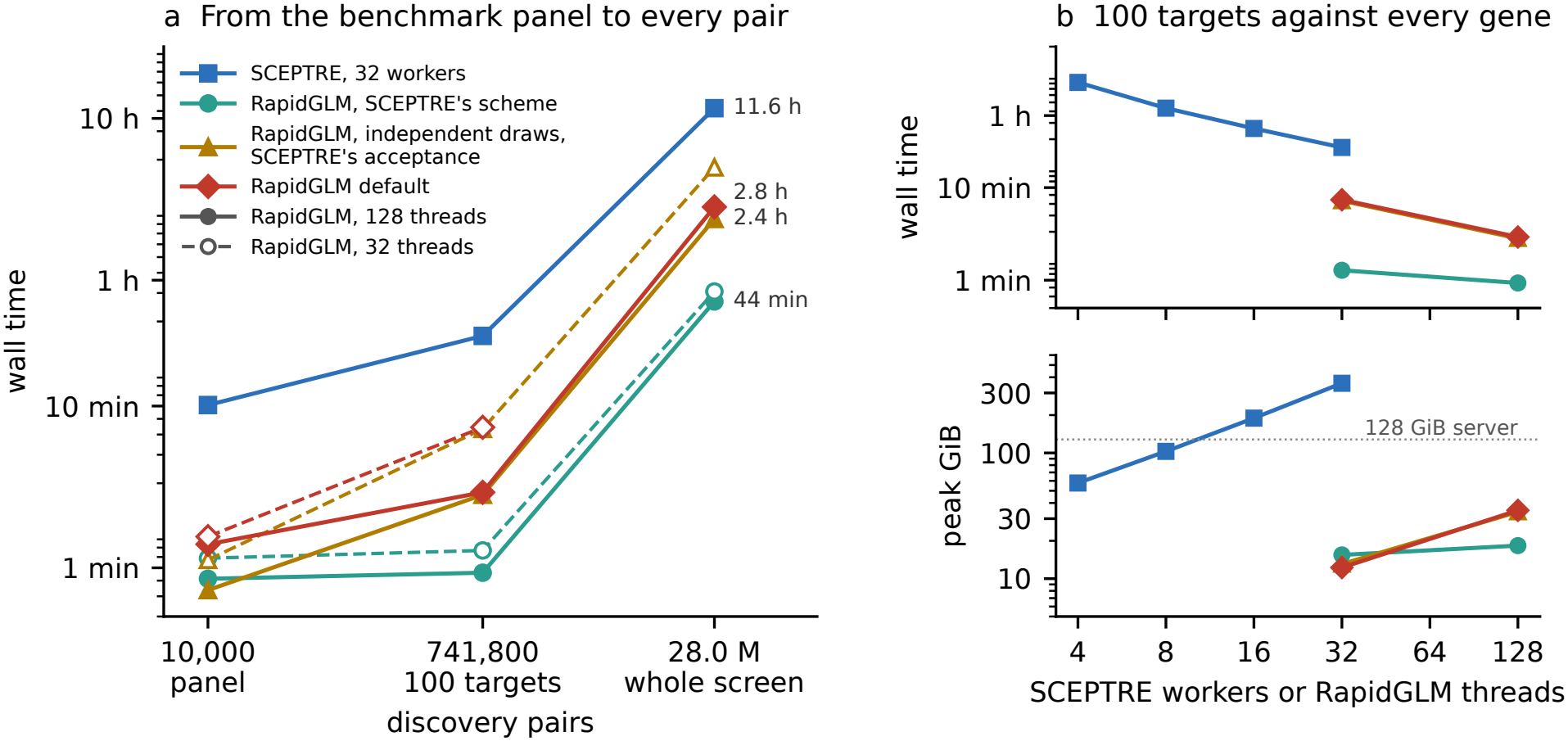
SCEPTRE and RapidGLM on the same processors, from the benchmark panel to the whole Gasperini screen. (a) Wall time against the number of discovery pairs (the benchmark panel; 100 targets against every gene; every pair of the test set) for SCEPTRE on 32 workers and RapidGLM on 128 threads (solid) and 32 threads (dashed), with SCEPTRE’s resampling scheme, with independent draws and SCEPTRE’s acceptance, and with RapidGLM’s default. (b) Time and peak memory on 100 targets against every gene, for SCEPTRE at 4–32 workers and RapidGLM at 32 and 128 threads; the dotted line marks 128 GiB. Bridges-2 nodes with two 64-core AMD EPYC 7742 processors.

#### On a smaller machine

The memory also sets what a smaller machine can do. SCEPTRE’s time on 100 targets fell 5.0-fold from 4 to 32 workers (Fig. 3b); scaled from there to the whole screen, on a server with 16 cores and 128 GiB, which holds 10 workers, it would take about 1.1 days (an extrapolation, Methods). On a cloud instance of that size, RapidGLM ran the whole test set, with every non-targeting pair (28,388,816 pairs), in 1.05 h with SCEPTRE’s scheme at a peak of 32.1GiB and in 8.5 h with its defaults at 40.0 GiB, with batch scoring switched off (Methods). Each release since 1.4.1 was run on the same instance type: with SCEPTRE’s scheme the run took 4.7 h in 1.4.1 and 4.6 h in 1.5.0, and with the defaults 16.0 and 9.6 h. The 1.6.0 tables equal 1.5.0’s byte for byte in both modes.

#### Agreement on the whole screen

Over the 27.2 million discovery pairs both tools tested, RapidGLM with SCEPTRE’s acceptance and independent draws made 4,453 discoveries at a false discovery rate of 0.1 and SCEPTRE 4,628, 4,182 of them shared (Jaccard 0.85; 0.86 with shared draws, 0.80 with SCEPTRE’s scheme and RapidGLM’s tail check, and 0.79 with the defaults), and the *p*-values of every mode agreed with SCEPTRE’s with Spearman correlation of at least 0.985. Each mode’s table was the same, byte for byte, on 32 and 128 threads (for the defaults, on the smaller sets) and, with independent draws, with batch scoring on or off. On a check panel of 148,490 pairs, the *p*-values equal those of the same pairs in the cloud runs. With the defaults the whole test set made 3,791 discoveries at a false discovery rate of 0.1, and with SCEPTRE’s scheme 3,886, 3,629 of them shared; the default run drew 18.1 billion draws and confirmed 5,351 pairs on 199,999 draws. Of the 368,579 tested non-targeting pairs, 4.9% had *p <* 0.05 and one was a discovery, and 35 of 128 positive controls were recovered. At this scale the discovery threshold lies beyond what an ordinary resampling budget can count. The largest *p*-value called was 1.4 × 10^−5^, below the 10^−4^ floor of a two-sided empirical *p*-value from 19,999 draws, so every discovery rests on a fitted tail or on deeper sampling, and the speed of the engine is what makes the deeper sampling affordable.

## Discussion

### What the speed is for

Most of these analyses were originally designed for a small set of conditions or perturbations, or for selected target–gene pairs. With single-cell technologies, thousands of perturbations can now be applied in one experiment and tested against every gene, in hundreds of thousands of cells, which calls for a scalable implementation. The Gasperini screen shows the scale. Testing each of its 3,776 targets against each of 7,418 genes, in 207,324 cells, takes 28.0 million tests, each with hundreds of draws or more. SCEPTRE took 11.6 hours for them on a 128-core supercomputer node, and would take about 1.1 days on a 16-core server, whose memory holds 10 of its workers (an extrapolation; Table 6). With the same resampling scheme, RapidGLM took 44 minutes on the node and 1.05 hours on a cloud instance of the server’s size. An analysis of a day or more becomes one of an hour, which can be repeated whenever its design, its filters or its seed change. A larger screen can be divided over machines and merged, because a result does not depend on the number of threads or on how a run is divided.

#### Speed and accuracy

Where each draw is expensive, a method has to compromise between time and accuracy. In SCEPTRE’s resampling scheme, all the pairs of a target share one set of draws, so the errors of their *p*-values are correlated, and a *p*-value too small to count is read from a skew-normal curve fitted to the draws, with no further draws to confirm it. A faster implementation does not need these compromises. By default, RapidGLM gives every pair its own draws, accepts a fitted curve only where the draws in its tail are no heavier than the curve says, and confirms the smallest *p*-values on 199,999 draws. The difference shows in the results. When we counted the tail directly for 26 pairs, over millions of draws each, the fitted *p*-value was too small for 7 of them, by factors of 8 to 7,500, and SCEPTRE reported 5 of these as discoveries (Results, Accurate small *p*-values). With these defaults the whole screen took 2.8 hours on the node, still 4.1 times faster than SCEPTRE.

#### Why a specialized engine

General-purpose GLM packages must handle any model, and fit one model per call. The count models of these analyses have structure that a general package cannot assume. At a fixed dispersion the log likelihood is concave in the coefficients, so Newton steps with the exact second derivatives converge in a few iterations, and a safeguard is applied only when a step needs it. Thousands of genes share one design, so the work that depends only on the design is done once. What remains is arithmetic over the samples, which vectorizes (Table 1).

#### From a fast engine to a fast analysis

A fast engine does not by itself make an analysis fast. Amdahl’s law [21] limits the gain from speeding up one part of a computation by the time spent in the rest: if the fits take 80% of an analysis, making them instantaneous gains at most a factor of five. Each optimization also changes which part dominates, so the next gain has to come from wherever the time has moved. Once the fits were fast, the time moved to a different place in each analysis (Table 3): to the dispersion estimates in the DESeq2 procedure; to the number of fits in edgeR’s workflow, which fits every gene again in each of its passes; to the draws at high MOI, and to the null fit of every pair at low MOI, in the SCEPTRE method; and to the scores of the permutations in cis-QTL mapping. At genome scale, memory became a limit as well (Methods). Each analysis was therefore optimized as a whole: its time was measured, the largest cost was removed, and its time was measured again. Table 4 shows the difference on one analysis. As the back end of PyDESeq2, RapidGLM replaces the fits, the dispersion estimates and the Wald tests of an otherwise unchanged workflow; with the whole workflow in the engine, behind the same kind of calls, the gain is larger. The interfaces offer this choice: a user can replace one call, one back end or a whole analysis, according to how much of a program they are willing to change.

#### Limitations

The four analyses were written from publications, documentation and outputs, not from the reference tools’ code, so a rule that none of these reveals can differ. Agreement is therefore measured for each analysis (Table 5); where we made a choice of our own it is documented, and options reproduce the reference tool’s behaviour where we know it. RapidGLM’s default check of the fitted tail changes the results of the SCEPTRE method for pairs whose draws contradict the fit, and takes longer (Results, Accurate small *p*-values); SCEPTRE’s acceptance remains an option. QTL mapping covers the cis effects of single variants, calibrated by permutations, not trans effects or fine-mapping. The engine runs on CPUs; tensorQTL and jaxQTL are designed for GPUs and were compared on CPUs only. The timings come from one workstation, one type of cloud instance and supercomputer nodes of one CPU type, and are compared only within a type of machine. SCEPTRE’s time on a 16-core server is an extrapolation, not a measurement.

#### Outlook

The engine is not specific to these four analyses. A method that fits a count model to each gene and tests a treatment by a Wald, likelihood-ratio or score statistic, with or without resampling, can be built on the C library and optimized in the same way, and the MIT licence lets it be embedded in software of any licence. Two possible directions could make the engine itself faster. One is GPUs, to which the matrix products of permutation scoring are suited, as tensorQTL [25] and jaxQTL [26] show. The other is wider vector instructions: AVX-512 instructions process twice as many numbers at a time as the AVX2 instructions that the kernels use now.

## Conclusions

RapidGLM is an engine for count regression and resampling built for the structure of these models: Newton steps where the likelihood allows them, shared work done once, and vectorized kernels where the time remains. Its implementations of the DESeq2 procedure, edgeR’s quasi-likelihood workflow, the SCEPTRE method and cis-QTL mapping agree with the reference tools to within their own variation and run several times faster, which makes analyses at the scale of whole screens and cohorts practical on one machine. It is also easy to adopt. The R and Python packages run each of the four analyses in one call, and provide drop-in replacements for R’s glm and `MASS::glm.nb` and for the statsmodels GLM class. As a demonstration, passing RapidGLM’s back end as PyDESeq2’s `inference` argument made an otherwise unchanged PyDESeq2 analysis 2.8 times faster on one thread (Table 4). RapidGLM is available from C, Python, R and the command line under the MIT licence.

## Methods

### Clean-room implementation

RapidGLM is distributed under the MIT licence. DESeq2 (LGPL, version 3 or later), edgeR and limma (GPL, version 2 or later), and SCEPTRE and FastQTL (GPL, version 3) are distributed under copyleft licences, which require software derived from their code to be distributed under the same terms. We therefore implemented their methods without reading their source code. Each implementation was written from the method’s publications, from the tool’s documentation, and from the tool’s outputs, with the tool run through its public functions as a black box. Where the publications and the documentation leave a rule open, such as a threshold or the handling of an edge case, the rule was set to reproduce the tool’s outputs; these rules are listed for each analysis, below or in Additional file 1. RapidGLM neither links to these tools nor includes them. The same applies to the other copyleft-licensed software in the comparisons, such as R and its MASS package. Because the source was not read, agreement with each tool is measured, and is reported for each analysis.

Software under permissive licences, which allow code to be read and reused with attribution, was read and is credited. PyDESeq2 [13], an implementation of the DESeq2 procedure in Python, is distributed under the MIT licence, not under DESeq2’s. Some computations of `bulk_de` follow its code and are listed under the DESeq2 procedure, and the Rust binding includes a port of its workflow. The cis-QTL mapper follows the design of jaxQTL (MIT) [26]. glmGamPoi [27] was distributed under the GPL before version 1.25 and is under the MIT licence from that version; only its MIT-licensed code was read, without the blocks that declare a DESeq2 or edgeR origin. The repository’s DEPENDENCIES.md records, for every tool and library, its licence, how it was used and what was read.

#### AI-assisted development

RapidGLM was written with AI-assisted programming under human direction, following the approach we describe for STAR Suite [28]. The comparisons with the reference tools, the binary128 reference fits (Benchmarks) and the comparisons between the interfaces (Engine: Interfaces) also verify the AI agents’ work.

### Engine

The engine fits a GLM to each of many responses, such as the genes of an experiment, and tests its coefficients; the four analyses are built on it. Throughout, *X*′ is the transpose of *X*, hats mark estimates from the fitted model and tildes those from the null model, as in Fig. 1.

### Fit

#### Model

For the observations *i* = 1, …, *n* of one response *y* (the samples or cells of a gene), a GLM specifies the mean, the variance and how the mean depends on the predictors:

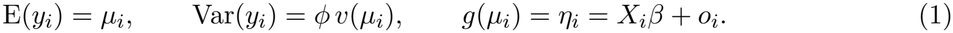

- *X*_*i*_ is row *i* of the design matrix *X*, which has a column for each predictor: the intercept, the treatment *x* and the covariates *z* of Fig. 1a. The engine uses *X* as given; adding an intercept, coding factors and normalizing are left to the caller.
- *β* holds the coefficients, *o*_*i*_ is the offset, *g* the link function, *v* the variance function and *ϕ* a scale, which is 1 except for the Gaussian family.
- The engine has four families. The score residual *r*_*i*_ and the weight *w*_*i*_ of each are used in the fit and are defined below. The NB dispersion is 1*/θ*. Observations may carry prior weights, which are left out of the formulas.

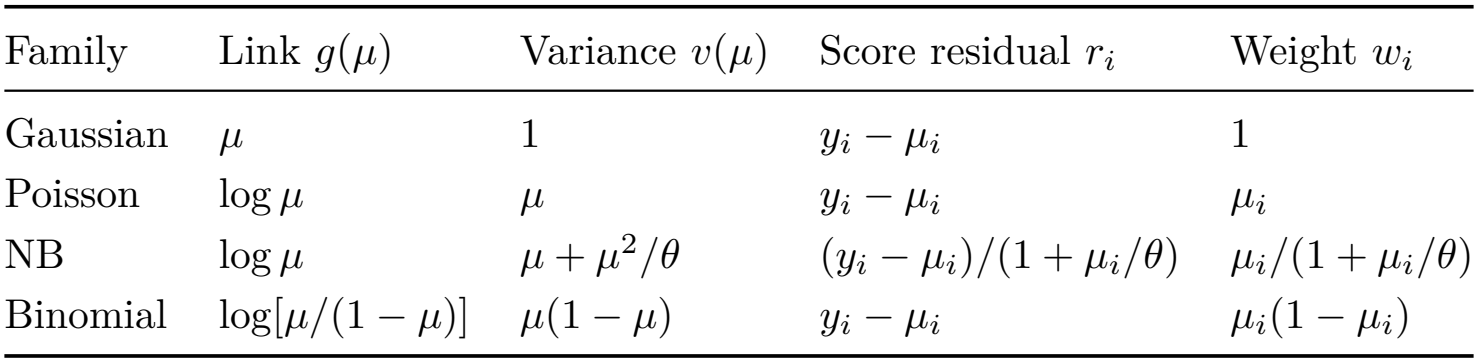

| Family | Link g(μ) | Variance v(μ) | Score residual r<sub>i</sub> | Weight w<sub>i</sub> |
|---|---|---|---|---|
| Gaussian | μ | 1 | y<sub>i</sub> − μ<sub>i</sub> | 1 |
| Poisson | log μ | μ | y<sub>i</sub> − μ<sub>i</sub> | μ<sub>i</sub> |
| NB | log μ | μ + μ<sup>2</sup>/θ | (y<sub>i</sub> − μ<sub>i</sub>)/(1 + μ<sub>i</sub>/θ) | μ<sub>i</sub>/(1 + μ<sub>i</sub>/θ) |
| Binomial | log[μ/(1 − μ)] | μ(1 − μ) | y<sub>i</sub> − μ<sub>i</sub> | μ<sub>i</sub>(1 − μ<sub>i</sub>) |

#### What is maximized

A fit finds the coefficients 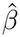 that maximize the log likelihood *l*(*β*). For the NB model at a given *θ*, with *µ*_*i*_ = exp(*X*_*i*_*β* + *o*_*i*_), it is

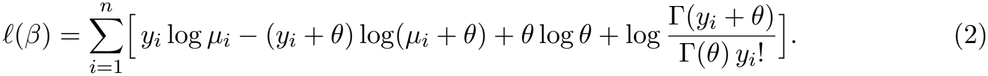

The slope and the curvature of *l* in the coefficients are

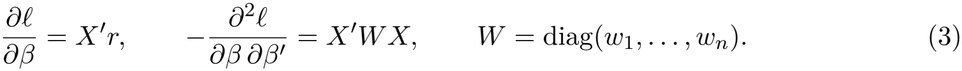

- The score residual is *r*_*i*_ = (*y*_*i*_ − *µ*_*i*_)(*∂µ*_*i*_*/∂η*_*i*_)*/v*(*µ*_*i*_): the distance of an observation from its fitted mean, scaled by the variance.
- The weights of the table, *w*_*i*_ = (*∂µ*_*i*_*/∂η*_*i*_)^2^*/v*(*µ*_*i*_), give the curvature exactly for the Gaussian, Poisson and binomial families. For the NB they give its average over the counts expected under the model (the expected information). The exact curvature at the observed counts (the observed information) has *w*_*i*_ = *θµ*_*i*_(*θ* + *y*_*i*_)*/*(*θ* + *µ*_*i*_)^2^.
- These weights are positive, so *l* is concave in *β*: at a given dispersion it has one maximum and no other peak.
- The deviance *D* is minus twice the log likelihood, up to a constant, so maximizing *l* minimizes *D*.

#### Newton iteration

Each iteration solves a linear system for a step *δ*^(*t*)^ from the current estimate *β*^(*t*)^, with *r*^(*t*)^ and *W* ^(*t*)^ evaluated there, and takes the step:

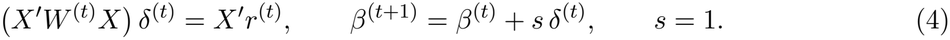

Three settings make the iteration aggressive.

1. *The step length is s* = 1. Every iteration takes the whole Newton step, with no damping, line search or trust region. Only step halving, the first safeguard below, reduces *s*.
2. *The curvature is exact. W* holds the weights of the observed information. Fisher scoring, the usual method for GLMs, uses those of the expected information, which differ for the NB. With the exact curvature, the number of correct digits about doubles with each iteration near the maximum.
3. *No iteration is spent on confirming convergence*. A fit has converged when the relative changes of the deviance and of the coefficients are below a tolerance *ϵ* = 10^−9^:

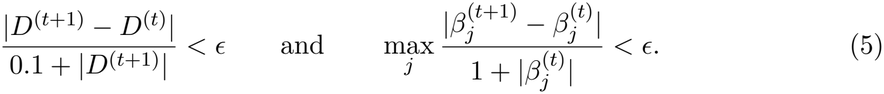

A fitter usually learns this from one more iteration, which changes nothing. The engine predicts that iteration from the factorization of *X*′*W* ^(*t*)^*X* that it already holds. With *u* = *X*′*r*^(*t*+1)^ the slope at the new estimate, the next step would be about *d* = (*X*′*W* ^(*t*)^*X*)^−1^*u*, and it would change the deviance by about *u*′*d*. If both meet the criteria, *d* is added as a final refinement and the fit ends, without another cross product *X*′*WX*, factorization or likelihood.

The count fits start from the least-squares fit of log 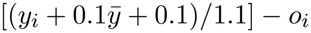 on *X*, where 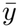 is the mean count.

#### Safeguards

The likelihood is concave, so the whole step rarely fails. The safeguards cover the fits where it does, and the data for which no estimate exists.

- *Step halving*. A step is accepted if the log likelihood is finite and no lower than before: *l*(*β*^(*t*)^ + *sδ*^(*t*)^) ≥ *l*(*β*^(*t*)^) − 10^−12^ (1 + |*l*(*β*^(*t*)^)|). Otherwise *s* is halved, up to 40 times, as in glm2 [29]. Every accepted estimate is therefore an improvement on the last. If no step is accepted, the fit ends with a failure status.
- *Nearly collinear designs*. The system of Eq. 4 is solved by a Cholesky factorization of *X*′*WX* with scaled columns. When the reciprocal condition number of that matrix is below 10^−8^, the step is solved instead by a Householder QR factorization of *W* ^1*/*2^*X*, which is more accurate for such designs.
- *Aliased columns*. Columns that are linear combinations of other columns are detected with the threshold of R’s `glm.`
- *Estimates at infinity*. A coefficient has no finite estimate when, for example, every count of a group is zero. Such coefficients are detected and flagged, and their standard errors are reported as not-a-number.
- *Limits*. A fit that has not converged after 100 iterations is reported as not converged. Every fit returns a status, which the analyses check.
- *Convergence without a confirming iteration* is used only when no weight changed by more than a quarter in the last step, and the final refinement is dropped if it lowers the likelihood.

Additional file 1 (Section S9) gives the linear algebra and these checks.

#### NB dispersion

The fits above take *θ* as given. Where it is estimated, the engine has two estimators.

- *Maximum likelihood*. Newton steps are taken in *β* and log *θ* together, again whole steps with the observed information, from a Poisson fit and a moment estimate of *θ*. The safeguards are step halving, a limit of 2 on the change of log *θ* in one step, the bounds 10^−6^ ≤ *θ* ≤ 10^8^ and, where the likelihood is not concave in log *θ*, a unit step up the slope in place of the Newton step.
- *Cox–Reid adjusted likelihood* [24, 30]. With few samples, maximum likelihood underestimates the dispersion, because the coefficients are estimated from the same samples. The adjustment corrects for this by maximizing 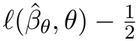 det(*X*′*WX*) over *θ*, where 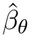 holds the coefficients fitted at *θ* and *W* the weights of the table. The engine alternates Newton steps in log *θ*, with analytic derivatives of the adjustment, and coefficient fits. The DESeq2 procedure and the edgeR workflow use this estimator.

### Resampling

A resampling test estimates a *p*-value by simulation. The treatment is simulated many times, and the statistics of the simulated treatments show how large the statistic gets when the treatment has no effect (Background). Fig. 4 follows one such test.

**Figure 4:**
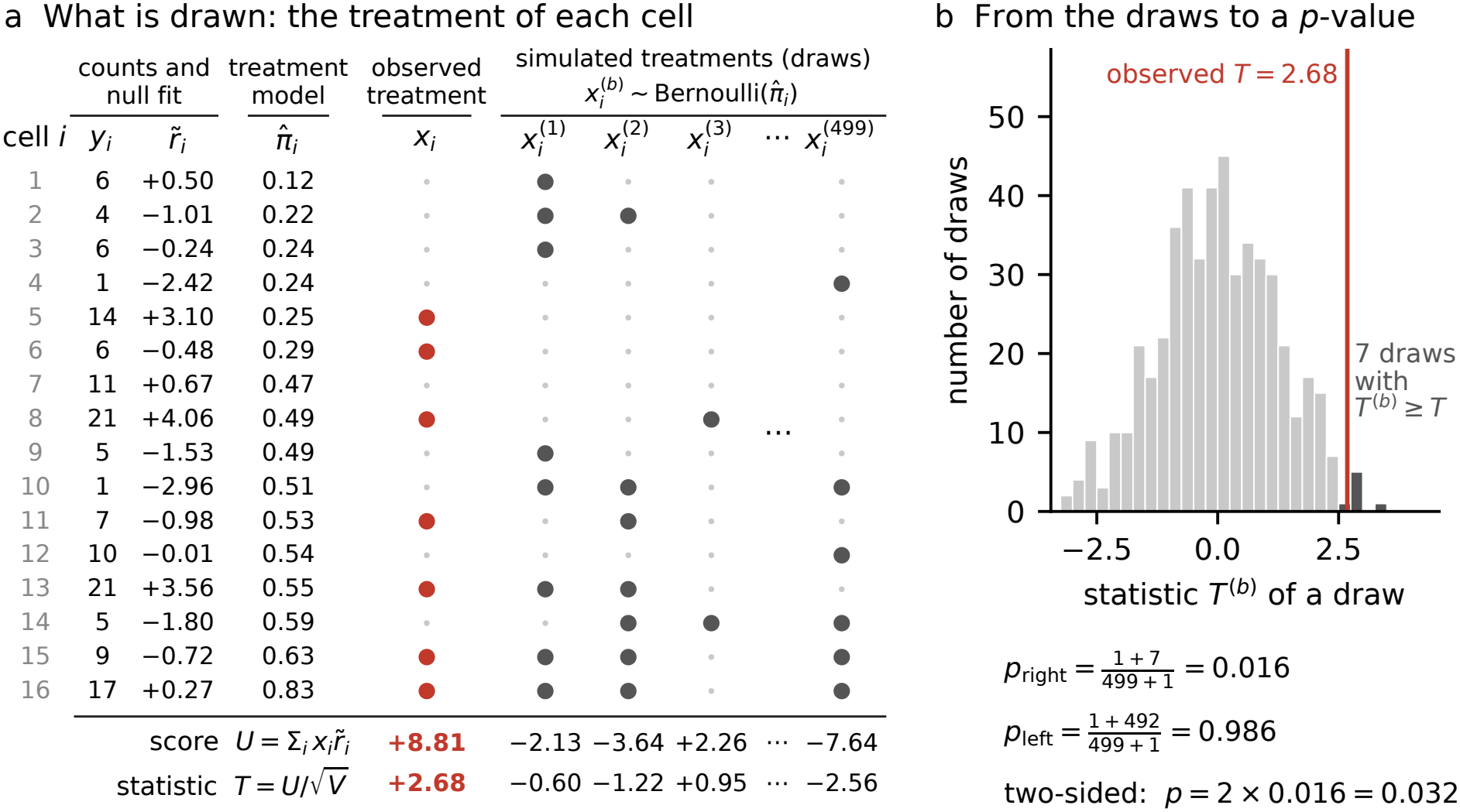
What a resampling test draws, and how the draws give a *p*-value. An illustration with simulated data: one gene in 16 cells (NB counts, one covariate), tested by the conditional randomization test with *B* = 499 draws. (a) Each row is a cell. The counts *y*_*i*_, the score residuals 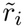 of the null fit and the probabilities of treatment 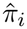 stay fixed; only the treatment is simulated. A filled dot marks a treated cell: the observed treatment in red, and four of the 499 simulated treatments in grey. Under each column are its score *U*, the sum of 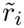 over its treated cells, and its statistic *T*. (b) The statistics of the 499 draws. Seven are at least as large as the observed statistic (dark bars), which gives the *p*-values below the histogram.

#### What is drawn

The steps below are those of the conditional randomization test [8], which SCEPTRE uses at high MOI, for one gene and one target.

1. *Fixed: the counts and the null fit*. The null model of the gene, which has the covariates but not the treatment, is fitted once. It gives each cell’s fitted mean 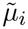, score residual 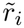 and weight 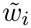.
2. *Fixed: the treatment model*. A logistic regression of the observed treatment *x* on the covariates, fitted once for the target, gives each cell’s probability of being treated, 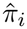.
3. *Drawn: the treatment of every cell*. Draw *b* of *B* simulates 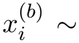 Bernoulli 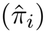: cell *i* is treated with probability 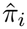 independently of the other cells. The counts, the covariates and the null fit do not change.
4. *Scored*. Each draw is scored against the null fit in the same way as the observed treatment (Scoring, below), which gives its statistic *T* ^(*b*)^.

A simulated treatment depends on the covariates alone, not on the counts, so *T* ^(1)^, …, *T* ^(*B*)^ show how the statistic varies when the treatment has no effect on the gene. A permutation test draws differently: it shuffles the observed values, the treatment among the cells of a pair (SCEPTRE method, at low MOI) or the response among the samples (cis-QTL mapping). Its *p*-value is computed in the same way.

*p***-value**. With *T* the statistic of the observed treatment, the *p*-values are

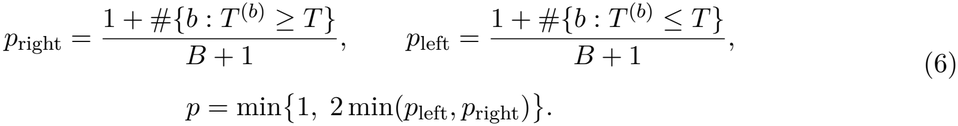

- *p*_right_ tests for an increase of the response, *p*_left_ for a decrease and the two-sided *p* for either.
- The 1 added to each count stands for the observed treatment itself, so a *p*-value is never zero [9].
- In Fig. 4b, 7 of the 499 draws have *T* ^(*b*)^ ≥ *T*, so *p*_right_ = 8*/*500 = 0.016 and *p* = 0.032.
- No *p*-value can be smaller than 1*/*(*B* + 1). Smaller values need more draws or a distribution fitted to the draws (SCEPTRE method, step 5; cis-QTL mapping, step 4).

#### Scoring

The statistic is the score statistic *T*_S_ of Fig. 1b. For a predictor *x* added to a null model whose design is *X*_0_ (the covariates, without the treatment),

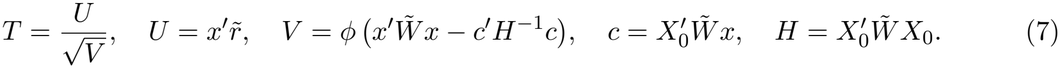

- *U* is the slope of the log likelihood in the coefficient of *x* at the null fit, and *V* is its variance when *x* has no effect.
- 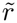 and 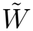 hold the score residuals and the weights of the table, at the null fit.
- The term *c*′*H*^−1^*c* allows for the estimated coefficients of the null model: it removes the part of *x* that the covariates predict.

A simulated treatment is 0 or 1 in each cell, so for draw *b* the products are sums over the set *S*_*b*_ of the cells that it treats (with *ϕ* = 1 for the count models):

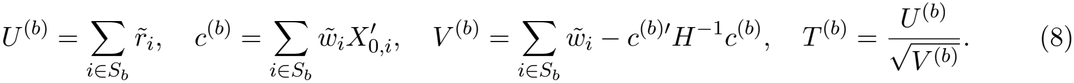

- No draw needs a fit. *H* is factorized once for a null fit and serves every draw of every target tested against the gene, and every variant of a cis-QTL scan.
- The cost of a draw is in proportion to the number of cells it treats, not to the number of cells. At high MOI, few cells carry a guide against any one target.
- The values that a cell adds to these sums (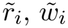 and 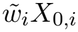) are stored together in one record, so a treated cell costs one read of memory (Table 2).
- A draw with *V* ^(*b*)^ = 0, such as one that treats no cell, is given the statistic 0 and stays among the draws.

### Optimizations

#### General

These apply to every fit and test.

- *Work shared between fits*. Fits of many responses to one design share *X*′*X*, its rank check, the grouping of identical rows and, for sparse designs such as those of factors, each row’s nonzero entries.
- *Grouped designs*. When the rows take exactly as many distinct values as there are columns (one-way and full-factorial layouts), a Newton step needs only per-group sums, an idea from glmGamPoi [27], and the fit starts from the group means.
- *Reuse of the factorization*. The factorization of a final fit gives the standard errors, the leverages (each observation’s influence on its own fitted value) and the Cook’s distances without a refit, and the score statistics of any number of added predictors.
- *Vector instructions (SIMD)*. The inner loops (the cross products *X*′*WX*, the exponentials and logarithms of the fitted means, and the random numbers of the draws) are kernels that process several numbers with one instruction. Each has a path for AVX2 with fused multiply–adds, chosen at run time, and a portable path.
- *Special functions*. The engine has its own exponential, logarithm, digamma and trigamma functions, which return the same bits on every processor. The terms of Eq. 2 that depend on the response alone, log Γ(*y*_*i*_ + *θ*) − log Γ(*θ*) − log *y*_*i*_!, are evaluated by Stirling’s series, with one logarithm per large count and a table for small counts.
- *Threads*. Responses are fitted in parallel, and results are identical at any thread count. On processors without AVX2 the cross products round differently in the last bits.
- *Random numbers*. Draws come from Philox streams [23] keyed by the seed and by what is tested (the pair of a target and a response in a screen, the gene in a cis-QTL scan), so a result does not depend on the thread count or on what else is in the run.

Additional file 1 (Section S9) describes the kernels and gives the accuracy of the special functions.

#### Of each analysis

Each analysis adds optimizations of its own, which are listed in Table 3 and described in its section below.

### Interfaces

Fig. 2 shows the layers.

- *C layer* (`librapidglm_c`). It converts inputs and results only, so its outputs equal those of the C++ calls bit for bit. Its option structures carry their size, so a program built against an older header works with a newer library.
- *Bindings*. The Python, R and Rust bindings call the C layer, and the Python and R packages run each of the four analyses in one call on data in memory. The command-line program is written in C++; it calls the analyses directly and runs them from files.
- *PyDESeq2* [13]. A back end, passed as PyDESeq2’s `inference` argument, replaces PyDESeq2’s model fits, dispersion estimates and Wald tests and leaves the rest unchanged. A PyDESeq2-style interface takes the same kind of calls and runs the whole DESeq2 procedure in the engine.
- *Drop-in fitters* stand in for `stats::glm, MASS::glm.nb` and the statsmodels `GLM` class.
- *Same results*. The Python package’s tables equal the command-line program’s bit for bit. The R package’s equal them for the bulk workflows and for cis-QTL mapping, and agree to 10^−10^ for the SCEPTRE method (Additional file 1, Sections S8, S9 and S14).

### DESeq2 procedure

`bulk_de` applies the engine to differential expression between groups of bulk or pseudobulk RNA-seq samples. It follows the procedure published for DESeq2 [3], where each step and the reasons for it are described; DESeq2 1.48.2 was run as a black box (Clean-room implementation). The procedure fits an NB model to each gene’s counts and tests one coefficient of the design, such as the effect of a treatment. A handful of samples cannot estimate a gene’s dispersion *α* = 1*/θ* well, so the genes share information.

1. *Normalization*. A size factor *s*_*i*_ accounts for the sequencing depth of sample *i* [31], and enters every fit as the offset *o*_*i*_ = log *s*_*i*_.
2. *Dispersions*. Each gene’s dispersion is estimated from its own counts, a trend is fitted to these estimates against the genes’ mean expression, and each estimate is shrunk towards the trend.
3. *Tests*. Each gene is fitted at its shrunken dispersion, and the coefficient is tested by the Wald or the likelihood-ratio test.
4. *Outliers and filtering*. A count that moves its gene’s fit too far, as measured by Cook’s distance [32], is replaced, or the gene is given no *p*-value. Genes of low mean expression are left out of the Benjamini–Hochberg adjustment [33, 34].

Additional file 1 (Section S10) gives every formula and constant. It marks the rules that DESeq2’s paper and documentation leave open, which were set to reproduce DESeq2’s outputs, and the choices that are our own. Three computations follow PyDESeq2’s code [13]: the robust dispersion used for Cook’s distances, the starting dispersion and the iterations of the floored fits.

#### Optimizations

Every step that fits a model uses the engine: its Newton fits, the work shared between genes fitted to one design, its special functions and its threads (Engine: Optimizations). Four further optimizations are specific to this procedure (Table 3).

- *Dispersions by Newton steps*. Both estimates of step 2 maximize, at the gene’s fitted means 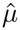,

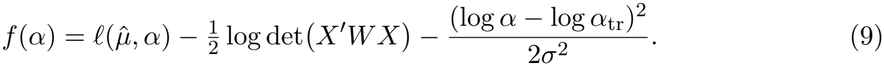 Here *l* is the log likelihood of Eq. 2 and the second term is the Cox–Reid adjustment (Engine: NB dispersion). The third term is the prior that shrinks the estimate towards the trend’s value *α*_tr_, with a variance *σ*^2^ estimated from all genes; the gene’s own estimate is computed without it. The DESeq2 paper describes a backtracking line search in log *α* for both maximizations. `bulk_de` takes Newton steps in log *α*, with the slope and the curvature of all three terms in closed form. With *A* = *X*′*WX* and primes for derivatives in log *α*, those of the adjustment are 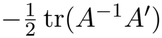 and 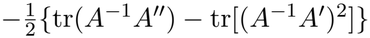.
- *Floored fits only where needed*. DESeq2 documents a lower limit of 0.5 on the fitted counts (`minmu`). A fit with this floor is computed by iteratively reweighted least squares, as in PyDESeq2. Where no fitted count is below the floor, the floor has no effect. Every gene is therefore fitted first by the engine’s Newton iteration (Eq. 4), and only the genes with a fitted count below 0.5, or with a coefficient that has no finite estimate, are refitted with the floor.
- *Outliers without a second pass*. Cook’s distances come from the factorization of each gene’s final fit (Engine: Optimizations), with no further fit. After counts are replaced, only their genes are estimated again, with the trend and the prior kept.
- *One compiled call*. The steps run in one call, with no interpreted code between them, and each pass over the genes runs in parallel; the results are identical on one and eight threads. The fits and the dispersion estimates can also be called on their own, which is how the PyDESeq2 back end replaces those parts of PyDESeq2 (Engine: Interfaces).

### Agreement with DESeq2

The datasets are described in Table 7.

- *Discoveries*, at a false discovery rate of 0.1. The Jaccard indices with DESeq2 were 0.98–1.00 on five validation datasets, with Spearman correlations of the *p*-values of at least 0.999, and 0.97–1.00 on the pseudobulk designs of 20 perturbations of the Replogle pilot. On the 647 thyroid samples RapidGLM made 7,178 discoveries and DESeq2 7,177.
- *False positives*. In 70 mock comparisons of four donors against four other donors of the same condition, where no gene truly differs, RapidGLM rejected 2.37% of the 70,000 tests at *p* ≤ 0.05 and DESeq2 2.39%.

### edgeR quasi-likelihood workflow

`edger_ql` applies the engine to the same question as the DESeq2 procedure, by a different method. It follows the default quasi-likelihood workflow published for edgeR 4 [5, 35], where each step and the reasons for it are described; edgeR 4.6.2 was run as a black box (Clean-room implementation). The workflow fits an NB model to each gene, with one NB dispersion *α* = 1*/θ* for all genes. The variability that this dispersion does not explain is captured by a quasi-dispersion, a multiplier of the variance that is estimated for each gene, and the genes share information through it.

- *Filtering and normalization*. Genes with too few reads are removed, and the library sizes are scaled by TMM factors [36].
- *NB dispersion*. The one dispersion maximizes the Cox–Reid adjusted likelihood (Engine: NB dispersion), summed over the genes of highest average expression.
- *Quasi-dispersions*. Each gene is fitted at a working dispersion derived from that value. Its quasi-dispersion *s*^2^ is estimated from its unit deviances *d*_*i*_, the terms that the samples add to its deviance, *D* = Σ _*i*_ *d*_*i*_. The *s*^2^ are then shrunk towards a trend in average expression by limma’s empirical Bayes method [37], which gives each gene a value 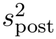.
- *Tests*. A hypothesis on *q* coefficients is tested by 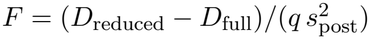, with *D* the deviances of the gene’s fits without and with those coefficients, and the *p*-values are adjusted by the Benjamini–Hochberg method [34].

Additional file 1 (Section S11) gives every formula and constant. It marks one constant that edgeR’s documentation leaves open, which was set to reproduce edgeR’s outputs (the value at which the search for the prior degrees of freedom of the shrinkage ends), and one choice that is our own (the moments of the unit deviance, below).

#### Optimizations

Every fit uses the engine: its Newton iteration with the observed information (Eq. 4), its kernels and its threads (Engine: Optimizations). Three further optimizations are specific to this workflow (Table 3).

- *Fits that start from the fit before*. The workflow fits each gene several times: at each trial value of the dispersion search of step 2, at the working dispersion of step 3 and again when that dispersion is rescaled, for the reduced design of the test, and for the fold changes, which are estimated from the counts with a small count added. Each fit starts from the coefficients of the fit before it.
- *Convergence without another factorization*. These fits end, in place of the test of Eq. 5, when the Newton decrement *λ*^2^, which is about the deviance that the fit could still gain, is below a threshold:

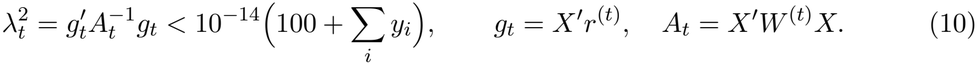

Computed directly, 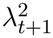 needs a new cross product *A*_*t*+1_ and its factorization, only to confirm that the fit has ended. A change of Δ in a linear predictor changes the weight of its sample by a factor of at most *e*^Δ^, so after a whole Newton step

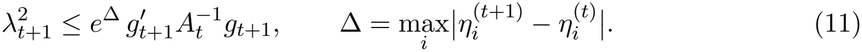 The right-hand side uses the factorization already in hand. When it is below the threshold the fit has converged; otherwise the next iteration computes 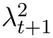 itself.
- *Exact moments of the unit deviance, from a table*. The quasi-dispersion of step 3 is

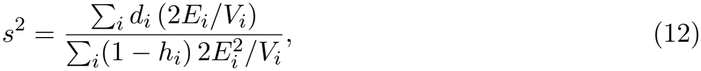

where *h*_*i*_ is the leverage of sample *i*, and *E*_*i*_ and *V*_*i*_ are the mean and the variance of its unit deviance under the NB model. `edger_ql` computes these moments exactly, as sums over the NB distribution, once for each dispersion on a grid of 64 means per decade; a sample then costs one interpolation in the table. edgeR uses an approximation that its documentation does not specify, from which the exact values differ by up to about 10^−3^ for dispersions up to 0.3.

#### Agreement with edgeR

The datasets are described in Table 7; discoveries are at a false discovery rate of 0.1.

- *Airway, the Kang panel and three simulations* (five of eight datasets). The discoveries equal edgeR’s, and the *p*-values agree to within 0.006 in log_10_. On all eight datasets the genes kept by the filter, the TMM factors (to 5.1 × 10^−15^) and the average log CPM agree to rounding.
- *GTEx thyroid sets* of 32, 128 and 647 samples. For 22, 7 and 9 of their genes, the deviance that edgeR’s `glmFit` reports differs from the deviance of its own fitted values, or `glmFit` flags the fit as failed; edgeR’s *F* statistics for those genes use the reported value. Without edgeR’s implementation the reason cannot be checked. The check uses edgeR’s public functions (Additional file 1, Section S11). Over the other genes the discoveries differ by at most 7 (7,480 against 7,477 for 647 samples).

### SCEPTRE method

`sceptre` applies the engine to single-cell CRISPR screens: it tests whether perturbing a target changes the counts of a response, a gene or a peak. It follows the method published for SCEPTRE [10, 11], where each step and the reasons for it are described; SCEPTRE 0.99.0 was run as a black box (Clean-room implementation). For each pair of a target and a response, the method computes the score statistic of the treatment *x*, which is 1 for the cells that carry a guide against the target, and estimates its *p*-value by simulating the treatment many times (Fig. 1b).

- *Quality control*. Cells with extreme total counts, numbers of detected features or mitochondrial fractions are removed. A pair is tested when at least seven treated cells and seven control cells have a nonzero count of its response.
- *Null model*. The response is fitted to the covariates without the treatment: a Poisson regression, then the maximum-likelihood NB dispersion at its fitted means. At high MOI the controls of a target are all the cells without its guides, and one null fit per response serves all its pairs.

At low MOI the controls are the cells with non-targeting guides, and the null model is fitted for each pair, on its treated and control cells.

- *Statistic*. The NB score statistic of Eq. 7 for the treatment *x*.
- *Resampling*. At high MOI the treatment is simulated by the conditional randomization test (Engine: Resampling; Fig. 4). At low MOI the treatment labels are permuted among the pair’s treated and control cells. The *p*-values are those of Eq. 6.
- *Small p-values*. The draws are made in stages: 499, 4,999 and 19,999 in all. The *p*-value of the first 499 draws is final unless it is at most 0.02, which settles most pairs. For the others, the *p*-value is read from a skew-normal distribution [38] fitted to the statistics of the draws, and can be smaller than 1*/*(*B* + 1).
- *Multiplicity*. The *p*-values are adjusted by the Benjamini–Hochberg method [34] within each kind of pair: discovery pairs, negative controls and positive controls.

Additional file 1 (Section S12) gives the full specification. It marks the rules that SCEPTRE’s papers and documentation leave open, which were inferred from SCEPTRE’s outputs: the first stage and its threshold, the test by which a fitted distribution is accepted (`fit_check=ks,` used for the comparisons with SCEPTRE below) and the family of pairs over which the *p*-values are adjusted. Two choices are our own: the cells of a low-MOI null fit, and the default acceptance of a fitted distribution (`fit_check=strict`). The default also requires the draws in the tail on the statistic’s side to be no heavier than the fit says, sends a *p*-value too small for its stage to check on to the next stage, and confirms the smallest on 199,999 draws (Results, Accurate small *p*-values).

#### Optimizations

The null fits, the treatment models, the scores of the draws (Eq. 8) and the random streams are the engine’s (Engine: Resampling and Optimizations). In a whole screen most of the time goes to the draws of the conditional randomization test, and their limit is reading the values of the treated cells, which are scattered through memory. The optimizations below are specific to this method (Table 3). Unless stated, none changes a result: every draw keeps its own random numbers and adds its cells in the same order, and the test suite compares every draw and statistic with a plain implementation that makes one draw at a time.

- *Quality control first*. Whether a pair is tested is decided before any model is fitted, from the target’s cells and the response’s nonzero cells, so a pair that fails costs nothing more. Targets and responses can be named without listing their pairs.
- *Work shared between pairs*. At high MOI, one null fit serves all the pairs of a response, and one treatment model and one sampler serve all the pairs of a target. Only the discoveries are refitted with the treatment in the model, to estimate its effect, and each such fit starts from the null fit.
- *A sampler whose cost follows the treated cells*. A simulated treatment is not drawn cell by cell. Each cell is given an interval of length *τ*_*i*_, the intervals are laid end to end, and the points of a Poisson process of unit rate are placed along them; the cells whose intervals hold a point are treated. With

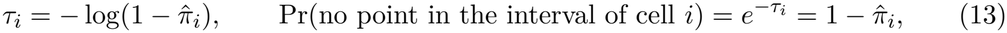

and disjoint intervals independent, the treatments are the independent Bernoulli 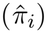 draws of the test, at a cost in proportion to the number of treated cells, not the number of cells. The gaps between the points come from ARS, the AES-based generator of Salmon et al. [23]. Processors without AES instructions use a thinning sampler [39], which draws from the same distribution with different random numbers, so its results differ as those of two seeds do; the run record names the sampler.
- *Draws in batches* (the three steps of Table 2). Draws are generated and scored 256 at a time, in blocks of cells, so that each block of a response’s values is read once for all the draws of a batch. The threads work together on a few pairs of one response, whose values stay in the cache that they share. The blocks of a Philox stream are computed eight at a time with vector instructions.
- *Batch scoring*, the default at high MOI from RapidGLM 1.6.0. Each thread takes one response and scores the next draws of all its pairs in one pass over the response’s values. It holds up to 8,192 draws of each of the response’s pairs at once, so its memory grows with the number of targets in a tile, one of the blocks of targets and responses into which a run is divided. With all 3,837 targets of the Gasperini test set (its 3,776 targets, the 51 non-targeting targets and those of the positive controls) in every tile, the default run on the whole test set exceeded a 128 GiB instance; with 64 targets per tile it ran in 33.9 GiB and gave the same table.
- *Null fits started from the response’s fit*, at low MOI, from RapidGLM 1.6.0. The null fits of a response’s pairs differ only in the cells of their targets. Each starts from *β*_ref_, the response’s fit on all cells, updated by one scoring step on the pair’s cells, the control cells *C* and the target’s cells *S*:

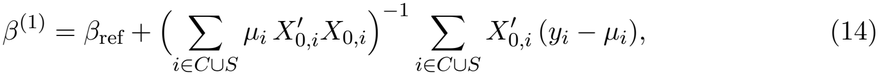

with *µ*_*i*_ the fitted means of the response’s fit. Both sums add up cell by cell: the part over *C* is computed once per response, and each pair adds the cells of its target. This one-step estimate [40] is used only as a start [41]; Newton iterations on the pair’s cells refine it to the convergence criteria of a fit from scratch, which is the fallback. The refined fit equals a fit from scratch to within those criteria, not bit for bit (Results, The same results).
- *Runs that can be divided*. A pair’s draws depend only on the seed, the response and the target (Engine: Optimizations), so the parts of a run that is divided into tiles, checkpoints or machines merge into the table of an undivided run. Optionally (`reuse_draws`), the draws of a target are shared by all its pairs, as in SCEPTRE’s resampling scheme (Table 6); the errors of a target’s *p*-values are then correlated.
- *An optional early end to the first stage* (`sequential_screen`). The first stage ends as soon as enough draws are at least as extreme as the statistic that its *p*-value must exceed the threshold. The pairs that go on to the later stages, and their results, are unchanged; a pair that ends early reports a sequential *p*-value [42].

#### Agreement with SCEPTRE

The two tools use different random draws, so pairs near the discovery threshold can differ, and the yardstick is the agreement between two runs of RapidGLM with different seeds. Discoveries are at a false discovery rate of 0.1 (datasets in Table 7).

- *SCEPTRE’s example panels*, at low and at high MOI. The statistics agreed with SCEPTRE’s to within 2.4 × 10^−7^, and the discoveries overlapped with mean Jaccard indices of 0.86–0.96, against 0.85–0.95 between two RapidGLM seeds.
- *Gasperini benchmark panel*, on all 207,324 cells. RapidGLM made 98 discoveries and SCEPTRE 101 (Jaccard index 0.90). Over six RapidGLM seeds the index was 0.90–0.96, against 0.90–0.97 between the RapidGLM runs.
- *Replogle pilot*, at low MOI (13,618 cells; the 165,034 pairs that both tools tested). RapidGLM made 10,982 discoveries and SCEPTRE 10,972 (Jaccard index 0.949; Spearman correlation of the *p*-values 0.989).

### cis-QTL mapping

qtl applies the engine to cis-eQTL mapping: it tests each gene against the genetic variants near it and reports, for the gene, the *p*-value of its strongest association, corrected by permutations for the number of variants tested. It follows the design of jaxQTL (MIT) [26], whose permutation scheme and Beta approximation follow FastQTL [12]; these papers describe each step and the reasons for it. FastQTL was run as a black box (Clean-room implementation).

1. *Null model*. Each gene is fitted to the covariates alone: a Gaussian model for normalized expression, or a Poisson or NB model for counts, with the log library size as offset.
2. *Scores*. Every variant within a window of the gene, 1 Mb by default on either side of its transcription start site, is tested by the score statistic *T* of Eq. 7, with the variant’s dosage as the added predictor *x*. The lead variant is the one with the largest |*T* |.
3. *Permutations*. Each of *B* permutations (1,000 by default) shuffles the gene’s response, with its offset, among the samples, refits the null model and keeps the largest |*T* | over the gene’s variants. The *B* maxima show how large the strongest statistic is when no variant has an effect.
4. *Gene-level p-value*. The share of the maxima that are at least as large as the observed one gives a *p*-value as in Eq. 6, which cannot be smaller than 1*/*(*B* + 1). A Beta distribution fitted to the maxima, in jaxQTL’s form, removes that limit.

Additional file 1 (Section S13) gives the full specification.

#### Optimizations

The null fits, the score statistic and the random streams are the engine’s (Engine: Resampling and Optimizations). Four further optimizations are specific to this analysis (Table 3).

- *All the variants of a gene in one pass*. With the dosages of the gene’s variants as the columns of a matrix *G*, Eq. 7 gives the statistics of all of them as

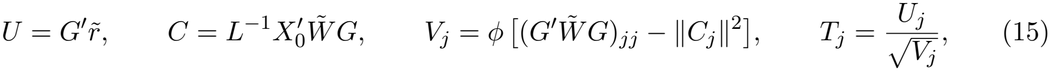

where *LL*′ = *H* is the factorization of the null fit and *C*_*j*_ is column *j* of *C*. The products with *G* are computed in one pass of the engine’s cross-product kernel. *C* and the term in brackets depend on the weights and not on the response, so where the weights do not change, as in a Gaussian model, they are computed once per gene and serve every permutation.
- *Gaussian models: 512 permutations at a time, in single precision*. For a Gaussian model, refitting the null model to a shuffled response is a projection. The covariates are removed from the dosages once per gene, *G*_⊥_ = *G* − *QQ*′*G* with *Q* an orthonormal basis of the covariates, and the scores of 512 shuffled responses, the columns of *Y*_*π*_, are then one matrix product:

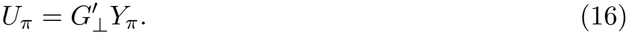

By default the product is computed in single precision, after each shuffled response has been centred and scaled in double precision, and each permutation’s maximum is taken as the product is formed, so the scores are never stored. The observed statistics, the Beta fit and the *p*-values are computed in double precision.

- *Count models: eight permutations refitted and scored together*. For a Poisson or NB model the weights change with every shuffle, so each permutation needs a new null fit and new products with *G*. Eight permutations are refitted in step: each follows the engine’s iteration on its own values, with the sums over the samples of all eight taken as one matrix product, and a fit that needs a safeguard is refitted alone. Their scores are then computed in one blocked product with *G*, which gives each permutation the statistics of scoring it alone, bit for bit. Only the order of the sums in the refits differs from that of separate refits (Results, The same results).
- *Shuffles and storage*. The random numbers of a shuffle are computed eight Philox blocks at a time, and the bounded integers that it needs are taken from them without a division [43]. Genotypes are kept as PLINK’s two-bit codes and decoded gene by gene, and each thread reuses its buffers from gene to gene.

#### Agreement with FastQTL, tensorQTL and jaxQTL

The comparison is on chromosome 22 of Geuvadis (Table 7), with the same 42 covariates and a window of 1 Mb for every tool (Additional file 1, Section S5). Each tool’s gene-level *p*-values are its Beta-approximated values, and eGenes are the genes with Storey’s *q <* 0.05 [44], computed with the `qvalue` package (version 2.40.0, defaults).

- *Gaussian model, 10,000 permutations*. RapidGLM found 267 eGenes, FastQTL 269 and tensorQTL 266 (Jaccard indices with RapidGLM 0.993 and 0.996), and RapidGLM reported FastQTL’s lead variant for 100% of genes. With its adaptive scheme, which stops a gene’s permutations early, after between 1,000 and 10,000, FastQTL found the same 267 eGenes as RapidGLM.
- *NB model, 1,000 permutations*. RapidGLM found 168 eGenes and jaxQTL’s NB score test 170 (Jaccard index 0.988; Spearman correlation of the gene-level *p*-values 0.9996).

### Benchmarks

#### Version and receipts

The version benchmarked is RapidGLM 1.6.0 (tag v1.6.0, commit 9c22070; measured at development commit 4505ef6, whose library source is identical). Unless stated below, every benchmark was measured with it in one run on one workstation, and every number comes from a validation receipt in the repository that records the version and commit measured. Additional file 1 (Section S14) gives the protocol in full.

#### Release 1.6.1

The current release is RapidGLM 1.6.1 (commit c9a061e). It adds two things to 1.6.0: cis-QTL mapping can be called through the C layer and from the R and Python packages, as the other three analyses can, and the repository includes the benchmark program of Table 2. Neither changes the calculations, so the benchmarks were not repeated: on the test inputs of the four analyses, the command-line programs of the two releases wrote identical result tables.

### Machines

- *Workstation*. AMD Ryzen Threadripper PRO 5965WX (24 cores, 125 GiB of RAM): every benchmark not named below.
- *Supercomputer*. Exclusive nodes of Bridges-2 [45] with two 64-core AMD EPYC 7742 processors and 512 or 256 GB of memory: SCEPTRE and RapidGLM on the Gasperini screen.
- *Cloud*. AWS m8id.8xlarge instances (16 cores of an Intel Xeon 6975P-C, 128 GiB of RAM): RapidGLM on every pair of the Gasperini test set.

### Timing rules

- *Time*. Wall-clock seconds, at one thread per physical core, with the cores’ second hardware threads idle.
- *Quiet host*. On the workstation, a command started only when the host was quiet, and was rejected and repeated if other activity took more than 5% of its cores.
- *Repeats*. The tools compared ran in rotation, one to three times each, and the minimum time is reported.

#### Reference tools

All ran only through their public interfaces.

- *DESeq2 procedure*. DESeq2 1.48.2 (Bioconductor 3.21), with BiocParallel on more than one thread; PyDESeq2 0.5.2.
- *edgeR workflow*. edgeR 4.6.2.
- *SCEPTRE method*. SCEPTRE 0.99.0 (GitHub commit 21f9ea0), with its `parallel` option on more than one thread.
- *cis-QTL mapping*. FastQTL (GTEx fork, commit 59bcdc0); tensorQTL 1.0.10 and jaxQTL 0.3.1, on CPUs only.
- *Engine*. R 4.5.1 (`glm, MASS::glm.nb`), glmGamPoi 1.25.1, fastglm 0.1.2, speedglm 0.3-5, statsmodels 0.14.4, scikit-learn 1.7.2, glum 3.4.1 and Ceres Solver 2.2.

### What each table times

- *Table 1.* Fits of data in memory, on one core, with each tool’s default settings: the minimum over up to seven repeats.
- *Table 2.* The null draws only, run in each earlier form by a program in the repository (`scripts/benchmark_crt_steps.py`). These runs did not wait for a quiet host.
- *Table 4.* For the command-line program, the whole process. For the R and Python packages, the calls on data in memory: `DESeq()`and `results()` for DESeq2; building the data set, deseq2() and the summary of the test for PyDESeq2.
- *Table 5.* RapidGLM ran through its R package for the DESeq2 procedure and the Gasperini panel, and from the command line for the other analyses. FastQTL was timed with RapidGLM in a later job: twelve chunks of the genes as twelve processes, a fixed 10,000 permutations, and the time of the whole command.
- *Table 6.* SCEPTRE and RapidGLM ran on the Gasperini screen with the same cells, covariates and seed.

### Definitions

- *Accuracy*. The error against a maximum-likelihood fit computed by Newton’s method in IEEE binary128 arithmetic, in standard errors of that fit. The 39 identifiable problems include badly scaled, nearly collinear and aliased designs, zero weights, large offsets, rare events and extreme dispersions (Additional file 1, Section S1).
- *Memory*. The peak of the summed proportional set size of a run’s process tree, which counts the memory that SCEPTRE’s forked workers share once.
- *SCEPTRE’s time on a 16-core server*. An extrapolation, not a measurement:

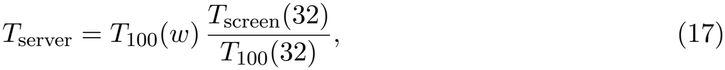

### Datasets

All datasets are public (Table 7). Scripts in the repository’s scripts directory prepare them from their published files (`prepare_benchmark_data.py, validate_bulk.py, prepare_crt_panels.py` and `prepare_replogle.py;` for Geuvadis, `eqtl_prep_expression.py, eqtl_prep_genotypes.sh` and `eqtl_make_covariates.py,` described in `docs/GEUVADIS.md`).

## Availability and requirements

**Project name:** RapidGLM

**Project home page:** https://github.com/morphic-bio/rapidglm

**Operating system(s):** Linux on x86-64; ARM64 builds tested under emulation

**Programming language:** C++17, with C, Python and R interfaces

**Other requirements:** CMake, Boost (headers), LAPACKE and a BLAS, OpenMP; R with cpp11 for the R package; Python 3 with NumPy for the Python package

**License:** MIT

**Any restrictions to use by non-academics:** none

## Declarations

## Ethics approval and consent to participate

Not applicable.

## Consent for publication

Not applicable.

## Availability of data and materials

Source, documentation, the Dockerfile and every validation receipt are in the public repository, https://github.com/morphic-bio/rapidglm. All benchmark inputs are public: airway (Bioconductor), Kang et al. (ExperimentHub), GTEx v8 thyroid (recount3), SCEPTRE’s example data, the Gasperini et al. screen (GEO GSE120861), the Replogle et al. screen (figshare) and Geuvadis with 1000 Genomes genotypes (BioStudies E-GEUV-1), with accessions and preparation scripts in the repository.

## Authors’ contributions

L.H.H. conceived the project, wrote the software, ran the analyses and wrote the manuscript. K.Y.Y. supervised the project and revised the manuscript. Both authors read and approved the final manuscript.

## Acknowledgements

This work used Bridges-2 at the Pittsburgh Supercomputing Center [45] through allocation BIO230124 from the Advanced Cyberinfrastructure Coordination Ecosystem: Services & Support (ACCESS) program [51], which is supported by U.S. National Science Foundation grants #2138259, #2138286, #2138307, #2137603 and #2138296.

## Additional files

**Additional file 1**. Supplementary methods and results (PDF): engine accuracy and speed; the solver comparison; the DESeq2 procedure, including Cook’s distances; edgeR’s quasi-likelihood workflow; cis-QTL mapping; the SCEPTRE method, including the tail counts against the skew-normal fit and the scaling of the conditional randomization test; the applications by interface; interfaces and reproducibility; and, in Sections S9–S14, the full specifications of the engine and of each analysis that Methods summarizes, and the benchmark protocol.

