## Supplemental Material for "RapidGLM: genome-scale differential expression, CRISPR screen and eQTL analysis"

### Additional file 1: Supplementary methods and results

Ling-Hong Hung    Ka Yee Yeung

Sections S1–S8 give the measurements behind the Results of the main text, and Sections S9–S14 the full specifications that its Methods summarize. Every number comes from a validation receipt (JSON) in the RapidGLM repository, named beside it, which records the version and commit measured. Unless the text names another version, the numbers are those of RapidGLM 1.6.0 (tag v1.6.0, commit 9c22070; measured at development commit 4505ef6, whose library source is identical), and the timings are wall-clock seconds on the benchmark workstation (AMD Ryzen Threadripper PRO 5965WX), with one thread per physical core, under the protocol of Section S14. Release 1.6.1 adds cis-QTL mapping to the C layer and to the R and Python packages, and the benchmark program of Table 2 of the main text; its calculations and results are those of 1.6.0 (main text, Methods). The datasets are described in the main text (Methods, Table 7). Reference tools were run only through their public interfaces:

- DESeq2 1.48.2 (Bioconductor 3.21), PyDESeq2 0.5.2 and edgeR 4.6.2;
- R 4.5.1 (`glm`, `MASS::glm.nb`), fastglm 0.1.2 and speedglm 0.3-5; statsmodels 0.14.4, scikit-learn 1.7.2 and glum 3.4.1; Ceres Solver 2.2;
- SCEPTRE 0.99.0, the development version on GitHub (`https://github.com/Katsevich-Lab/sceptre`, commit 21f9ea0 of 2026-08-31); its latest release, v0.10.3, is an earlier version;
- glmGamPoi 1.25.1, the Bioconductor development version (`https://github.com/const-ae/glmGamPoi`). Its Bioconductor 3.21 release, 1.20.0, an earlier version that ships with DESeq2 1.48.2, took the same time to within 0.6% in a separate comparison (Section S7);
- FastQTL (the GTEx fork, commit 59bcdc0), tensorQTL 1.0.10, jaxQTL 0.3.1 and qvalue 2.40.0.

#### S1 Engine accuracy and speed

**Accuracy.** The problems have $n = 2{,}000$ observations and Gaussian, Poisson, binomial and NB responses with estimated dispersion. They cover:

- baseline designs, and columns scaled by $10^{-6}$ to $10^{6}$;
- uncentered covariates, near-collinearity and exact aliasing;
- prior weights including zeros, large offsets, and sparse and large counts;
- rare events, extreme dispersions and all-zero groups.

Reference fits are maximum-likelihood solutions computed by Newton's method in IEEE binary128. Errors are in reference standard errors (SE). The work that the engine shares or skips (main text, Results) was in place for these measurements. Receipt: `validation/GLM_COMPARISON.json`.

**Drop-in fitters.** Through R's `rapidglm::glm_fit` and Python's statsmodels-compatible `GLM`, single fits of the same problems (18 for each) were a median of 1.7 (1.0–2.2) and 3.1 (1.8–17.9) times faster than `glm.fit` and statsmodels, and `rapidglm::glm_nb` was 3.7 (2.9–4.8) times faster than `MASS::glm.nb` on the 3 NB problems. Both carry their host's per-call overhead, and the R method leaves Gaussian fits, a single weighted least-squares solve, to `stats::glm.fit`, which is why its range starts at 1.0. Receipt: `validation/GLM_COMPARISON.json`.

Table S1: Accuracy on identifiable problems (RapidGLM with R-compatible aliasing). Failures: tool errors, missing identifiable coefficients, or coefficient errors above $10^{-3}$ reference SE (minimum-norm answers to aliased designs fail by this test).

| **Fitter** | **Problems** | **Failures** | **Median error (SE)** | **Max error (SE)** | **Max SE error** |
|---|---|---|---|---|---|
| RapidGLM | 39 | 0 | $3.3 \times 10^{-15}$ | $1.7 \times 10^{-6}$ | $2.1 \times 10^{-8}$ |
| R `glm.fit` | 26 | 0 | $1.8 \times 10^{-10}$ | $3.6 \times 10^{-6}$ | $1.9 \times 10^{-5}$ |
| R `glm` | 26 | 0 | $1.8 \times 10^{-10}$ | $3.6 \times 10^{-6}$ | $1.9 \times 10^{-5}$ |
| fastglm | 26 | 3 | $2.2 \times 10^{-10}$ | $1.1 \times 10^{-6}$ | $1.1 \times 10^{-1}$ |
| speedglm | 26 | 12 | $5.3 \times 10^{-12}$ | $3.4 \times 10^{-8}$ | $1.9 \times 10^{-5}$ |
| statsmodels GLM | 26 | 4 | $9.3 \times 10^{-12}$ | $7.4 \times 10^{-4}$ | $1.1 \times 10^{-1}$ |
| glum (defaults) | 26 | 8 | $1.6 \times 10^{-6}$ | $3.4 \times 10^{-4}$ | $6.2 \times 10^{-1}$ |
| glum (tight) | 26 | 6 | $2.8 \times 10^{-12}$ | $2.9 \times 10^{-4}$ | $6.2 \times 10^{-1}$ |
| scikit-learn (defaults) | 26 | 13 | $4.6 \times 10^{-7}$ | $3.5 \times 10^{-4}$ | – |
| scikit-learn (tight) | 26 | 11 | $3.8 \times 10^{-12}$ | $2.6 \times 10^{-7}$ | – |
| RapidGLM `glm_nb` (R) | 13 | 0 | $1.9 \times 10^{-14}$ | $1.3 \times 10^{-6}$ | $1.6 \times 10^{-6}$ |
| R `MASS::glm.nb` | 13 | 0 | $3.1 \times 10^{-9}$ | $1.5 \times 10^{-4}$ | $1.4 \times 10^{-4}$ |
| statsmodels NB | 13 | 8 | $1.7 \times 10^{-4}$ | $3.3 \times 10^{-4}$ | $1.9 \times 10^{-2}$ |

Table S2: Single-fit time relative to RapidGLM's C++ engine (median over problems with $n \geq 10^5$; one thread). R links the reference BLAS.

| **Fitter** | **Poisson** | **Binomial** | **Gaussian** |
|---|---|---|---|
| R `glm.fit` | 6.6 | 5.6 | 6.9 |
| R `glm` | 7.7 | 6.6 | 8.6 |
| statsmodels | 5.6 | 5.6 | 11.1 |
| fastglm | 2.7 | 2.5 | 2.8 |
| speedglm | 3.7 | 3.2 | 2.6 |
| glum (tight) | 3.9 | 1.4 | 1.9 |
| scikit-learn (tight) | 1.1 | 0.9 | 1.4 |

Table S3: Many responses with one design: 1,000 count responses on 5,000 observations, five design columns. RapidGLM's `fit_batch` runs across threads; the other rows loop over single fits on one thread (statsmodels NB timed on 200 responses and scaled to 1,000). Receipt: `validation/ENGINE_BATCH.json`.

| **Tool** | **Family** | **Threads** | **Seconds** |
|---|---|---|---|
| rapidglm fit_batch | Poisson | 1 | 0.72 |
| rapidglm fit_batch | Poisson | 4 | 0.18 |
| rapidglm fit_batch | Poisson | 16 | 0.06 |
| R glm.fit | Poisson | 1 | 4.17 |
| R rapidglm::glm_fit | Poisson | 1 | 2.07 |
| Python statsmodels | Poisson | 1 | 3.88 |
| Python rapidglm.fit | Poisson | 1 | 0.77 |
| rapidglm fit_batch | NB | 1 | 2.22 |
| rapidglm fit_batch | NB | 4 | 0.56 |
| rapidglm fit_batch | NB | 16 | 0.15 |
| R glm.nb | NB | 1 | 103.27 |
| R rapidglm::glm_nb | NB | 1 | 8.16 |
| R glmGamPoi::glm_gp | NB | 1 | 4.44 |
| Python statsmodels | NB | 1 | 18.89 |
| Python rapidglm.fit | NB | 1 | 2.34 |

**Grouped designs.** When the design's rows take exactly as many distinct values as it has columns (intercept-only, one-way and full-factorial layouts), the linear predictor is constant within each group of equal rows. A step then needs only per-group sums and the inverse of the matrix of distinct rows, with no cross product or factorization. The idea of solving such designs group by group comes from glmGamPoi [1].

**Quasi-likelihood tests of the general engine.** The quasi-likelihood $F$ test [2] of the engine's hypothesis tests follows the recipe of glmGamPoi [1]; the edgeR workflow has its own (main text, Methods). Each response gets a Cox–Reid adjusted dispersion; a trend is fitted to these against the mean by a weighted running median; the full and reduced models are refitted at the trended dispersion; and the quasi-likelihood dispersions are shrunk toward a fitted $F$ prior [3], whose scale follows a spline in the mean when there are at least 100 responses.

**What was read of glmGamPoi.** Besides its paper and documentation, only its MIT-licensed code was read (development commit 95cffb79, version 1.25.1); blocks there that declare a DESeq2 or edgeR origin were skipped. RapidGLM's code for both parts was written independently, and glmGamPoi otherwise ran only as a black box (`DEPENDENCIES.md`).

#### S2 Solvers against Ceres Solver

Identical problems were fitted with RapidGLM and with Ceres Solver 2.2 [4]:

- *Fixed dispersion:* NB fits with log size-factor offsets, at dispersions from `bulk_de`. Ceres used Levenberg–Marquardt and dogleg trust regions on deviance residuals $\operatorname{sign}(y-\mu)\sqrt{d_i}$, whose sum of squares is the deviance, and L-BFGS on the negative log likelihood.
- *Maximum-likelihood dispersion:* coefficients and $\log\theta$ jointly. Ceres used BFGS and L-BFGS.
- *Tolerances:* each Ceres method ran at its default tolerances and at tight ones (function and gradient $10^{-12}$, parameter $10^{-10}$).

Every answer is scored by one numerically stable log likelihood. A gene is "worse" when its log likelihood falls more than $10^{-6}$ (relative) short of the best answer any method found for it. RapidGLM is timed two ways: sharing among the genes the work that depends only on the design (computed once and not timed), as its batch and bulk interfaces do, and by a plain `fit` of each gene on its own. The estimates are the same.

Harness: `benchmarks/solvers`. Receipt: `validation/SOLVERS.json`.

Table S4: Solver comparison (one thread).

| **Data** | **Dispersion** | **Method** | **Seconds** | **Genes worse** | **Largest shortfall** |
|---|---|---|---|---|---|
| airway | fixed | RapidGLM Newton | 0.18 | 0 | $1.4\times10^{-9}$ |
| airway | fixed | RapidGLM Newton, plain `fit` | 0.19 | 0 | $1.4\times10^{-9}$ |
| airway | fixed | Levenberg–Marquardt (default) | 0.25 | 0 | $3.4\times10^{-6}$ |
| airway | fixed | Dogleg (default) | 0.26 | 0 | $3.4\times10^{-6}$ |
| airway | fixed | L-BFGS (default) | 0.37 | 9,323 | $5.7\times10^{1}$ |
| airway | fixed | Levenberg–Marquardt (tight) | 0.35 | 0 | $3.1\times10^{-9}$ |
| airway | fixed | Dogleg (tight) | 0.55 | 0 | $5.3\times10^{-8}$ |
| airway | fixed | L-BFGS (tight) | 0.50 | 0 | $9.5\times10^{-8}$ |
| airway | ML | RapidGLM Newton | 0.98 | 198 | $6.0\times10^{-4}$ |
| airway | ML | RapidGLM Newton, plain `fit` | 1.07 | 198 | $6.0\times10^{-4}$ |
| airway | ML | BFGS (default) | 1.13 | 17,804 | $2.8\times10^{1}$ |
| airway | ML | L-BFGS (default) | 1.11 | 17,884 | $2.8\times10^{1}$ |
| airway | ML | BFGS (tight) | 1.75 | 6 | $3.3\times10^{-1}$ |
| airway | ML | L-BFGS (tight) | 1.96 | 8 | $3.3\times10^{-1}$ |
| simulation | fixed | RapidGLM Newton | 1.15 | 0 | $7.6\times10^{-11}$ |
| simulation | fixed | RapidGLM Newton, plain `fit` | 1.26 | 0 | $7.6\times10^{-11}$ |
| simulation | fixed | Levenberg–Marquardt (default) | 1.99 | 0 | $2.7\times10^{-3}$ |
| simulation | fixed | Dogleg (default) | 2.05 | 0 | $2.7\times10^{-3}$ |
| simulation | fixed | L-BFGS (default) | 3.12 | 362 | $1.4\times10^{1}$ |

| Data | Dispersion | Method | Seconds | Genes worse | Largest shortfall |
|---|---|---|---|---|---|
| simulation | fixed | Levenberg–Marquardt (tight) | 3.57 | 0 | $2.6 \times 10^{-9}$ |
| simulation | fixed | Dogleg (tight) | 3.70 | 0 | $2.7 \times 10^{-9}$ |
| simulation | fixed | L-BFGS (tight) | 4.02 | 0 | $6.4 \times 10^{-10}$ |
| simulation | ML | RapidGLM Newton | 2.46 | 0 | $1.3 \times 10^{-5}$ |
| simulation | ML | RapidGLM Newton, plain `fit` | 2.58 | 0 | $1.3 \times 10^{-5}$ |
| simulation | ML | BFGS (default) | 10.30 | 120 | $4.2 \times 10^{1}$ |
| simulation | ML | L-BFGS (default) | 10.29 | 120 | $4.2 \times 10^{1}$ |
| simulation | ML | BFGS (tight) | 12.94 | 0 | $8.5 \times 10^{-6}$ |
| simulation | ML | L-BFGS (tight) | 13.18 | 0 | $5.6 \times 10^{-5}$ |
| thyroid | fixed | RapidGLM Newton | 5.39 | 0 | $7.9 \times 10^{-8}$ |
| thyroid | fixed | RapidGLM Newton, plain `fit` | 5.91 | 0 | $7.9 \times 10^{-8}$ |
| thyroid | fixed | Levenberg–Marquardt (default) | 8.11 | 0 | $7.2 \times 10^{-4}$ |
| thyroid | fixed | Dogleg (default) | 8.32 | 0 | $1.1 \times 10^{-3}$ |
| thyroid | fixed | L-BFGS (default) | 12.23 | 23,163 | $2.5 \times 10^{3}$ |
| thyroid | fixed | Levenberg–Marquardt (tight) | 12.13 | 0 | $8.9 \times 10^{-9}$ |
| thyroid | fixed | Dogleg (tight) | 12.43 | 0 | $1.3 \times 10^{-8}$ |
| thyroid | fixed | L-BFGS (tight) | 24.87 | 3 | $4.0 \times 10^{-2}$ |
| thyroid | ML | RapidGLM Newton | 12.54 | 0 | $1.4 \times 10^{-5}$ |
| thyroid | ML | RapidGLM Newton, plain `fit` | 13.54 | 0 | $1.4 \times 10^{-5}$ |
| thyroid | ML | BFGS (default) | 37.79 | 21,364 | $8.4 \times 10^{3}$ |
| thyroid | ML | L-BFGS (default) | 37.75 | 21,475 | $8.4 \times 10^{3}$ |
| thyroid | ML | BFGS (tight) | 76.31 | 2 | $2.2 \times 10^{1}$ |
| thyroid | ML | L-BFGS (tight) | 80.45 | 3 | $3.2 \times 10^{-2}$ |

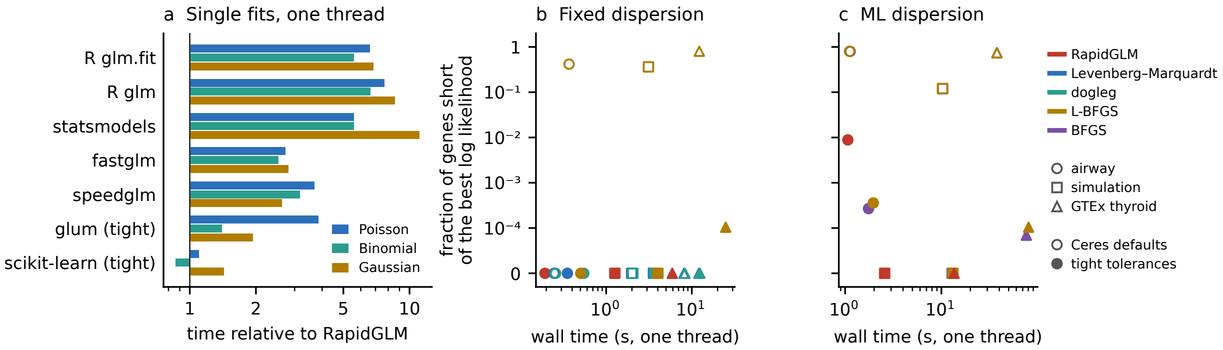


Figure S1: The engine against standard fitters and against Ceres Solver. (a) Median time of a single fit by standard GLM fitters, relative to RapidGLM's C++ engine, over the problems with at least 100,000 observations (Table S2). (b, c) RapidGLM and the methods of Ceres Solver on the same per-gene NB problems, at a fixed dispersion (b) and with the dispersion fitted by maximum likelihood (c): time on one thread against the fraction of genes whose log likelihood is more than $10^{-6}$ (relative) short of the best answer any method found (Table S4). Colour gives the method and shape the dataset. Ceres is shown at its default tolerances (open symbols) and at tight ones (filled); a fraction of zero is drawn at the foot of the axis. RapidGLM's time is that of a plain fit of each gene on its own. Its only shortfalls, on airway with a fitted dispersion, are genes whose $\theta$ reaches its bound of $10^{8}$ (at most $6 \times 10^{-4}$).

**Defects found and fixed.** The comparison exposed four defects in RapidGLM's joint dispersion fit. Each was fixed with a regression test, and the first three fixes were validated against the commit before them (`validation/JOINT_NEWTON_FIX.json`).

1. A moment start with no apparent overdispersion placed $\theta$ at its bound, where the likelihood is flat.

2. On profiles with an interior maximum, a dip and a lower Poisson asymptote, unit steps in $\theta$ overshot the maximum while the coefficients lagged.
3. Diverging coefficients made the information singular while $\theta$ still moved.
4. Interior local maxima below the Poisson limit were not compared with that limit.

### S3 The DESeq2 procedure

Table S5: Agreement with DESeq2 1.48.2 (discoveries at a false discovery rate of 0.1). sim3 and sim3b: simulations with three samples per group; sim8: eight per group. Receipt: `validation/BULK_DE.json`.

| Dataset | Discoveries (RapidGLM / DESeq2) | Jaccard | $p$-value Spearman |
|---|---|---|---|
| airway | 4,833 / 4,834 | 0.986 | 0.9998 |
| Kang | 635 / 635 | 1.000 | 1.0000 |
| sim3 | 239 / 239 | 0.983 | 1.0000 |
| sim3b | 201 / 201 | 1.000 | 1.0000 |
| sim8 | 253 / 253 | 1.000 | 0.9999 |

In 70,000 tests of a mock-comparison null (70 four-versus-four donor splits of the Kang panel), RapidGLM rejected 2.37% and DESeq2 2.39% at $p \leq 0.05$.

**Floored fits.** Genes with a fitted count below 0.5, or a coefficient without a finite estimate, are fitted as DESeq2 fits them (main text, Methods; Section S10), and Table S5 includes these fits. On the pseudobulk designs of the 20 perturbations of the Replogle pilot, the Jaccard indices with DESeq2 are 0.97–1.00 (`validation/REPLOGLE_PSEUDOBULK.json`). Before release 1.4.0, which added these fits, development runs on Perturb-seq pseudobulk designs gave 0.09–0.88.

**Cook's distances in large multi-factor designs.** Cook's distance measures how much one sample moves a gene's fitted coefficients. A gene with a large distance in a group of three to six samples is set aside (no $p$-value). In groups of seven or more samples, the outlying count is replaced and the gene refitted. The distances depend on a robust dispersion $\alpha^{\text{rob}}$, and on GTEx thyroid its estimator decides the agreement with DESeq2.

DESeq2's paper [5] states that Cook's distances use "a method-of-moments estimate $\alpha^{\text{rob}}$, using a robust estimator of variance". Its documentation describes this as a robust within-group variance, but specifies neither the estimator nor the groups.

- *Recovering the values.* Inverting DESeq2's Cook's distances, using its `cooks`, `mu` and hat-value assays, gives one $\alpha^{\text{rob}}$ per gene, consistent across samples to $10^{-12}$ (analysis of 2026-10-01, carried over in `validation/COOKS_THYROID.json` and described in `docs/BULK_DE.md`).
- *What the values show.* They have a floor of exactly 0.04. Above the floor, none of the moment estimators we tested reproduced them (all-sample or within-group; variance, MAD or trimmed variance; residuals about group or fitted means).

**The default: DESeq2's dispersion as implemented in PyDESeq2.** PyDESeq2 (MIT) [6] implements DESeq2's robust dispersion, and RapidGLM reimplements it with attribution (`legacy_cooks=true`, the default from release 1.2.0).

- *Variance.* In each design group of at least three samples, take the trimmed mean of the normalized counts, then the scaled trimmed mean of their squared deviations from it. Trim and scale are 1/3 and 2.04 below 3.5 samples, 1/4 and 1.86 below 23.5, and 1/8 and 1.51 otherwise. The variance $v$ is the maximum over groups; without such groups it is 1.51 times the 1/8-trimmed variance of all samples.

- *Dispersion.* $\alpha^{\text{rob}} = (v - \bar{\mu})/\bar{\mu}^2$, floored at 0.04.
- *Flagging.* A gene is not flagged when three or more samples have larger counts than the sample with the largest distance.

These dispersions equal the values recovered from DESeq2's outputs to $2 \times 10^{-12}$. On the 128-sample subset RapidGLM then sets aside 219 genes and DESeq2 224, 217 of them the same. Applied to DESeq2's own distances, RapidGLM's rule, with or without the larger-count exception, gives DESeq2's 224 genes exactly, so the 7 genes set aside by DESeq2 only and the 2 by RapidGLM only differ in their distances (`validation/COOKS_FLAGS.json`):

- *No distances* (2 genes, DESeq2 only). A design level has only zero counts, so the fit has a coefficient without a finite estimate, and RapidGLM computes no Cook's distances for such a fit.
- *Replaced counts* (7 genes, 5 of them DESeq2 only). Counts were replaced in a group of seven or more samples. DESeq2's reported distances are the same before and after the replacement, and its flags follow them; RapidGLM flags on the distances of the refitted gene.

Discoveries agree with DESeq2's with Jaccard 0.997 (128 samples) and 1.000 (647 samples), and at least 0.98 on every validation dataset. With flagging disabled in both tools the discoveries are identical on both thyroid sets. Receipts: `validation/COOKS_LEGACY.json`, `validation/COOKS_THYROID.json`, `validation/COOKS_FLAGS.json`.

**RapidGLM's own estimator.** With `legacy_cooks=false` RapidGLM uses the estimator that was its default before release 1.2.0, chosen on two-group simulations: squared MADs of normalized counts, pooled within design groups of at least three samples. It gives larger distances when there are many small groups. On the 128-sample subset it sets aside 753 genes against DESeq2's 224, and discoveries agree with Jaccard 0.58 (128 samples) and 0.90 (647 samples). Everything else agrees on that subset: size factors, dispersions, fold changes and $p$-values have median relative differences of $10^{-4}$ or less (`docs/BULK_DE.md`). With flagging disabled in both tools, discoveries still agree only with Jaccard 0.97 (128 samples) and 0.96 (647 samples), because the distances also choose the counts that are replaced: in 284 genes of the 128-sample subset, against 67 with the default. Receipt: `validation/COOKS_THYROID.json`.

### S4 edgeR's quasi-likelihood workflow

**Agreement.** On airway, the Kang panel and three simulations the discoveries at a false discovery rate of 0.1 equal those of edgeR 4.6.2, and $p$-values agree to within 0.006 in $\log_{10}$ (Table S6). On every dataset the kept genes, TMM factors (to $5.1 \times 10^{-15}$ relative) and average log CPM agree to rounding, and the common dispersion to within the tolerance of the search. Receipt: `validation/EDGER_QL.json` (`scripts/validate_edger.py`).

**Fits whose reported deviance is not that of their fitted values.** On the GTEx thyroid sets of 32, 128 and 647 samples, the check of edgeR's fits (Section S11) finds 22, 7 and 9 genes for which `glmFit`, for the full or the reduced design, reports a deviance that is not the deviance of its own fitted values, or flags the fit as failed. edgeR's $F$ statistics for those genes use the reported deviance. Without edgeR's implementation the reason cannot be checked. Table S6 counts discoveries over all the kept genes, these included. Over the other genes, with the Benjamini–Hochberg adjustment over those genes, the discoveries differ by at most 7 (7,480 against 7,477 for 647 samples).

**Times.** Table S7 gives the wall times of the workflow on the bulk datasets; the main text (Table 5) reports GTEx thyroid.

Table S6: Agreement with edgeR 4.6.2 (discoveries at a false discovery rate of 0.1). Both tools keep the same genes and find the same common dispersion to the digits shown. Prior d.f.: the prior degrees of freedom of the quasi-likelihood dispersions. Discoveries and the Jaccard index are over all the kept genes. $|\Delta \log_{10} p|$: the largest difference over the genes whose edgeR fits are consistent (see text). Receipt: `validation/EDGER_QL.json`.

| Dataset | Kept genes | Common dispersion | Prior d.f. (RapidGLM / edgeR) | Discoveries (RapidGLM / edgeR) | Jaccard | Largest $\lvert\Delta \log_{10} p\rvert$ |
|---|---|---|---|---|---|---|
| airway | 16,860 | 0.0113 | 5.39 / 5.39 | 6,568 / 6,568 | 1.0000 | 0.002 |
| Kang | 999 | 0.0233 | 9.15 / 9.15 | 695 / 695 | 1.0000 | 0.003 |
| sim3 | 3,142 | 0.1260 | 6.73 / 6.74 | 187 / 187 | 1.0000 | 0.003 |
| sim3b | 3,214 | 0.1381 | 5.54 / 5.54 | 165 / 165 | 1.0000 | 0.003 |
| sim8 | 3,196 | 0.0879 | 11.82 / 11.82 | 262 / 262 | 1.0000 | 0.006 |
| thyroid32 | 28,596 | 0.1185 | 4.10 / 4.10 | 1,364 / 1,387 | 0.9834 | 0.005 |
| thyroid128 | 28,681 | 0.1225 | 3.91 / 3.91 | 822 / 830 | 0.9904 | 0.018 |
| thyroid | 28,764 | 0.1364 | 3.88 / 3.88 | 7,481 / 7,479 | 0.9995 | 0.095 |

Table S7: The edgeR workflow: wall time in seconds on airway and on GTEx thyroid and its subsets. edgeR was run on one thread; RapidGLM ran from the command line. Receipt: `validation/APPLICATIONS_BULK.json`.

| Dataset | edgeR, $T$=1 | RapidGLM, $T$=1 | RapidGLM, $T$=8 |
|---|---|---|---|
| airway | 0.617 | 0.253 | 0.269 |
| thyroid32 | 5.06 | 1.14 | 0.525 |
| thyroid128 | 17.0 | 2.25 | 0.948 |
| thyroid | 77.7 | 8.17 | 3.34 |

### S5 cis-QTL mapping

**Data preparation.** Geuvadis gene counts (E-GEUV-1, GENCODE v12) were rounded to the nearest integer, as some are fractional assignments; library sizes are the sums of the rounded counts over all genes, before any filter. Transcription start sites are the Geuvadis gene coordinates. Expressed genes passed the GTEx v8 thresholds with RPKM in place of TPM: RPKM at least 0.1 and a count of at least 6 in at least 20% of samples. Normalized expression for the Gaussian models is TMM-normalized counts per million (TMM factors from edgeR), inverse normal transformed gene by gene with average ranks for ties. Genotypes are the Geuvadis release (1000 Genomes phase 1 with imputation [7, 8]), kept for the RNA-seq samples, biallelic SNVs only, called in every sample, with minor allele frequency at least 0.05. The genotype principal components come from SNVs pruned for linkage disequilibrium, and the number of expression principal components from a parallel analysis of 20 column-permuted replicates at the 95th percentile. Every model and tool used the same 42 covariates: five genotype principal components, 36 expression principal components and sex. The preparation scripts are in the repository (`scripts/eqtl_prep_expression.py`, `eqtl_prep_genotypes.sh`, `eqtl_make_covariates.py` and `eqtl_tmm_factors.R`).

**Reference runs.** FastQTL and tensorQTL took the normalized expression and the same covariates; FastQTL ran its adaptive permutations (1,000 to 10,000) and tensorQTL its fixed 10,000, each with its Beta approximation. FastQTL was later also run with a fixed 10,000 permutations (main text, Methods). jaxQTL's NB score test took the counts with offset the log library size and 1,000 permutations. Every tool used a window of 1 Mb around the transcription start site. Every tool's eGenes are the genes with Storey's $q < 0.05$ [9], computed with the `qvalue` package (version 2.40.0, defaults) from its gene-level $p$-values over the genes it tested validly.

**Agreement.** On chromosome 22, RapidGLM's Gaussian model found 267 eGenes against 267 for FastQTL's adaptive run and 266 for tensorQTL (Jaccard 1.000 and 0.996), with FastQTL's lead variant for 100% of genes. Its NB model found 168 against 170 for jaxQTL's NB score test (Jaccard 0.988; gene-level $p$-value

Spearman 0.9996; the same lead variant for 95% of genes). Table 5 of the main text gives the times. Receipt: `validation/EQTL_COMPARE.json` (`scripts/compare_eqtl.py`).

### S6 The SCEPTRE method

**Agreement under SCEPTRE's staging.** RapidGLM's `sceptre` reproduces the staging that SCEPTRE's outputs show (Section S12). Because the two tools draw different resamples, borderline pairs can differ, and SCEPTRE's results do not change with R's seed, so its own Monte Carlo spread cannot be measured by reseeding it. The relevant floor is therefore the agreement between RapidGLM runs with different seeds. These comparisons accept the skew-normal fit as SCEPTRE does (`fit_check=ks`). Receipt: `validation/SCEPTRE_STAGING.json`.

Table S8: Agreement with SCEPTRE 0.99.0 over ten seeds (discoveries at a false discovery rate of 0.1). Receipt: `validation/SCEPTRE_STAGING.json`.

| Panel | SCEPTRE | RapidGLM | Jaccard with SCEPTRE | Jaccard between seeds |
|---|---|---|---|---|
| Low-MOI example | 32 | 30–32 | 0.963 | 0.948 |
| High-MOI example | 11 | 10–16 | 0.863 | 0.854 |

On the full Gasperini screen the overlap with SCEPTRE's discoveries turns on RapidGLM's draws: over six seeds RapidGLM made 97–103 discoveries (SCEPTRE: 101) with Jaccard indices of 0.90–0.96, and two RapidGLM seeds overlapped with 0.90–0.97. On the public high-MOI example panel, where ten seeds are few, 200 seeds averaged 12.2 discoveries against SCEPTRE's 11. Receipt: `validation/PERTURBATION_SEEDS.json`.

**Requests without a list.** Pairwise quality control is decided before any cells are gathered (main text, Methods), so a request that names targets and responses costs little for the pairs that fail it. On a synthetic screen of 20,000 cells in which 5% of 12.4 million peak–target pairs pass (high MOI, 12 threads, with the sequential screen and early stopping), listing every pair took 117 s and 7.9 GB, and naming the targets and responses 91 s and 1.1 GB, with identical records for the tested pairs. Receipt: `validation/UNTESTABLE_PAIRS.json`.

**Streams and families in earlier releases.** Releases before 1.3.0 keyed a pair's stream by its place in the list and adjusted $p$-values over every listed pair. Both remain as options, and with them release 1.3.0 reproduced the tables of 1.2.0 byte for byte. A pair listed twice within a kind is an error; a pair listed under two kinds is one test with one $p$-value.

**Tail counts against the skew-normal fit.** For a pair that passes the first stage, the SCEPTRE method reads its $p$-value from a skew-normal curve fitted to the resampled statistics, which extrapolates below what the draws can count. With `fit_check=ks` RapidGLM accepts the fit on its Kolmogorov–Smirnov discrepancy alone; the agreement above was measured under that rule. To test the extrapolation we counted the tail directly on 100 targets against all 7,418 genes of the Gasperini screen. The 25 pairs that were a discovery in any of three runs, and one further pair, were each repeated 50 or 200 times with independent draws, 19,999 per repeat, giving 1 or 4 million draws per pair. For 7 pairs the number of draws beyond the observed statistic has probability below $10^{-3}$ under the fitted $p$-value (Table S9). All 7 have positive statistics: there the resampled statistics have a heavier right tail than the fitted curve.

With the default check (Section S12), 53 of 721,266 pairs drew the confirmation and 3 fits were rejected there. Discoveries fell from 20 to 14, for 4% more draws (Table S10); on the runs of Table S12 the default takes 16–46% longer. A rejected pair cannot report a two-sided $p$-value below $10^{-5}$, so a true effect whose resampled statistics are not skew-normal is not discovered at a genome-wide threshold. Receipts: `validation/SKEW_NORMAL_CHECK.json` (the direct counts, made on 2026-10-06 with an earlier release) and `validation/CRT_SCALING.json`.

Table S9: Pairs whose direct tail count contradicts the skew-normal $p$-value (counted with an earlier release). Counted: two-sided $p$-value from 1 or 4 million draws. Fitted: median over the repeats in which the KS check accepted the fit. Accepted: percentage of repeats that reported a skew-normal $p$-value under the KS check and under the strict check with 19,999 draws as the last budget. SCEPTRE: its $p$-value for the pair on the same panel (*, a discovery at a false discovery rate of 0.1).

| Response | Target | Statistic | Counted | Fitted | KS (%) | Strict (%) | SCEPTRE |
|---|---|---|---|---|---|---|---|
| ENSG00000084453 | chr3.4961_top_two | 9.11 | $1.0 \times 10^{-6}$ | $1.3 \times 10^{-10}$ | 40 | 0 | $6.2 \times 10^{-11}$* |
| ENSG00000185222 | chr9.4011_top_two | 7.22 | $2.3 \times 10^{-4}$ | $9.4 \times 10^{-8}$ | 84 | 0 | $8.0 \times 10^{-4}$ |
| ENSG00000151690 | chr22.1919_top_two | 6.82 | $1.0 \times 10^{-5}$ | $1.0 \times 10^{-7}$ | 100 | 2 | $1.7 \times 10^{-7}$* |
| ENSG00000152422 | chr5.980_top_two | 6.52 | $4.4 \times 10^{-5}$ | $5.4 \times 10^{-8}$ | 98 | 0 | $2.0 \times 10^{-7}$* |
| ENSG00000138772 | chr9.3066_top_two | 6.07 | $1.6 \times 10^{-5}$ | $2.1 \times 10^{-6}$ | 100 | 42 | $1.6 \times 10^{-6}$* |
| ENSG00000083123 | chr6.4971_second_two | 5.58 | $3.2 \times 10^{-5}$ | $7.6 \times 10^{-7}$ | 100 | 44 | $1.0 \times 10^{-6}$* |
| ENSG00000131724 | chrX.2160_top_two | 5.01 | $9.5 \times 10^{-4}$ | $5.2 \times 10^{-5}$ | 80 | 0 | $4.7 \times 10^{-5}$ |

**Scaling the CRT.** The batched draws, the ring of active pairs and the shared samplers (main text, Methods; Section S12) keep the conditional randomization test fast without changing any result. Table S10 gives wall times on panels of the Gasperini screen: every pair of $T$ targets and $G$ response genes, both drawn at random with a fixed seed (`scripts/prepare_crt_panels.py`). The whole screen, every target against every gene (28.0 million pairs), was measured separately (main text, Table 6). Receipt: `validation/CRT_SCALING.json`.

Table S10: CRT scaling on the Gasperini screen (207,324 cells, 16 threads on 16 cores) with SCEPTRE's acceptance of the skew-normal fit (`ks`) and with RapidGLM's defaults (strict check and confirmation draws). Tested: pairs with at least seven nonzero cells in each arm.

| | | Seconds | | Draws (millions) | | Discoveries | |
|---|---|---|---|---|---|---|---|
| **Targets × genes** | **Tested** | `ks` | default | `ks` | default | `ks` | default |
| 50 × 1,000 | 48,596 | 55 | 55 | 29.4 | 31.1 | 5 | 6 |
| 200 × 1,000 | 193,824 | 166 | 178 | 116.6 | 122.5 | 7 | 5 |
| 100 × 7,418 | 721,266 | 646 | 673 | 437.0 | 455.4 | 20 | 14 |

### S7 Applications by interface

Table S11: The DESeq2 procedure: every tool and interface (seconds at one and eight threads; discoveries and Jaccard against DESeq2 at one thread). DESeq2 with glmGamPoi was run on one worker only. In a separate comparison, made with an earlier release on another workstation (Intel Core i9-13900KF), these runs and the batch fit of Table S3 took the same time to within 0.6% with glmGamPoi's Bioconductor 3.21 release (1.20.0) in place of 1.25.1 (`validation/GLMGAMPOI_RELEASE.json`). Receipt: `validation/APPLICATIONS_BULK.json`.

| **Data** | **Tool** | $T$=1 | $T$=8 | **Discoveries** | **Jaccard** |
|---|---|---:|---:|---:|---:|
| airway | RapidGLM CLI | 0.888 | 0.267 | 4,833 | 0.986 |
| airway | RapidGLM Python | 0.874 | 0.28 | 4,833 | 0.986 |
| airway | RapidGLM PyDESeq2-style | 0.94 | 0.309 | 4,833 | 0.986 |
| airway | RapidGLM R | 0.847 | 0.244 | 4,833 | 0.986 |
| airway | PyDESeq2 | 41.3 | 9.47 | 4,690 | 0.931 |
| airway | PyDESeq2 + RapidGLM | 2.35 | 2.01 | 4,663 | 0.930 |
| airway | DESeq2 | 5.12 | 2.48 | 4,834 | 1.000 |
| airway | DESeq2 + glmGamPoi | 12.3 | – | 8,452 | 0.570 |
| thyroid32 | RapidGLM CLI | 4.08 | 0.736 | 1,620 | 0.987 |
| thyroid32 | RapidGLM Python | 3.92 | 0.727 | 1,620 | 0.987 |
| thyroid32 | RapidGLM PyDESeq2-style | 4.02 | 0.767 | 1,620 | 0.987 |
| thyroid32 | RapidGLM R | 4 | 0.673 | 1,620 | 0.987 |
| thyroid32 | PyDESeq2 | 79.4 | 15.5 | 1,637 | 0.985 |
| thyroid32 | PyDESeq2 + RapidGLM | 9.61 | 4.57 | 1,642 | 0.982 |
| thyroid32 | DESeq2 | 25.5 | 6.89 | 1,615 | 1.000 |
| thyroid32 | DESeq2 + glmGamPoi | 31.2 | – | 2,051 | 0.773 |
| thyroid128 | RapidGLM CLI | 8.28 | 1.55 | 790 | 0.997 |
| thyroid128 | RapidGLM Python | 7.97 | 1.37 | 790 | 0.997 |
| thyroid128 | RapidGLM PyDESeq2-style | 8.12 | 1.51 | 790 | 0.997 |
| thyroid128 | RapidGLM R | 8.27 | 1.51 | 790 | 0.997 |
| thyroid128 | PyDESeq2 | 87.6 | 17.1 | 792 | 0.997 |
| thyroid128 | PyDESeq2 + RapidGLM | 15.4 | 6 | 787 | 0.991 |
| thyroid128 | DESeq2 | 95.9 | 26.3 | 792 | 1.000 |
| thyroid128 | DESeq2 + glmGamPoi | 71.7 | – | 841 | 0.926 |
| thyroid | RapidGLM CLI | 29.4 | 5.29 | 7,178 | 1.000 |
| thyroid | RapidGLM Python | 28.9 | 4.88 | 7,178 | 1.000 |
| thyroid | RapidGLM PyDESeq2-style | 29.4 | 5.43 | 7,178 | 1.000 |
| thyroid | RapidGLM R | 30 | 5.54 | 7,178 | 1.000 |
| thyroid | PyDESeq2 | 127 | 24.4 | 7,177 | 1.000 |
| thyroid | PyDESeq2 + RapidGLM | 45.7 | 12.5 | 7,176 | 1.000 |
| thyroid | DESeq2 | 401 | 70.2 | 7,177 | 1.000 |
| thyroid | DESeq2 + glmGamPoi | 232 | – | 7,168 | 0.989 |

Table S12: The SCEPTRE method on the Gasperini screen: times and agreement with SCEPTRE 0.99.0 over the discovery pairs both tools tested. `ks`: SCEPTRE's acceptance of the fitted tail (`fit_check=ks`); the other RapidGLM rows use its defaults. Receipt: `validation/APPLICATIONS_PERTURBATION.json`.

| **Cells** | **Tool** | **Threads** | **Seconds** | **Discoveries** | **Jaccard** | **Spearman** |
|---|---|---|---|---|---|---|
| 50,000 cells | RapidGLM CLI | 1 | 260 | 24 | 0.846 | 0.986 |
| 50,000 cells | RapidGLM CLI | 8 | 36 | 24 | 0.846 | 0.986 |
| 50,000 cells | RapidGLM Python | 1 | 264 | 24 | 0.846 | 0.986 |
| 50,000 cells | RapidGLM Python | 8 | 36 | 24 | 0.846 | 0.986 |
| 50,000 cells | RapidGLM R | 1 | 300 | 24 | 0.846 | 0.986 |
| 50,000 cells | RapidGLM R | 8 | 43 | 24 | 0.846 | 0.986 |
| 50,000 cells | RapidGLM CLI, `ks` | 1 | 222 | 24 | 0.846 | 0.986 |
| 50,000 cells | RapidGLM CLI, `ks` | 8 | 31 | 24 | 0.846 | 0.986 |
| 50,000 cells | RapidGLM Python, `ks` | 1 | 223 | 24 | 0.846 | 0.986 |
| 50,000 cells | RapidGLM Python, `ks` | 8 | 30 | 24 | 0.846 | 0.986 |
| 50,000 cells | RapidGLM R, `ks` | 1 | 246 | 24 | 0.846 | 0.986 |
| 50,000 cells | RapidGLM R, `ks` | 8 | 36 | 24 | 0.846 | 0.986 |
| 50,000 cells | SCEPTRE | 1 | 1380 | 24 | 1.000 | 1.000 |
| 50,000 cells | SCEPTRE | 8 | 207 | 24 | 1.000 | 1.000 |
| 207,324 cells | RapidGLM CLI | 8 | 223 | 98 | 0.895 | 0.986 |
| 207,324 cells | RapidGLM Python | 8 | 208 | 98 | 0.895 | 0.986 |
| 207,324 cells | RapidGLM R | 8 | 238 | 98 | 0.895 | 0.986 |
| 207,324 cells | RapidGLM CLI, `ks` | 8 | 153 | 98 | 0.895 | 0.986 |
| 207,324 cells | RapidGLM Python, `ks` | 8 | 152 | 98 | 0.895 | 0.986 |
| 207,324 cells | RapidGLM R, `ks` | 8 | 182 | 98 | 0.895 | 0.986 |
| 207,324 cells | SCEPTRE | 8 | 1108 | 101 | 1.000 | 1.000 |

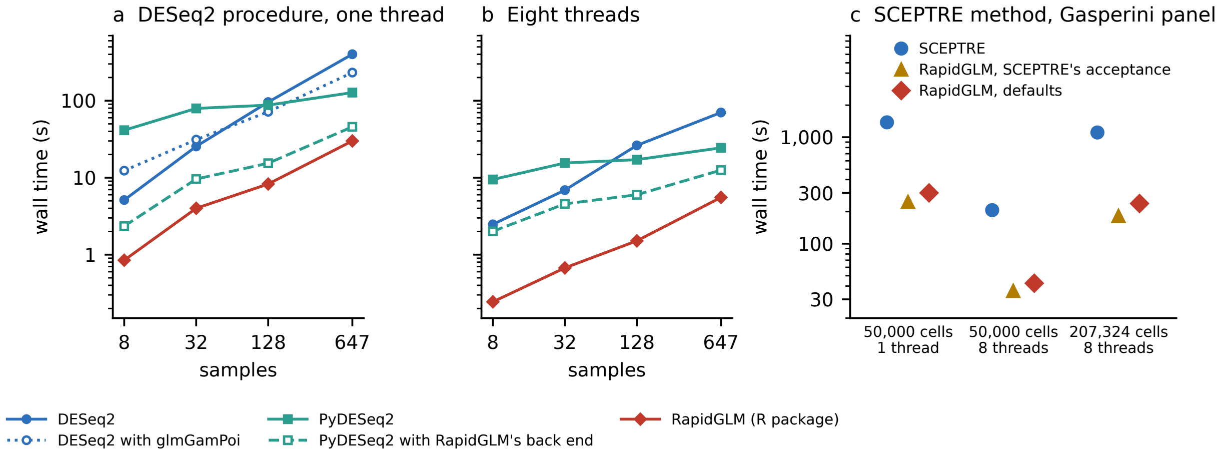


Figure S2: Times of the applications. (a, b) The DESeq2 procedure: wall time against the number of samples (airway, 8; GTEx thyroid, 32, 128 and 647) on one thread (a) and on eight (b), for DESeq2, DESeq2 with glmGamPoi (one worker only), PyDESeq2, PyDESeq2 with RapidGLM's back end, and RapidGLM's R package (Table S11). (c) The SCEPTRE method on the Gasperini screen: wall time of SCEPTRE and of RapidGLM's R package, with SCEPTRE's acceptance of the fitted tail (`fit_check=ks`) and with RapidGLM's defaults (Table S12).

### S8 Interfaces and reproducibility

- *C layer.* `librapidglm_c` has versioned option structures; a caller built against an older header passes a shorter structure and gets defaults for the rest.

- *Resampled results.* A pair's record is a function of the data, the options, the seed, the response's identifier (its ID, or its column when no IDs are given) and the target's name. It does not depend on the thread count, on the other pairs of the run, or on how the run is split into tiles or jobs; tests compare such runs record by record. A QTL permutation is likewise keyed by the seed and the gene's identifier.
- *Cross-interface tests.* The Python package's tables equal the command-line program's bit for bit. The R package's equal them for the bulk workflows and, from release 1.6.1, for cis-QTL mapping, and agree to $10^{-10}$ for the SCEPTRE method.
- *PyDESeq2 back end.* It is tested method by method against PyDESeq2's default inference on identical inputs.
- *Other languages.* Any language with a C foreign-function interface can call the library; as an example, `cargo test` runs the repository's Rust binding in continuous integration.
- *Dockerfile.* The Dockerfile builds the library, command-line program, C layer, Python package and R package and runs their test suites; continuous integration runs it on every push.

**Sections S9–S14** give in full what the Methods of the main text summarize: the constants, fallbacks and kernels of the engine and of each analysis, and the benchmark protocol. They describe RapidGLM 1.6.0. A number marked as a development measurement was made before the release.

### S9 The engine: fitting, shared work, kernels and interfaces

**Fitting.** Fisher scoring solves $(X'WX)\delta = X'r$, $r$ the score residuals, with $w_i = a_i(\partial\mu/\partial\eta)^2/V(\mu_i)$; for the NB at fixed $\theta$ the step uses the observed information, $w_i = a_i\theta\mu_i(\theta + y_i)/(\theta + \mu_i)^2$, which is positive for the log link. Step halving, as in `glm2` [10], requires a nondecreasing likelihood; on the concave count likelihoods Newton's method then converges reliably, and quadratically near the optimum. Count fits converge when the relative deviance change $|D_{\text{new}} - D_{\text{old}}|/(0.1 + |D_{\text{new}}|)$ and the relative coefficient change fall below the tolerance; after a step solved by Cholesky factorization, the score at the new estimate and that step's factor give a chord step and the deviance change it promises, and when both meet the criteria, and no working weight has moved by more than a quarter, the chord step is applied as a final refinement and the fit ends. Systems are solved by a Jacobi-scaled Cholesky factorization, computed in-house, while its reciprocal condition number is at least $10^{-8}$, and by Householder QR of $\sqrt{W}X$ otherwise. Aliased columns are detected by that QR with R's threshold. For unpenalized count fits, a candidate boundary direction is projected into the null space of the positive-count design; a certificate of zero change on positive counts and nonpositive change on zero counts identifies coefficients without a finite maximum, which are flagged, with not-a-number standard errors.

**Shared work.** Most of the time of a small fit goes to work other than the Newton steps themselves, and the engine shares or skips that work where the estimates do not need it. Fits of many responses to one design share $X'X$, its rank check, the grouping of identical rows and, for sparse designs such as those of factors, each row's nonzero entries; when the rows take exactly as many distinct values as there are columns (one-way and full-factorial layouts), a step needs only per-group sums, an idea from glmGamPoi [1]. The terms $\log\Gamma(y + \theta) - \log\Gamma(\theta) - \log y!$ and the polygamma sums of dispersion estimation, which depend on the response alone, are evaluated by Stirling's series in forms free of cancellation, one logarithm per large count, with a table for small counts. A score scan fits each response's null model once and tests many predictors against it, the pattern behind SCEPTRE's pairs and cis-QTL variants; the factorization of that fit gives the score statistics, and the fit of each gene in the bulk workflows gives its standard errors, leverages and Cook's distances without a refit.

**Dispersion.** Maximum-likelihood fits take joint Newton steps in $(\beta, \log\theta)$ from a Poisson pilot and the moment estimate of $\theta$; where the Schur complement shows the likelihood is not concave in $\log\theta$ the step in $\log\theta$ is a unit step up the slope, and step halving on the joint likelihood keeps every iterate an improvement. Cox–Reid fits, as for DESeq2's gene-wise dispersions [11, 12], maximize $\ell(\beta,\theta) - \frac{1}{2}\log\det(X'WX)$ with analytic first and second derivatives of the adjustment, alternating with coefficient fits, with Aitken extrapolation where the alternation converges slowly near the Poisson limit.

**Kernels and special functions.** The kernels of the quasi-likelihood and QTL workflows hold tiles of four rows by eight columns of a cross product in registers through all samples, summing each term in sample order; $X'WX$ sums only the tiles that reach the diagonal and copies the upper triangle from the lower; the leverages of all observations are squared norms of a triangular transform, the scores of many variants are computed at once, and linear predictors and their exponentials four at a time. After a whole Newton step the decrement at the new point is bounded by its value with the previous information times $e^{\Delta}$, $\Delta$ the largest change of a linear predictor, so a fit at a fixed dispersion can stop without another factorization at exactly the point where the full test would have stopped. Together these made edgeR's quasi-likelihood workflow 4.1 times faster on one thread, with the same discoveries (development measurement). The exponential evaluates $2^k\exp(r)$ with $r = x - k\log 2$ and the degree-13 Taylor polynomial of $\exp(r)$. The logarithm and $\log(1+x)$ take the exponent and mantissa from the bits and evaluate a degree-19 polynomial by Estrin's scheme with exact splits, and the rounding correction of $\log(1+x)$ is formed without dividing. Evaluated over arrays in vector lanes, `log` took 1.3 ns per element on one core of the benchmark host against 3.6 ns for the C library, and `exp` 0.8 ns against 4.6; one value at a time RapidGLM's `log` was slower than the C library's (5.5 ns), so the engine uses the array forms where whole arrays are evaluated, the exponentials of fitted means and edgeR's unit deviances, and the C library's functions and Boost's polygamma functions elsewhere. End to end the special functions changed workflow times by 0–5% (development measurement); their purpose is results that are identical across processors. Their largest measured errors are 0.67, 0.68 and 0.86 units in the last place for the logarithm, $\log(1+x)$ and the exponential, and 3.7 and 2.7 for RapidGLM's digamma and trigamma functions (`validation/SPECIAL_FUNCTIONS.json`). Each kernel has an AVX2 path with fused multiply–adds, chosen at run time, and a portable path, and the special functions also an Arm Advanced SIMD path; every alternative measured, and the reason for each choice, is listed in the repository's `METHODS.md`. Results are identical at any thread count. The special functions give the same bits on every path; on processors without AVX2 and fused multiply–adds the portable cross-product kernels round differently in the last bits. The portable paths are tested on x86-64 and, under emulation, on ARM64 (29 selected kernel, sampler, QTL and quasi-likelihood test cases; no native ARM64 timing is claimed).

**Interfaces.** The C layer, `librapidglm_c`, converts inputs and results only, so its outputs equal those of the C++ calls bit for bit. Designs are dense and row-major, and counts are compressed sparse column matrices, the layout of R's `dgCMatrix` and SciPy's `csc_matrix`. No C++ exception crosses the interface, and option structures begin with their size, so a caller built against an older header keeps working with a newer library. The Python package binds the C layer with `ctypes`, so its batch, scan and application calls need no compiler, and it accepts NumPy arrays, pandas data frames and SciPy sparse matrices. On it sit two forms for users of PyDESeq2 [6]: an inference back end, which replaces PyDESeq2's model fits, dispersion estimates and Wald tests and leaves the rest of PyDESeq2 unchanged, and a PyDESeq2-style interface, which takes the same kind of calls and runs the whole DESeq2 procedure in the engine. The R package compiles the C layer in behind a short `cpp11` shim and accepts `dgCMatrix` counts. For code that must stay in R or Python, drop-in fitters stand in for single-model calls: `rapidglm::glm_fit` as a `method` for `stats::glm`, `rapidglm::glm_nb` for `MASS::glm.nb`, and a statsmodels-compatible `GLM` class. The results do not depend on the interface: the Python package's tables equal the command-line program's bit for bit, and in the test suites the R package's equal them for the bulk workflows and, from release 1.6.1, for cis-QTL mapping, and agree to $10^{-10}$ for the SCEPTRE method (Section S8). The Rust binding declares the C functions by hand and is tested in continuous integration. It includes a port of PyDESeq2's workflow, made from PyDESeq2's MIT-licensed source, that runs the per-gene fits and dispersion estimates in RapidGLM. On GTEx thyroid on eight threads the port took 10.7 s, against 12.9 s in the same test for PyDESeq2 with the back end, and all 5,516 of its discoveries at a false discovery rate of 0.05 were among the back end's 5,517. The port and the back end differ in their fits of low counts (the back end floors the fitted counts, as PyDESeq2 does), in the solver of the dispersion trend and in the handling of Cook's distances, so this compares two workflows, not two timings of one computation.

#### S10 The DESeq2 procedure, step by step

`bulk_de` follows the procedure published for DESeq2 [5] and its documentation. DESeq2 1.48.2 was run only as a black box. Rules that the documentation leaves open were set from its outputs and are marked *calibrated*; rules of our own are marked as such. The robust dispersion of Cook's distances, the starting dispersion and the floored fits follow PyDESeq2's implementation (MIT) [6], which PyDESeq2 states follows DESeq2's code. The robust dispersions so computed equal those recovered from DESeq2's outputs to $2 \times 10^{-12}$. Counts $y_{ig}$ of gene $g$ in sample $i$ are fitted for $m$ samples against $p < m$ design columns, and coefficient $j$ is tested.

1. *Size factors* [13]. $s_i$ is the median, over genes without zeros, of $y_{ig}/G_g$, with $G_g$ the geometric mean of gene $g$. The `poscounts` variant takes $G_g$ as the $m$th root of the product of the positive counts, takes medians over positive counts only and scales the factors to geometric mean 1. Factors can also be supplied.
2. *Base means* $\bar{\mu}_g = m^{-1} \sum_i y_{ig}/s_i$. Genes with only zero counts are reported and skipped.
3. *Initial means.* When the design has as many distinct rows as columns, $\hat{\mu}_{ig}$ is $s_i$ times the mean normalized count of sample $i$'s design cell, floored at 0.5. Otherwise it comes from an NB fit at the starting dispersion, falling back to a Poisson fit and then to least squares; a fit with a fitted count below 0.5 is redone as a floored fit (step 8) and its unfloored means are used. The starting dispersion is the smaller of a rough value, $\max\{0, \sum_i (r_i^2 - f_i)/f_i^2/(m-p)\}$ from a least-squares regression of the normalized counts on the design (fitted values $f_i$ floored at 1, residuals $r_i$), and a moments value, $(v - \bar{x}\bar{\mu}_g)/\bar{\mu}_g^2$ with $v$ the variance of the normalized counts and $\bar{x}$ the mean of $1/s_i$; it is clamped to $[10^{-8}, \max(10, m)]$.
4. *Gene-wise dispersions* $\alpha_g$ maximize the Cox–Reid adjusted profile likelihood [11] at $\hat{\mu}$, by Newton's method in $\log \alpha$ from the starting dispersion. Genes whose starting dispersion is at the floor of $10^{-8}$ keep it, as in DESeq2's output (*calibrated*). When the size factors differ widely, as between pseudobulk samples of very different numbers of cells, the moments value is often negative, so such genes start and stay at the floor.
5. *Trend*, over the genes with $\alpha_g \geq 10^{-7}$ (*calibrated*). The parametric trend $\alpha_{\text{tr}} = a_0 + a_1/\bar{\mu}$ is a gamma GLM with identity link, refitted without the genes whose ratio $\alpha_g/\alpha_{\text{tr}}$ lies outside $[10^{-4}, 15]$ until the summed squared changes of the log coefficients fall below $10^{-6}$. If that fit fails, the trend is the running median of $\log \alpha_g$ over $\log \bar{\mu}$, with a window of 10% of the genes and at least 100 (our substitute for DESeq2's local regression). A constant trend, the mean of the $\alpha_g$, is also available.
6. *Prior.* $\log \alpha_g \sim N(\log \alpha_{\text{tr}}, \sigma^2)$ with $\sigma^2 = \max\{s_{\text{lr}}^2 - \psi_1((m-p)/2),\, 0.25\}$, where $s_{\text{lr}}$ is the median absolute deviation of $\log \alpha_g - \log \alpha_{\text{tr}}$ over the trend's genes and $\psi_1$ the trigamma function. For $m - p \leq 3$, $\sigma^2$ is the value on a grid of 1,000 points in $[0, 1]$ whose simulated density of $\log(\chi^2_{m-p}/(m-p)) + N(0, \sigma^2)$ is closest, in Kullback–Leibler divergence, to that of the observed residuals; the simulation uses 20,000 common draws and histograms of 100 bins over the observed range (our choice).
7. *Maximum a posteriori dispersions* maximize the adjusted likelihood plus the log prior, starting from the trend. Dispersion outliers, with $\log \alpha_g > \log \alpha_{\text{tr}} + 2 s_{\text{lr}}$, keep $\alpha_g$.
8. *Tests.* Each gene is fitted at its final dispersion with offsets $\log s_i$. The Wald statistic is $\beta_j/\text{SE}_j$, and the likelihood ratio test compares the fit with that of the design without column $j$ ($\chi_1^2$); the $\log_2$ fold change is $\beta_j/\log 2$. Where a fitted count of the maximum-likelihood fit falls below 0.5 (DESeq2's documented `minmu`), or a coefficient has no finite estimate, the gene is refitted as DESeq2 fits it, with fitted counts floored at 0.5, by PyDESeq2's iterations: iteratively reweighted least squares on $\mu_i = \max(s_i e^{x_i' \beta}, 0.5)$, with a ridge of $10^{-6}$ and coefficients held within $[-30, 30]$, until the deviance changes by at most $10^{-8}$ of itself plus 0.1. The Wald standard errors of these floored fits come from the sandwich $(H + 10^{-6} I)^{-1} H (H + 10^{-6} I)^{-1}$ with $H = X'WX$ at the floored means (*calibrated*: this reproduces DESeq2's standard errors, where PyDESeq2 uses the unfloored means), and the likelihood

ratio statistic is the difference of the two floored deviances. A group of zeros then has DESeq2's finite fold change.

9. *Cook's distances* [14] are $D_i = R_i^2 h_i/\{p(1-h_i)^2\}$, with $R_i = (y_i - \mu_i)/(\mu_i + \alpha^{\mathrm{rob}}\mu_i^2)^{1/2}$ and $h_i$ the leverages of the final fit; the cutoff is the 0.99 quantile of $F(p, m-p)$. The robust dispersion $\alpha^{\mathrm{rob}}$, which DESeq2's paper and documentation do not specify in reproducible detail, is computed as PyDESeq2 computes it (Section S3). In design cells of at least seven samples, counts whose distance exceeds the cutoff are replaced by $s_i$ times the 20% trimmed mean of the normalized counts, rounded down, and the gene is estimated again from step 3 with the trend and prior kept. Otherwise a gene whose largest distance over the samples in cells of three to six exceeds the cutoff gets no $p$-value, unless three or more samples have larger counts than the sample with that distance.
10. *Independent filtering* [15] on $\bar{\mu}_g$. At each of 50 quantile levels, from the fraction of genes with a base mean of zero to 0.95, the genes with $\bar{\mu}_g$ at or above the type-7 quantile pass (the inclusive rule is *calibrated*), and their Benjamini–Hochberg rejections [16] at the target false discovery rate are counted. A LOWESS curve [17] (span 0.2, *calibrated*; three robustness passes) is fitted to the rejections against the level, and the chosen level is the lowest whose rejections exceed the curve's maximum minus the residual standard deviation, $\{\sum \mathrm{residual}^2/49\}^{1/2}$. Adjusted $p$-values are computed over the genes that pass; the others have none.
11. *Fold-change shrinkage* (optional), with the paper's empirical normal prior, as `lfcShrink` computes it with `type="normal"` for a supplied model matrix. Each design column that varies across samples gets a zero-centred normal prior on its $\log_2$ coefficient with variance $(q/z_{0.975})^2$, where $q$ is the upper 0.05 quantile of the absolute maximum-likelihood coefficients, weighted by $1/(1/\bar{\mu}_g + \alpha_{\mathrm{tr}})$ as documented. The weights are scaled to sum to the number of genes, and the type-7 rule is applied to the sample with each value repeated by its weight (*calibrated*: this reproduces DESeq2's prior variances from its own estimates). Constant columns get variance $10^6$. The posterior mode is found by Fisher scoring with step halving; its standard error is the sandwich $(H+P)^{-1}H(H+P)^{-1}$, with $P$ the prior precision and $H$ from fitted counts floored at 0.5 (*calibrated*). Where counts were replaced, the replaced counts are used, consistent with the test (our choice; DESeq2's `lfcShrink` uses the original counts, a black-box observation).

#### S11 edgeR's quasi-likelihood workflow, step by step

`edger_ql` follows the default quasi-likelihood workflow of edgeR 4 [2, 18], the sequence `filterByExpr`, `normLibSizes`, `glmQLFit` and `glmQLFTest`. It was written from the papers, the documentation of edgeR and limma and black-box runs of edgeR 4.6.2, which settled the rules the documentation leaves open. Library sizes are $L_i$, with mean $\bar{L}$, for $n$ samples, $p$ design columns and $G$ genes.

1. *Filtering.* A gene is kept when its count per million reaches that of 10 reads in the median library in at least $k$ samples and its total count is at least 15. Here $k$ is the inverse of the design's largest leverage, $k_0$, or $10 + 0.7(k_0 - 10)$ when $k_0 > 10$. Library sizes are then recomputed over the kept genes.
2. *TMM factors* [19]. The reference sample is the one whose upper quartile of count shares is closest to the mean of those quartiles. For each sample, the genes with reads in it and in the reference are trimmed by 30% of their log ratios and 5% of their average log shares at each end (ties take average ranks), and the factor is the inverse-variance weighted mean log ratio. The factors are scaled to a product of 1.
3. *Average log CPM*, from a one-group NB fit at dispersion 0.05 of the counts plus $2L_i/\bar{L}$, with offsets $\log(L_i + 4L_i/\bar{L})$.
4. *The NB dispersion* $\alpha = 1/\theta$ maximizes the Cox–Reid adjusted profile likelihood summed over the $\lceil tG \rceil$ genes of highest average log CPM, $t = 0.02 + 0.98\{20/(G\sqrt{n-p})\}^{1/3}$ (1 when $G\sqrt{n-p} \le 20$). It is found in $\alpha^{1/4}$ on $[0, 4^{1/4}]$ by Brent's method, to a tolerance of $10^{-5}$.
5. *Quasi-dispersions.* Each gene is fitted at the working dispersion $\alpha/a$. Let $E_i$ and $V_i$ be the mean and variance of the NB unit deviance at mean $\mu_i/a$ and dispersion $\alpha$. Observation $i$'s unit deviance $d_i$ is divided by $V_i/(2E_i)$ and given $(1-h_i)\,2E_i^2/V_i$ degrees of freedom, with $h_i$ its leverage; the gene's $s^2$ is the sum of the first over the sum of the second, with $d$ degrees of freedom. The scale $a$ is the larger

of 1 and the 0.9 quantile of the fourth power of a LOWESS fit (span 0.5, three robustness passes) of $(s^2)^{1/4}$ on average log CPM, computed at $a = 1$ and again at that value, with every gene refitted each time. The moments $E_i$ and $V_i$ are exact (our choice): sums over the NB distribution, by quadrature for counts above 1,000, tabulated for each dispersion on 64 points per decade of the mean. edgeR uses an undocumented approximation of them that differs from the exact values by up to about $10^{-3}$ for dispersions up to 0.3, and by more at larger dispersions and means.

6. *Moderation* [3], with limma's weighting for unequal degrees of freedom. $s^2$ is floored at $10^{-12}$ of its median. The terms $\log s^2 - \psi(d/2) + \log(d/2)$, weighted by $1/\psi'(d/2)$, are smoothed against average log CPM by a weighted LOWESS (span $0.3{+}0.7(500/G)^{1/3}$, or 1 for $G \leq 500$; anchors at 1/200 of the range). The prior degrees of freedom $d_0$ maximize the likelihood of the scaled $F$ model, with the prior scale tied to the trend through $\psi(d_0/2) - \log(d_0/2)$. The search ends at 8,134.84478, the value edgeR 4.6.2 reports when the likelihood keeps increasing (*calibrated*). The posterior is $s^2_{\text{post}} = (d_0 s^2_{\text{prior}} + d\, s^2)/(d_0 + d)$.
7. *Tests.* $F = (D_{\text{reduced}} - D_{\text{full}})/(q\, s^2_{\text{post}})$ on $q$ and $d_0 + d$ degrees of freedom, with the deviances at $\alpha/a$; Benjamini–Hochberg adjustment over the kept genes.
8. *Fold changes*, from fits at $\alpha/a$ of the counts plus $0.125 L_i/\bar{L}$ with offsets $\log(L_i + 0.25 L_i/\bar{L})$.

Fits at a fixed dispersion use Newton's method with the observed information, warm-started across passes: whole steps while the Newton decrement (about the deviance still to gain) shrinks, then steps halved until the deviance does not increase, until the decrement is below $10^{-14}$ times 100 plus the gene's total count. After a whole step, convergence is confirmed without a new factorization when the decrement computed with the previous information, times $e^{\Delta}$ with $\Delta$ the largest change of a linear predictor, is below that threshold; otherwise the next step computes the decrement. A fit that has not converged in 60 steps is refitted by the general fitter. Leverages come from the Cholesky factor of $X'WX$, or from a Householder QR of $W^{1/2}X$ when fitted means near zero make that factor ill-conditioned.

**The check of edgeR's fits.** `scripts/reference_edger_fits.R` fits every kept gene with edgeR's `glmFit`, for the full and the reduced design, at the working dispersion that `glmQLFit` uses, and compares the deviance `glmFit` reports with the deviance of its fitted values (edgeR's `nbinomUnitDeviance`, summed). It also records the iteration counts and failure flags that `glmFit` returns. The check uses only edgeR's public functions.

### S12 The SCEPTRE method in full

`sceptre` follows the SCEPTRE method as its papers [20, 21] and online documentation describe it. SCEPTRE 0.99.0 was run only as a black box.

**Inputs and quality control.** The inputs are sparse responses (genes or peaks), fixed binary guide calls with each guide's target, a design with offsets, and pairs of three kinds: discovery pairs, negative controls and positive controls. Guides are called upstream. Cells outside the 1st to 99th percentiles of total count or of detected features, or with a mitochondrial fraction above 0.2, are removed, and at low MOI so are cells with no guide or more than one. A target's guides are pooled by default; testing each guide, with or without a Bonferroni combination, is optional. A pair passes pairwise quality control, and is tested, when at least seven treated cells and seven control cells have a nonzero count of its response. This is decided first, from the target's cells and the response's nonzero cells alone, so a pair that fails costs nothing more. A request may list pairs or name targets and responses; in the second case one pass over each response's nonzero cells decides pairwise quality control for every target, the pairs that pass are tested with the records they would have in a list, and the others are only counted.

**Null model and statistic.** Controls are, at high MOI, the cells without the target's guides and, at low MOI, the cells with non-targeting guides. Following SCEPTRE's documented response model, the null model is a Poisson regression of the response on the design with its offsets, after which the NB dispersion is estimated by maximum likelihood with the fitted means held fixed. With complement controls one null fit per response serves all its pairs; with non-targeting controls the null is fitted for each pair on its treated and control cells (our choice of subset). That per-pair fit starts by default from the response's reference: the null fitted once on all retained cells. At the reference, each cell contributes additive totals (the Poisson score and information, and the terms of the conditional dispersion score and curvature with their derivative in the coefficients); the non-targeting cells' totals are summed once per response, and a pair adds those of its target's cells in a fixed order. One Fisher-scoring step in the coefficients and one Newton step in $\log\theta$ from the pair's totals give a one-step estimator [22], which differs from the pair's fit by the share of other targets' cells in the reference. It is therefore used only as a warm start [23]: Newton passes over the pair's cells, with step halving, refine it to the full fit's convergence criteria, and $\theta$ is then re-estimated at the converged means. A pair whose refinement does not converge within 30 passes or meets a singular information, and every pair of a response whose reference has a boundary, is fitted from scratch. A pair's totals depend only on its set of cells, so permuting labels within the pair leaves its null unchanged, as for the fit from scratch. With score residuals $r_i = (y_i - \mu_i)/(1 + \mu_i/\theta)$ and weights $w_i = \mu_i/(1 + \mu_i/\theta)$, the statistic for treatment indicator $g$ is $z = g'r/I^{1/2}$ with $I = g'Wg - (X'Wg)'(X'WX)^{-1}X'Wg$, the score of SCEPTRE's documentation (the statistic $T$ of the main text).

**Resampling.** At high MOI the conditional randomization test [24] redraws every cell's treatment indicator independently, with its probability $p_i$ under a logistic regression of the indicator on the design. By default each cell is given an interval of length $-\log(1 - p_i)$, the intervals laid end to end in cell order, and the cells whose intervals hold at least one point of a unit-rate Poisson process, generated by adding standard exponential gaps, are treated. An interval holds no point with probability $1 - p_i$ and disjoint intervals are independent, so the labels are independent Bernoulli($p_i$), and the expected number of points, $\sum_i -\log(1-p_i)$, is about the expected number of treated cells. The gaps come from a 256-layer ziggurat [25] fed by ARS with ten rounds, the AES-based counter generator of Salmon et al. [26], keyed by the first two words of the draw's Philox stream, so a draw is a function of the seed, the test and the draw alone. On processors without AES instructions the cells are instead grouped by levels $q = 2^{-b}$ with $q/2 < p_i \le q$, and the candidates of a Bernoulli($q$) process are visited through geometric gaps and each accepted with probability $p_i/q$ [27]. Both samplers draw the same law, from different streams, so CRT $p$-values depend on the kind of processor within Monte Carlo error; the run record states the sampler used. Either way a draw costs time in proportion to the expected number of treated cells, not the number of cells. In RapidGLM 1.6.0 the null draws of 100 targets against all 7,418 genes of the Gasperini screen took 947 s with the Poisson-process sampler against 1,273 s with thinning (1.34×; 1.33× on 50 targets against 1,000 genes; Section S14). At low MOI the labels are permuted among the treated and control cells. Each resampled indicator is scored against the same null fit. Empirical $p$-values count ties, $(1 + \#\{\text{draws as extreme}\})/(B + 1)$ [28], and two-sided values are twice the smaller one-sided tail, capped at 1.

**Staging.** Budgets are cumulative: 499, 4,999 and 19,999 draws by default. The first budget always ends with its empirical $p$-value, which is final unless it is at most 0.02. Only then are the later budgets drawn, and a skew-normal distribution [29], fitted to the resampled statistics by moments, gives the $p$-value. SCEPTRE's documentation does not specify this staging; Barry et al. [21] give a first stage of 500 resamples and a threshold of about 0.01 as examples. We inferred the budget of 499 and the threshold of 0.02 from SCEPTRE 0.99.0's outputs: its first-stage $p$-values are multiples of 2/500, the smallest at that stage was 0.024, and 2.1% of 1,500 null pairs went on to the second stage. Under `fit_check=ks`, used for every comparison with SCEPTRE, a fit with admissible moments is accepted when its Kolmogorov–Smirnov discrepancy from the draws is at most $1.36/\sqrt{B}$; otherwise the next budget is drawn, and after the last the empirical $p$-value is reported. This acceptance rule, too, was inferred from SCEPTRE's outputs; its documentation names no acceptance test. Optionally (`sequential_screen`), the first budget ends as soon as its outcome is certain: every 32 draws, once the number of draws at least as extreme as the statistic has reached the smallest number at which the empirical $p$-value of the whole budget exceeds the threshold. A pair that goes on has drawn the whole budget, so its result is unchanged. A pair that ends early reports the sequential $p$-value of Besag and Clifford [30], that number divided by the number of draws made (for a two-sided test, twice the value of the side that reached it last, at most 1), raised to at least the smallest $p$-value at which a whole first budget ends. The later budgets can end early in the same way (`early_stop_p`). Neither option is a default.

**The default acceptance.** With `fit_check=strict`, RapidGLM's default, a fit must also be sound in the tail on the statistic's side. Each of the following comparisons is made at the 1% level: the draws at least as extreme as the statistic are no more than a Poisson count with the fit's expectation allows; the fit expects at least 1/100 of a draw beyond the most extreme draw; the draws beyond the fitted quantiles with 1, 4, 16, ... expected draws beyond them, down to a tail of 1/64, are within Poisson limits; and the excess kurtosis of the draws exceeds the fit's by no more than $3(24/B)^{1/2}$. A $p$-value that the budget cannot check, one for which the fit expects fewer than half a draw beyond the statistic, goes on to the next budget; after the last it is confirmed on 199,999 draws in all, and the fit made on those draws is reported if it is sound and the empirical $p$-value otherwise. A pair whose fit is rejected reports the empirical $p$-value of its draws.

**Random streams and multiplicity.** Each pair has its own Philox stream [26], and draw $k$ uses the stream's draw $k$. The stream is keyed by the run's seed, the response (the 64-bit FNV-1a hash of its identifier, or its column when no identifiers are given) and the target (the hash of its name, or of the guide's for a per-guide test). A pair's record is therefore a function of the data, the options, the seed, the response and the target: it does not depend on the thread count, on which other pairs are listed or in what order, or on how a run is divided into tiles or jobs. Optionally (`reuse_draws`, at high MOI), the streams are keyed by the target instead of the pair, so the draws of a target are shared by all its pairs, as in SCEPTRE's resampling scheme (Table 6 of the main text); the errors of a target's $p$-values are then correlated. Adjusted $p$-values are Benjamini–Hochberg values within each kind of pair, over the pairs that pass pairwise quality control. SCEPTRE's documentation does not name the family; its outputs show this one: on the Gasperini panel its discoveries are those of the adjustment over the pairs that pass its pairwise quality control, not over all listed pairs.

**Scaling.** At the scale of whole screens the conditional randomization test dominates the time, and its limit is scattered memory reads. Draws are generated and scored 256 at a time, each probability level's cells read once per batch, with every draw keeping its own stream and consuming it in the same order as alone. The blocks of a Philox stream are computed eight at a time with AVX2 instructions where the processor has them. All threads draw chunks of 32 draws from a small ring of active pairs, response by response, so that the response's per-cell records and the pairs' samplers are shared in cache, and idle threads fit the next responses' null models. With complement controls, a target's sampler is built once for all its pairs. By default in that setting (batch scoring), each thread instead takes one response and scores all its pairs together, stage by stage: every active pair contributes its next draws, and all of them are scored in one pass over the response's per-cell records (residual, weight and weighted design row) as a sparse-by-dense product, in blocks of 32,768 cells so that a block's records stay in cache while every draw adds those of its cells. Each draw still adds its cells in ascending order, so statistics, $p$-values and draws are bit for bit those of scoring pair by pair. With RapidGLM 1.6.0 on 16 threads, 100 targets against all genes of the Gasperini screen (721,266 pairs) took 673 s with batch scoring against 1,041 s without it (1.55×; 455 million draws), and 646 against 1,013 s with SCEPTRE's acceptance (1.57×). Batch scoring holds up to 8,192 draws of each of a response's pairs at once, so its memory grows with the number of targets in a tile: with the automatic tiling, which put all 3,837 targets of the Gasperini test set in every tile, the default run on the whole test set exceeded a 128 GiB instance, and with 64 targets per tile it ran in 33.9 GiB and gave the same table. With shared draws, each group of about 2,048 stored first-stage draws is also kept in cell-major form, and a tiled run keeps each response's inverse information and each target's propensity model across tiles. Because draw $k$ of a pair always uses its own key and is written to slot $k$, none of this changes any result; the test suite compares every draw and statistic with a plain, unbatched implementation.

#### S13 cis-QTL mapping: scores, permutations and their kernels

`qtl` follows the design of jaxQTL (MIT) [31], whose permutation scheme and Beta approximation in turn follow FastQTL [32]. Its inputs are phenotypes anchored at their transcription start sites (or gene bodies), PLINK genotypes, covariates and, for counts, library sizes.

- *Cis variants.* A gene's cis variants are those on its chromosome within a window, 1 Mb by default, on either side of its anchor. A missing dosage is replaced by the variant's mean dosage.

- *Null model.* Each gene is fitted against the covariates alone: a Gaussian model for normalized expression, or a Poisson or NB model (log link, offset the log library size, NB dispersion by maximum likelihood) for counts. A gene whose null fit fails, or has a coefficient without a finite estimate, is reported and not tested.
- *Scores.* Every cis variant is tested by its efficient score statistic against the null fit (the statistic $T$ of the main text, Methods, Engine; written $z$ here). With $W$ the working weights, $r$ the score residuals, $LL' = X'WX$ and $G$ the variants' dosages, the scores of all variants are $u = G'r - (L^{-1}X'WG)'(L^{-1}X'r)$ with information $I = \text{diag}(G'WG) - |L^{-1}X'WG|^2$ (squared column norms) and $z = u/(I\phi)^{1/2}$; the sums over samples run in one pass of the cross-product kernel. The lead variant has the largest $|z|$.
- *Permutations.* Each of $B$ permutations (1,000 by default) shuffles the response and its offset together against the covariates and genotypes, refits the null and keeps the largest $|z|$ over the gene's variants. Permutation $b$ of a gene is drawn from a Philox stream keyed by the seed and the gene's identifier, so results do not depend on the thread count or on the other genes of the run. The generator values of a shuffle are computed eight Philox blocks at a time, and each bounded integer of its Fisher–Yates swaps is taken from them without a division [33], so every shuffle is that of the generator drawn value by value. Genotypes are kept as the two-bit codes of the PLINK file and decoded gene by gene, and each thread keeps its buffers from gene to gene. For a Gaussian null without weights the working weights are the same in every refit, so the part of the scores that depends on them alone ($L$, $L^{-1}X'WG$ and $I$) is computed once per gene; each refit is then a least-squares residual, and the permutations are scored in batches of 512 by one cross product of the dosages with the batch of residuals. Unless double precision is requested, that product is computed in single precision for Gaussian nulls without prior weights, offsets, ridge or robust covariance: each permuted response is residualized and scaled to unit residual variance in double precision, the products are summed in single precision within panels of 128 samples (six-by-sixteen register tiles with fused multiply–adds, or a portable path) and in double precision across panels, and each permutation's maximum is taken inside the tiles. Permutation maxima within $10^{-5}$ (relative) below the observed statistic count as extreme ($10^{-9}$ in double precision). For Poisson and NB nulls each permutation's null is refitted. Eight permutations are scored together: their weighted designs, side by side, form one blocked product with the gene's genotypes, packed once per gene, and the sums still run over the samples in order, so each permutation's statistics are bit for bit those of scoring it alone. The same eight nulls are refitted in lockstep: each follows the single-fit algorithm on its own values (least-squares start, Poisson iterations with step halving, then for NB the moment dispersion and joint Newton steps), with the sums over samples of all eight taken as dense products; a fit that leaves the regular path, for example at a possible boundary or an ill-conditioned step, is refitted alone. Only the order of the sums differs from separate refits, which remain available.
- *Gene-level p-value.* The permutation $p$-value is $(1 + \#\{b : T_b \geq |z|\})/(B + 1)$ [28], with $T_b$ the permutation maxima. The Beta approximation follows jaxQTL: the maxima are referred to a noncentral $\chi_1^2$ distribution whose noncentrality makes the moment estimate of the first Beta shape equal to 1 (one effective test), and Beta$(k, n)$ is fitted to the resulting $p$-values by maximum likelihood (Newton steps with step halving, from the moment estimates). The gene-level $p$-value of the observed statistic is the Beta distribution function at that noncentral $\chi_1^2$ tail probability of $z^2$.

### S14 Benchmark protocol

**The benchmark protocol in full.** The version described is RapidGLM 1.6.0 (tag v1.6.0, commit 9c22070; measured at development commit 4505ef6, whose library source is identical), and every benchmark except the all-pairs Gasperini runs, the runs on the Bridges-2 supercomputer, the steps of Table 2 of the main text and the comparison with FastQTL in Table 5 was measured with it in one run of 24 stages: the engine (accuracy, single fits, batch fits and the comparison with Ceres Solver), the four applications against their reference tools, the analyses at scale, the special functions, functional tests on every interface, and each optimization new in 1.6.0 against the same build with that optimization switched off. Timings are wall-clock seconds on one AMD Ryzen Threadripper PRO 5965WX workstation (Zen 3; 24 cores, 48 logical CPUs, 125 GiB of RAM), under an exclusive allocation of fixed cores. Every timed command ran one thread per physical core (one thread on one core, eight on eight, twelve on twelve and sixteen on sixteen) with the cores' second hardware threads idle; drivers and untimed checks ran on six other cores, and from the bulk application stage on a resident service was confined to the remaining ones. Each timed command started only after two ten-second samples of a quiet host, the CPU frequency policy (governor, boost and simultaneous multithreading) was checked to be unchanged throughout, and a command during which other activity took more than 5% of its cores or their second threads was rejected and repeated. BLAS ran one thread per worker. The full Replogle analyses ran 16 single-thread workers, one per core, each taking the next perturbation from a shared queue, and the cis-QTL comparisons used twelve cores. The all-pairs Gasperini runs (RapidGLM 1.4.1, 1.5.0 and 1.6.0) each used one AWS m8id.8xlarge instance (16 cores of an Intel Xeon 6975P-C with simultaneous multithreading off, 128 GiB of RAM and local NVMe storage), and are compared only with each other. The 1.6.0 default run was repeated twice, once with batch scoring switched off and once with 64 targets per tile, each on its own instance of the same type, with the same image, binary, inputs and seed. The steps of Table 2 of the main text were timed after the release, with the released 1.6.0 library, whose source is that of the measured commit, by `scripts/benchmark_crt_steps.py` (`docs/CRT_STEPS.md`) on the same workstation: 16 threads on sixteen cores under an exclusive allocation, without the quiet-host gate. The program times only the null draws, in six forms that each change one thing from the one before: the draw code of the first implementation, which the benchmark carries because the library no longer contains it; the released per-draw code, still one draw at a time; draws generated 256 at a time and scored in blocks of cells; the pair pipeline, with one sampler per target; the same in the native build, where the forms before it are built with portable code only; and the native build with the Poisson-process sampler. Step 1 of the table is the second and third forms together: on the larger panel the released per-draw code (its sampler and generator, and one packed record of a cell's values in place of three arrays) made the draws 1.50 times faster than the first implementation's, and batching a further 1.14 times. Before step 3 the generator computed two Philox blocks per call; the portable build compared here computes eight, in scalar code. For the smaller panel each form ran once untimed and three times timed, in alternating order, and the median is reported; for the larger panel each ran once. The three timed runs of a form differed by at most 1.3% of their median. The five forms that share the thinning sampler use the same random numbers and made the same draws (31 million on the smaller panel and 457 million on the larger), and they wrote the same table, byte for byte, on each panel, and the pipeline's tables were also compared, pair by pair and bit for bit, with those of `sceptre()` on the same inputs. For every command the program recorded the processor time that other processes used on the timed cores and on their second hardware threads; the latter was at most 1.4% of those threads' capacity during a command. The comparison with FastQTL at a fixed 10,000 permutations (Table 5 of the main text) was also run after the release, as a separate job on the same workstation with the binaries and inputs of the 24-stage run. Its three commands, RapidGLM, FastQTL and RapidGLM again, each ran on the same twelve cores (6–8 and 10–18), chosen because they and their second hardware threads were idle, and each passed the quiet-host gate and the 5% rule at its first attempt. FastQTL ran as twelve processes, one per core (`--permute 10000 --chunk` *k* `12`), and its time is that of the whole command; its chunks took 260 to 634 s, and 5,621 s of processor time in all, against RapidGLM's 422 s. RapidGLM took 35.6 s before FastQTL and 35.3 s after it, and its results agreed with FastQTL's exactly as its results of the 24-stage run did. Table 5 gives the slower of RapidGLM's two times. In the 24-stage run FastQTL used its adaptive scheme, which stops a gene's permutations early, after between 1,000 and 10,000, and took 528 s. The comparison with SCEPTRE at scale ran on the Bridges-2 supercomputer at the Pittsburgh Supercomputing Center [34], on nodes with two 64-core AMD EPYC 7742 processors (simultaneous multithreading off): RM-512 nodes with 512 GB for SCEPTRE and for RapidGLM's 128-thread runs with independent draws, and RM nodes with 256 GB for RapidGLM's other runs and for SCEPTRE on fewer workers. Each run had an exclusive node, and RapidGLM's times for the same runs on the two node types differed by at most 4%. RapidGLM 1.6.0 was built on a node from the source of the cloud runs and passed its tests there. The pair sets were the benchmark panel, 100 targets against every gene with the panel's controls, and every discovery pair of the test set with the panel's controls, all with the same cells, covariates and seed; SCEPTRE's calibration check sampled 5,000 negative pairs from each. Wall time is from GNU time, and memory is the peak of the summed proportional set size of the process tree, which counts the memory that SCEPTRE's forked workers share once. SCEPTRE's time on a server is an extrapolation, not a measurement: its time on 100 targets at the number of workers the server's memory holds, interpolated between 4, 8, 16 and 32 workers, times the ratio of its whole-screen time to its 100-target time on 32 workers. The reference tools ran only through their public interfaces: DESeq2 1.48.2 (Bioconductor 3.21), PyDESeq2 0.5.2, edgeR 4.6.2, glmGamPoi 1.25.1,

SCEPTRE 0.99.0 (GitHub development version, commit 21f9ea0), FastQTL (GTEx fork, commit 59bcdc0), tensorQTL 1.0.10, jaxQTL 0.3.1, R 4.5.1 (`glm`, `MASS::glm.nb`), fastglm 0.1.2, speedglm 0.3-5, statsmodels 0.14.4, scikit-learn 1.7.2, glum 3.4.1 and Ceres Solver 2.2. To separate the solver from the surrounding code, the comparison with Ceres Solver [4] fitted the same per-gene NB problems with each method on airway, a simulation and GTEx thyroid, and scored every answer by one numerically stable log likelihood; it also exposed four defects in RapidGLM's joint dispersion fit, fixed before these measurements (Section S2). On more than one thread, DESeq2 ran through BiocParallel and SCEPTRE with its `parallel` option, as their documentation describes.

Each application ran at one and eight threads where the tool supports them, with the tools' runs repeated in rotation, and the minimum wall time is reported; the receipts record the number of repeats (three for the smaller bulk datasets, two for the 647-sample thyroid set and one for the perturbation runs). For Table 5 of the main text, RapidGLM ran through its R package for the DESeq2 procedure and the Gasperini panel, and from the command line for the edgeR workflow, the Replogle pilot and cis-QTL mapping; its SCEPTRE runs used SCEPTRE's acceptance of the fitted tail (`fit_check=ks`). The engine's fits were timed on data in memory, on one core, with each tool's default settings, as the minimum over up to seven repeats. Where a reference tool was timed on a subset of the responses, as statsmodels on 200 of the 1,000 responses of the batch benchmark, its time is scaled in proportion to the number of responses. In the analyses at scale the reference tools ran on the whole analysis. For the interfaces (Table 4 of the main text), the command-line program's time is that of the whole process, including reading the input files, and the Python and R packages' times are those of the calls on data already in memory; DESeq2's is that of `DESeq()` and `results()`, and PyDESeq2's, with and without the back end, that of building the data set, `deseq2()` and the summary of the test. Discoveries are compared by the Jaccard index at a false discovery rate of 0.1 (eGenes at $q < 0.05$), over the genes or pairs both tools tested, and $p$-values by Spearman correlation. Every number comes from a validation receipt in the repository that records the version and commit measured.
